# A mosaic of genomic architectures underpins parasitism loss in a jawless vertebrate

**DOI:** 10.64898/2026.05.11.724254

**Authors:** Arne Jacobs, Nolwenn Decanter, Ole K. Tørresen, Benedicte Garmann-Aarhus, Maria Capstick, Quentin Rougemont, Frédéric Guillaume, Romane Normand, Julien Tremblay, Jean-Pierre Destouches, Anne-Laure Besnard, Ahmed Souissi, Gilles Lassalle, Solenn Stoeckel, Eric J. Petit, Siv NK Hoff, Daniel J. Park, Bernard J. Pope, Sissel Jentoft, Leif Asbjørn Vøllestad, Kjetill S. Jakobsen, Guillaume Evanno

## Abstract

Lampreys are the only ancestrally parasitic vertebrate lineage, yet parasitism has been repeatedly lost alongside a suite of life-history changes, such as loss of migration and juvenile feeding and accelerated maturation. Combining whole-genome resequencing, haplotype-resolved assemblies, hybrid-zone genotyping, multi-tissue transcriptomics, and sperm phenotyping, we map this life-history syndrome in European *Lampetra* to six chromosomes spanning a mosaic of genomic architectures: a ∼20 Mb low-recombination region on chromosome 1 lacking chromosomal rearrangements within *Lampetra* but involving inter-specific rearrangements across deep lamprey lineages; a translocated inversion with ecotype-dependent sperm-velocity effects; and ecotype-divergent deletions overlapping genes crucial for nervous system (*CNTNAP2*) and reproductive development (*FSHR*). However, this genomic basis is not shared with a convergent sister lineage, pointing to independent routes to a recurring life-history transition in lampreys.

## Introduction

Life-history variation within species has often been linked to chromosomal rearrangements that differ in orientation between divergent populations and suppress recombination in heterozygotes. Examples include large chromosomal inversions impacting local adaptation in seaweed flies (*1*) and Atlantic silverides (*2*), life-history in monkeyflowers (*3*), or migratory strategies in cod (*4*, *5*) and rainbow trout (*6*), or reproductive strategies in ruff (*7*). While most studies have focused on simple structural variants, such as inversions, more complex intraspecific rearrangements can also facilitate adaptation under gene flow, such as inverted-translocations in Timema stick insects (*8*), or chromosomal fusions and fissions in butterflies (*9*), threespined sticklebacks (*10*), and codfishes (*11*). Yet, recombination might be reduced without polymorphic rearrangements between life-history forms, e.g. through historical interspecific chromosomal fusions (*10*, *12*) or recombination deserts on sex-chromosomes (*13*). The underlying genomic mechanisms are not well understood, though, and empirical studies linking low recombination regions to life-history evolution in the absence of polymorphic structural variation have been largely missing - despite the potential importance for adaptation.

Parasitic feeding is ubiquitous across the tree of life, yet comparatively rare in vertebrates, with famous examples including vampire bats (*14*), vampire finches (*15*) and lampreys (*16*). In contrast to other vertebrates, the ancestor of all extant lampreys was likely parasitic (*16*, *17*). However, lampreys have repeatedly lost their parasitic lifestyle in many independent lineages, involving drastic changes in development and life-history between closely related, genetically distinct species (*16–19*). Although they are phenotypically indistinguishable as larvae, they are very distinct as adults. In contrast to parasitic lampreys, non-parasitic species show a suite of paedomorphic changes, including loss of the juvenile feeding phase (non-trophic), intestinal degeneration during metamorphosis, residency in natal streams, and accelerated sexual maturation (Fig. 1A) (*16*, *20*). This makes non-parasitic lampreys to our knowledge the only vertebrates not feeding as juveniles and/or adults, and creates a coordinated life-history syndrome (referred to as parasitic vs non-parasitic). Lampreys have fascinated biologists for over a century (*21*) and they become increasingly important for comparative genomic studies to elucidate the evolution of key vertebrate features (*22*, *23*). Yet, the genomic basis of this fascinating life-history syndrome within this early-diverging jawless vertebrate lineage remains unresolved.

**Figure 1:**
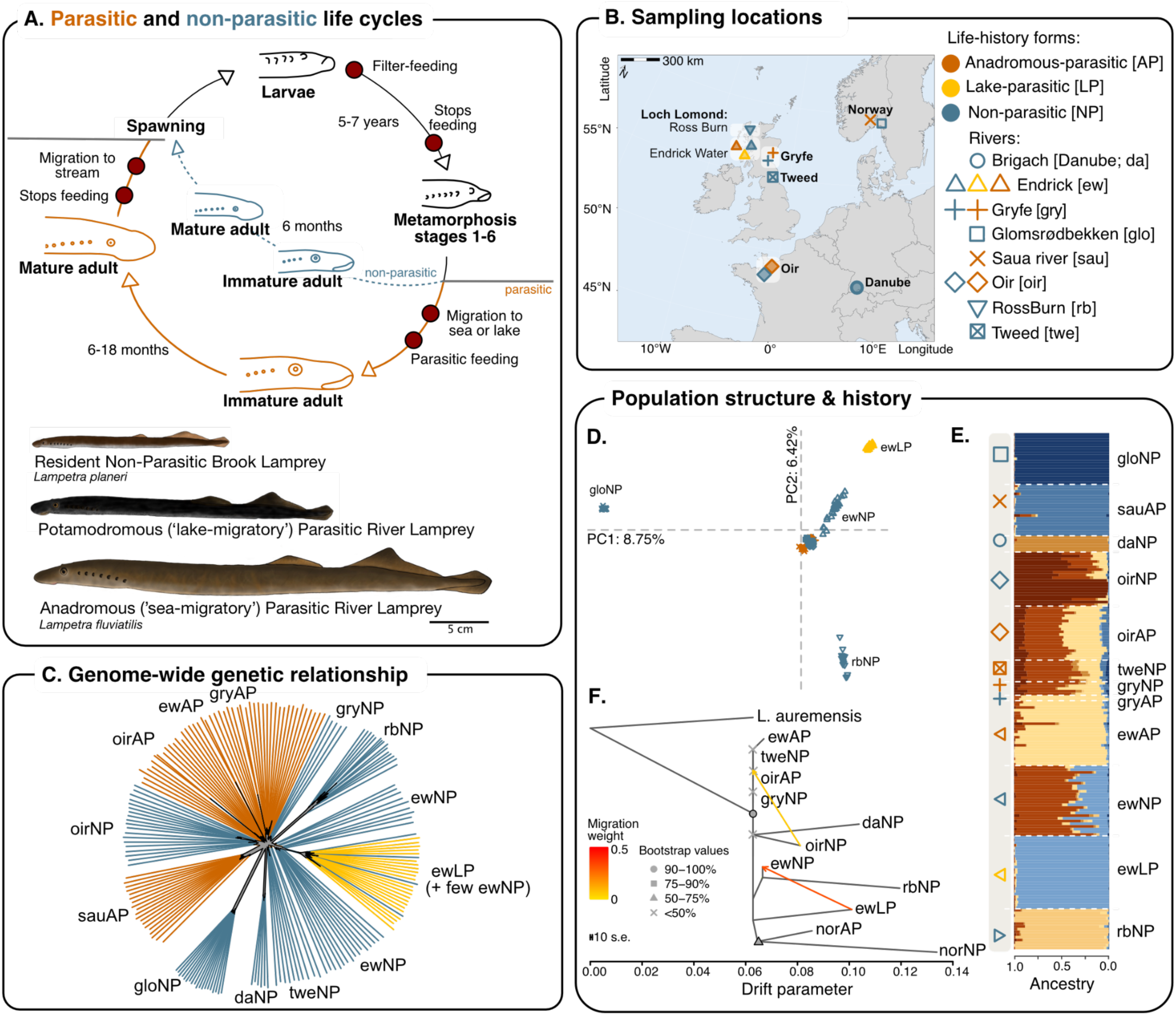
Life-history variation and population structure of European lampreys. **(A)** Schematic representation of the life cycle of parasitic (orange) and non-parasitic (blue) European lampreys, highlighting how life-histories diverge between ecotypes at the end of metamorphosis. The drawings below are representative examples of the three ecotypes from the river Endrick, UK, with taxonomic classification (by A Jacobs). **(B)** Map of Europe showing the sampling locations of individuals used for whole-genome sequencing. The exact locations are given in Table S1. Life-history forms are highlighted by colour, whereas different populations are distinguished by shape. **(C)** NeighbourNet network based on genetic distances calculated for neutral SNPs (thinned, non-divergent SNPs outside high-LD regions) showing the phylogenetic relationship between ecotypes across populations. The grey area shows the phylogenetic outline to highlight close relationships, with the exact phylogenetic network topology shown in fig. S1. A network for thinned non-divergent SNPs is shown in fig. S1. Branches are coloured by ecotype and populations are named as shown in panel (B). **(D)** Principal components analysis based on thinned, non-divergent SNPs for all ecotypes and populations. Additional PCAs and eigenvalue distributions for all SNPs and thinned, non-divergent SNPs are shown in fig. S3. PCAs and ancestry plots for the Loch Lomond populations and Oir population are given in fig. S5. **(E)** Genome-wide ancestry estimated based on thinned, non-divergent SNPs with *PCAngsd* (K=8). Individuals are ordered by ecotype (coded by colour) and populations (shape) (see panel (B) for detail). Estimates for other levels of K are shown in fig. S4. **(F)** Genome-wide maximum-likelihood tree with bootstrap support and the two strongest gene flow events as estimated with *TreeMix* based on neutral SNPs. Populations are named following the legend in (B). *TreeMix* trees with different numbers of migration events and residual correlations are provided in fig. S6.

In this study, we take advantage of the recent loss of parasitism in the European lamprey species complex (*Lampetra* spp.) to resolve the underlying genomic basis by integrating comparative genomics of multiple haplotype-resolved genome assemblies, population genomics of parasitic and non-parasitic individuals from across Europe, targeted genotyping in a narrow hybrid zone, and transcriptomic analyses of multiple tissues. We show that life-history variation is associated with a multitude of major effect loci concentrated in a few genomic regions containing crucial genes for development, behaviour and metabolism. Interestingly, we find a major parasitism-associated region with low recombination that is maintained in the absence of genomic rearrangements between ecotypes despite ongoing gene flow, suggesting a crucial role of ancient interspecific rearrangements in intraspecific evolution.

## Results

### Rapid evolution of European lamprey life-history diversity under ongoing gene flow

European lamprey life-history forms are taxonomically divided into the ancestral anadromous-parasitic river lamprey (*Lampetra fluviatilis*), lake-migratory parasitic river lamprey (*L. fluviatilis*), and the resident non-parasitic brook lamprey (*L. planeri*), (referred to as ‘ecotypes’ hereafter) (*16*, *19*, *24*). These ecotypes have likely evolved recently (<1 MYA) (*17*) and frequently hybridise under secondary contact (*24*, *25*) due to incomplete reproductive isolation (*24*, *26*, *27*). We investigated the evolutionary relationship of ecotypes from eight rivers across Europe (different combinations of ecotypes) using whole genome sequencing data for 189 individuals (mean coverage depth of 8.4x; range: 2.7x to 24.3×) (Fig. 1A,B table S1). Genome-wide phylogenetic relationships based on neutral SNPs (see methods for SNP filtering) suggested a recent and rapid divergence of ecotypes within and across rivers, as individuals largely cluster by ecotype and river but are only weakly divergent, as highlighted by a star-shaped phylogeny with high uncertainty in phylogenetic clustering (Fig.1C, fig. S1). This pattern is similar for all SNPs across the genome, with slightly stronger clustering by river and ecotype (fig. S1). The lack of ecotype clustering across rivers and uncertain phylogenetic relationship is consistent with the rapid colonisation of rivers and repeated divergence from a shared common ancestor, which is supported by a lack of clustering by ecotype or river for mitochondrial genomes (fig. S2). An exception is the lake-parasitic Endrick ecotype, which was highly divergent from other parasitic populations, suggesting an independent origin of this lake-specialised ecotype (Fig. 1C,D, fig. S3-S6). Yet, the non-parasitic Endrick ecotype showed strong signals of introgression from the sympatric lake-parasitic ecotype (Fig. 1D-F, fig. S5-S6) (*24*). Furthermore, the longer branch lengths (fig. S1 and S6) and stronger divergence (Fig.1D; fig. S4) of non-parasitic populations without sympatric parasitic individuals (e.g. daNP, rbNP, gloNP, names in Fig. 1B), compared to those in sympatry with parasitic lampreys, suggests strong isolation and genetic drift without recurrent genetic influx from the anadromous forms (*28*). Closer comparisons of sympatric ecotypes showed strong historical and ongoing admixture in sympatry (Fig. 1D-F, fig. S3-S6), with non-parasitic ecotypes displaying >25% anadromous-parasitic and lake-parasitic ancestry in the Oir and Endrick, respectively (Fig. 1E; fig. S5). Parasitic forms showed weaker proportions of non-parasitic ancestry (Fig.1E, fig. S5), in line with size-related mating behaviour between ecotypes (*24*, *29*).

### Major-effect loci control life-history divergence

To identify the genomic basis of life-history, we performed genome-wide association analysis across all parasitic (n=91) and all non-parasitic (n=98) individuals. The presence of genetic admixture between ecotypes, replication across watersheds, and the ability to control for differences in migration strategy through parasitic ecotypes with different migration tactics (lake-parasitic vs anadromous-parasitic; Fig.1B) makes this a powerful approach. We identified seven genomic regions on six chromosomes with significant life-history associations (p < 10^-8^) (Fig. 2A) (these binary-trait quantitative trait loci are hereafter referred to as ‘parasitism-QTL’). The strongest associations (p=1.42e-32) were in a narrow ∼100kb region on chromosome (chr) 30, yet most significant SNPs (7550 out of 7812 significant SNPs) mapped to a ∼20Mb region on chr1, structured into seven individual peaks (Fig. 2A,B; fig. S7). Furthermore, we identified a broad ∼5Mb parasitism-QTL on chr56 and three weaker parasitism-QTL on chr6, chr8 and chr25. The parasitism-QTL on chr1 and chr30 were also significantly associated with life-history across all three admixed sympatric ecotypes in the Endrick (fig. S8), supporting their consistent role in this life-history shift.

**Figure 2:**
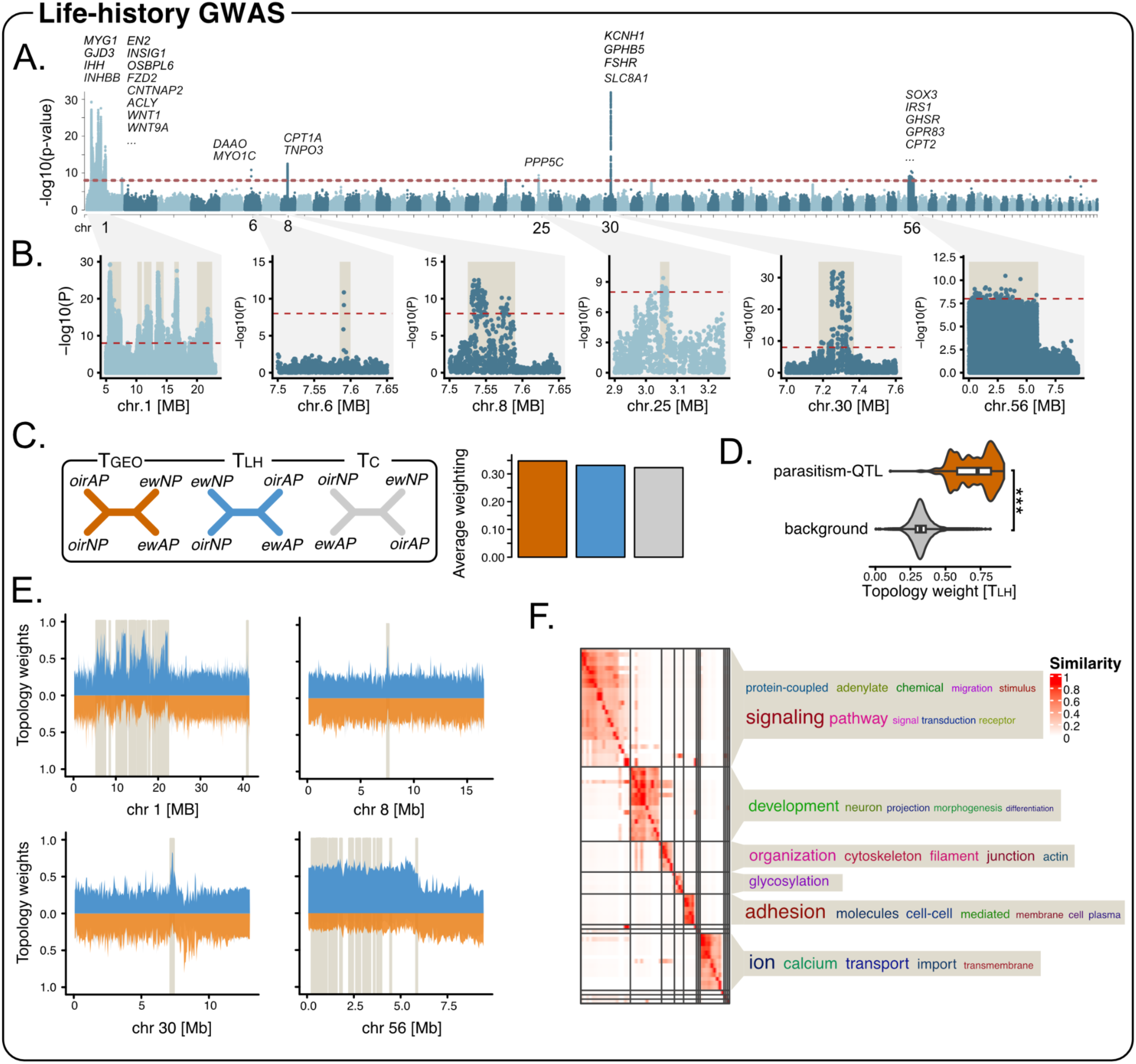
Genomic signals of parasitism-loss in European lampreys. **(A)** Manhattan plot showing genetic associations for life history (parasitic vs non-parasitic) estimated using a linear mixed model in *GEMMA* across all populations. The red dashed line shows the significance threshold of p = 10^-8^. We show some key candidate genes with parasitism-QTL above individual peaks (not all shown for chr1 and chr56). The full list of candidate genes, their full names, and locations are given in Supplementary file 1, with functional annotations in Supplementary file 1. **(B)** Candidate regions on chromosomes with parasitism associated genomic regions (“Parasitism-QTL”), zoomed in from the plot above (A). **(C)** Four-taxon topologies for the anadromous-parasitic (AP) and non-parasitic (NP) ecotypes from the Endrick (ew) and Oir that were tested using *TWISST*. T_GEO_ shows the geographic topology, clustering individuals by location of origin; T_LH_ shows the life-history topology, clustering individuals by ecotype; T_C_ shows the control topology without any clustering by geography and ecotype. The barplot on the right shows their average weighting by topology (proportion across all trees) across the genome. **(D)** Comparison of the life-history topology (T_LH_) weights for 50kb genomic windows overlapping parasitism-QTL and the genomic background (genomic regions without parasitism-QTL) across the genomes. The genomic background distribution was obtained by the random resampling with replacement of x loci for 1,000 times (x = number of windows overlapping parasitism-QTL). Asterisks highlight the significance level for differences in topology weight distributions estimated using a Kruskal-Wallis test (W=145644457, p-value < 2.2e-16). **(E)** Distribution of topology weights across the four chromosomes (chr 1, 8, 30, 56) with top parasitism-QTL (highlighted by brown rectangles). To aid visualisation, the T_LH_ (blue) distribution is upwards, whereas the distribution for T_GEO_ (orange) is plotted downwards. **(F)** Semantic similarity clustering of overrepresented biological process gene ontology (GO) terms for genes overlapping parasitism-QTL, with similarities shown using a white to red scale. Descriptions of GO terms are shown on the right, with the size of words describing overrepresented terms shown relative to their frequency. Full GO term overrepresentation results are shown in Supplementary file 1 and fig. S12.

To further test if parasitism-QTL are consistently involved in life-history evolution across independent watersheds, we performed topology-weighting analyses across sympatric life-history pairs, which test the degree of monophyly of the three possible phylogenies across genomic regions (*30*, *31*). These tests suggested that haplotypes around parasitism-QTL were largely shared across populations (Fig 2C). Parasitism-QTL showed a higher weighting for the life-history topology (T_LH_), which clustered individuals by life-history across rivers, compared to the geographic topology (T_Geo_), which clustered individuals by river of origin (Kruskal-Wallis test for T_LH_ weights: W=145644457, p-value < 2.2e-16) or the control tree (T_C_), despite T_Geo_ being the most common topology across genome, likely due to ongoing gene flow (Fig. 2C,D, fig. S9). A skew toward the T_LH_ topology across all parasitism-QTL is unlikely under incomplete lineage sorting (ILS) (*TWISST’N’TERN*: D_LR_=-0.0177; P=0.015, fig. S10), as all topologies are equally likely under ILS, suggesting the maintenance of parasitism-associated regions through selection (*31*). Peaks in T_LH_ weighting around parasitism-QTL on chr1 and chr30 suggested that ecotypes share similar haplotypes in these regions (Fig. 2E), with the lack of fixation for the T_LH_ topology (T_LH_ weight < 1; Fig. 2D; fig. S9) indicating the presence of genetic variation in parasitism-QTL across rivers, e.g. through genetic drift and gene flow.

The repeated loss of parasitism via similar phenotypic changes across independent lamprey taxa (*16*), might indicate a shared genomic basis (*32*). However, comparisons to parasitic and non-parasitic ecotypes from a North American sister lineage to the European lamprey (*Occidentis ayresii*) (*33*), which diverged ∼7.5MYA (*17*), indicated that genomic regions homologous to parasitism-QTL in European *Lampetra* were not differentiated between *O. ayresii* ecotypes (fig. S11). This suggests that parasitism was lost through alternative genomic routes (*17*, *33*). Yet, similar to European lamprey, *O. ayresii* ecotypes seemed to be differentiated by a mosaic of genomic architectures, including putative structural variants (fig. S11 B,C), indicating comparable genomic architectures.

### Parasitism-QTL are associated with major developmental and metabolic pathways

The diversity of genes under parasitism-QTL (Fig. 2A, Supplementary file1) and associated gene ontology terms (Fig. 2F, fig. S12, Supplementary file 1) reflect the myriad phenotypic changes associated with parasitism loss in lampreys, including the loss of juvenile migration and feeding, developmental variation during metamorphosis, earlier sexual maturation, and drastic changes in metabolism in non-parasitic lampreys (*16*, *34*). For example, the strongest peak on chr30 contained a glycoprotein hormone gene (*GPHB5*) and follicle-stimulating hormone-receptor gene (*FSHR*) (*35*), which are crucial for sexual development and reproduction, known key traits distinguishing parasitic and non-parasitic lampreys (*16*, *36*). Furthermore, the peak on chr1 contained the *INHBB* gene, which is part of the TGF-ꞵ superfamily and encodes a highly-conserved protein involved in reproductive development and fertility, e.g. by regulating follicle-stimulating hormone secretion (*37*). The dual effect on the FSH pathway might indicate a potential functional interaction between divergent genes on chr1 and chr30. Many genes on chr1 play key roles in major signalling pathways involved in development (e.g. Hedgehog; Transforming growth factor (TGF)-ꞵ; WNT; Insulin-like Growth Factor (IGF)) (*38–41*). This included thyroid-hormone responsive genes (e.g. *IHH, GLI2, GALNT12, WNT1*), with thyroid hormone being the master regulator of metamorphosis (*42*); the developmental timepoint at which phenotypic differences between ecotypes arise (*16*, *20*, *34*). Parasitism-QTL further overlapped numerous metabolic genes involved in fatty-acid metabolism and glycolysis on chr1 (e.g. *HACL1, INSIG2, OSBPL6, PHYH, PDK2, PFKP*), chr8 (*CPT1A),* chr25 (*PPP5C*), and chr56 (*CPT2*, *IRS1*) (Supplementary file 1). This suggests that ecotypes likely differ in metabolism (*43*, *44*), which aligns with differences in trophic ecology; non-parasitic lamprey do not feed as juveniles, while the parasitic lamprey actively migrate, parasitically feed and rapidly grow (*16*, *36*). Additionally, we identified genes with known roles in feeding behaviour/appetite on chr1 and chr56 (*INSIG2, GHSR*, *GPR83).* Lastly, many parasitism-QTL genes were associated with brain/neural development and function (e.g. *AMPH, EN2, CNTNAP2, NLGN3, DLG2*), pointing toward differences in brain development between ecotypes, with the brain drastically restructuring during lamprey metamorphosis, reflecting ontogenetic shifts in behaviour and ecology (*45*).

The functional role of genes under parasitism-QTL was further investigated through differential expression analyses in several tissues (i.e. brain, liver, gonad and gill) between spawning-ripe anadromous-parasitic and non-parasitic individuals from the Oir (fig. S13). Of the 177 genes under parasitism-QTL, 47 (50 transcripts) were differentially expressed in at least one tissue (ranging from 2 in the gonad to 34 in the liver; fig. S13), including key metabolic genes in the liver, such as *CPT1A* and *CPT2*, which play a role in fatty-acid metabolism, and *IRS1*, an insulin receptor. It is important to note that we only assayed gene expression in adults, and other genes under parasitism-QTL might be either differentially spliced (*46*, *47*) or differentially regulated in other life stages (*48*).

### Interspecific genomic rearrangements are associated with recombination cold spots

To elucidate how parasitism-QTL are maintained despite extensive gene flow, we tested if they colocalised with recombination-suppressing chromosomal rearrangements between ecotypes (on chromosomes 1, 25, 30, 56). The presence of three distinct genomic clusters in PCAs for parasitism-QTL on chr56 indicated a large rearrangement, and a smaller putative rearrangement around the parasitism-QTL at the end of chr1 (Fig. 3A, fig. S14-15). Although some of the other regions display three broad clusters in region-specific PCAs, these still show evidence of recombination (e.g. intermediate individuals between clusters) and are likely associated with low-recombination regions but not recombination-suppressing SVs (*1*). Comparisons of five genome assemblies for both ecotypes, including new haplotype-resolved, chromosome-scale genomes for the parasitic and non-parasitic ecotypes from Scandinavia (*49*) and France (Oir, provided in this study) (table s4), and genomes for two evolutionarily distant species [sea lamprey (*Petromyzon marinus*) and Korean lamprey (*Lethenteron reissneri*)], that diverged between 5 and 25MYA, (*17*, *50*, *51*), confirmed the presence of chromosomal rearrangements between ecotypes on chr56, but did not provide evidence for large rearrangements on chr1 or the other chromosomes (Fig. 3B, fig. S16-17). While most of the genome showed high collinearity across *Lampetra* genomes (fig.S19), we detected a ∼3Mb inverted-translocation (*‘trans-inv56*’) between ecotypes on chr56 (Fig. 3B, fig. S17). The same region was also translocated across more distantly related species, indicating high structural turnover (Fig. 3B). *Trans-inv56* showed high linkage disequilibrium (fig. S18) and low recombination rates in the non-parasitic Oir ecotype (Fig. 3C, fig. S19-S20, fig. S21-22 for linkage maps). However, population-level recombination rates were only slightly decreased in the non-parasitic Endrick ecotype (Fig. 3C, fig. S19), potentially because this ecotype harbours all three *trans-inv56* karyotypes.

**Figure 3:**
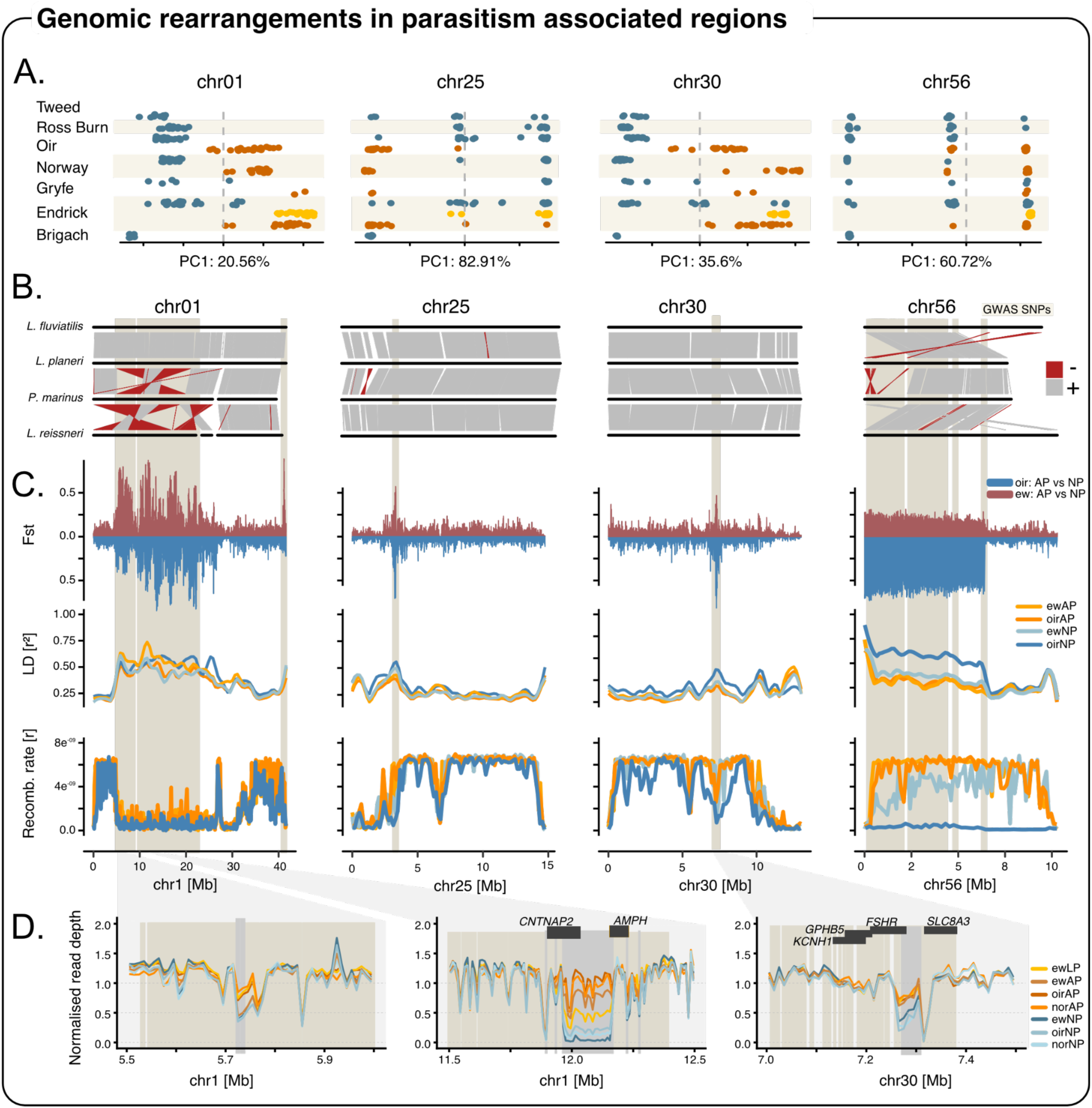
Genomic rearrangements around parasitism-QTL. **(A)** Genomic principal component scores (PC1) showing the genetic variation for all SNPs within parasitism-QTL by population for chrs. 1, 25, 30, 56. Eigenvalues (in %) of PC1 for each genomic region are shown below each plot. Each dot represents an individual coloured by ecotype and ordered by population (n=189, n= 2 - 27 by ecotype and population, see table S1). Anadromous-parasitic = orange, lake-parasitic = yellow, non-parasitic = blue. The plot for chr1 is across all peaks between 5Mb and 22Mb. PCA plots (PC1 vs PC2) for each parasitism-QTL are shown in fig. S14-S15. **(B)** Genomic synteny between the anadromous-parasitic Endrick ecotype (kcLamFluv1.1 assembly), non-parasitic Oir ecotype (*L. planeri* - haplotype 2), sea lamprey (*P. marinus* - kcPetMar1) and Korean lamprey (*L. reissneri -* ASM1570882v1), estimated with *ntSynt*. Shown are chromosomes (chrs. 1, 25, 30, 56) with major parasitism-QTL (highlighted in beige), ordered by the extent of synteny. Grey bars connect syntenic, collinear regions, whereas red-bars show inverted regions with reversed strands. Synteny analyses across the entire genome and additional assemblies are shown in fig. S16 and S17. **(C)** Top: Genetic differentiation (Fst) between anadromous-parasitic and non-parasitic ecotypes in the Endrick (ewAP, ewNP) and Oir (oirAP, oirNP) across the four chromosomes containing the main parasitism-QTL. Fst between Endrick ecotypes is plotted upwards and for Oir ecotypes downwards to aid visualisation (both on the same scale from 0 to 1). Middle: Linkage disequilibrium (LD) values (r^2^) for replicated ecotypes from the Oir and Endrick summarized in 10kb windows across each chromosome and plotted using locally estimated scatterplot smoothing with a 0.1 span. Lines are coloured as shown in the legend. Bottom: Recombination rates for anadromous-parasitic and non-parasitic ecotypes from Endrick and Oir across candidate chromosomes, with lines coloured the same way as LD values. **(D)** Evidence for copy number variation within parasitism-QTL. Left plot: Normalised read depth indicates reduced read depth in non-parasitic compared to parasitic ecotypes within a narrow region on chr1 as highlighted by a grey bar. This is supported by increased divergence in read depth between ecotypes across populations as measured using read-depth based Qst estimates (fig.S29). Middle plot: Variation in normalised read depth indicates the presence of a large deletion in the non-parasitic ecotype in the Endrick (ew), Oir (oir) and Norwegian (nor) rivers at around 12Mb on chr1 (grey bar). The deletion is located between two parasitism-QTL peaks, but without any significant SNPs within the deletion region as no sites were called in non-parasitic individuals in this region. Genes (*CNTNAP2 and AMPH*) partially overlapping this putative deletion are shown as dark grey rectangles at the top. Right plot: The parasitism-QTL on chr30 shows signs of reduced read depth in non-parasitic ecotypes, which could indicate a heterozygous deletion in this ecotype. The putative deletion overlaps the follicle-stimulating hormone receptor gene (*FSHR*).

Despite the lack of evidence for large genomic rearrangements between ecotypes around the other parasitism-QTL (fig. S16-S17) recombination rates were generally lower around parasitism-QTL compared to the genomic background (fig. S20), and these regions also showed increased Fst and LD (Fig. 3C). Compared to most of the genome, the low-recombination region on chr1 showed substantial interspecific genomic rearrangements across lamprey lineages, including fusions, translocations and inversions, but no rearrangements between parasitic and non-parasitic European lamprey ecotypes (Fig.3B, fig. S17). We hypothesise that these ancient interspecific rearrangements have potentially led to long-term recombination suppression on chr1, e.g. by altering the chromatin landscape (*12*, *52*), contributing to the maintenance of co-adapted alleles under gene flow (*10*, *13*, *53*).

While we did not detect chromosomal rearrangements between ecotypes around most parasitism-QTL, except chr56, we found evidence for divergent copy-number variants (CNVs) between sympatric ecotypes based on normalised read depth (Fig. 3D; fig. S23), which were most likely deletions in non-parasitic lamprey compared to the parasitic reference. Three divergent CNVs (CNV-based QST > mean SNP-based Fst) between ecotypes were located within parasitism-QTL on chr1 and chr30 (Fig. 3D). Two of these CNV partially overlapped annotated genes, *CNTNAP2* and *AMPH* on chr1, which have known roles in brain development and neural function, and *FSHR* on chr30, alluding to a putative functional role of CNVs in ecotype divergence.

### Maintenance of life-history variation through selection in parasitic ecotypes

We hypothesised that selection would act more strongly on parasitic ecotypes to maintain co-adapted trait combinations related to feeding and migration. In contrast, trait loss, as observed in non-parasitic ecotypes (e.g. skipping the juvenile migration and feeding phase), has regularly been linked to relaxed selection (*16*, *54*, *55*). Parasitism-QTL showed higher levels of genetic differentiation compared to the genomic background (Fig. 4 A,B, fig. S24), with genetic differentiation being moderately correlated across ecotype pairs in the Oir and Endrick (Pearson correlation of Fst: r = 0.37, t = 163.28, df = 175083, p-value < 2.2e-16). However, not all parasitism-QTL were strongly differentiated in all ecotype pairs, e.g. with *trans-inv56* only showing weak differentiation between ecotypes in the Endrick compared to strong differentiation in the Oir (Fig. 3C,4A, fig. S25), which is likely explained by the presence of all three karyotypes in the non-parasitic Endrick ecotype. In contrast, parasitic individuals were either homozygous for the “parasitic” karyotype (higher frequency in parasitic ecotype; AP*^trans-inv56^*/AP*^trans-inv56^*) or heterozygous (NP*^trans-inv56^*/AP*^trans-inv56^*), but never homozygous for the “non-parasitic” karyotype (NP*^trans-inv56^*/NP*^trans-inv56^*), suggesting potential selection against the non-parasitic karyotype in parasitic lampreys. Thus, we argue that *trans-inv56* might not determine ecotype but rather impacts one or multiple ecotype-linked phenotypes (Fig. 3A, fig. S14). Overall, genetic differentiation within parasitism-QTL was higher between parasitic and non-parasitic ecotypes than between sympatric parasitic ecotypes (i.e. anadromous-vs lake-parasitic in the Endrick) (Fig. 4B), supporting their general role in parasitism-loss.

**Figure 4:**
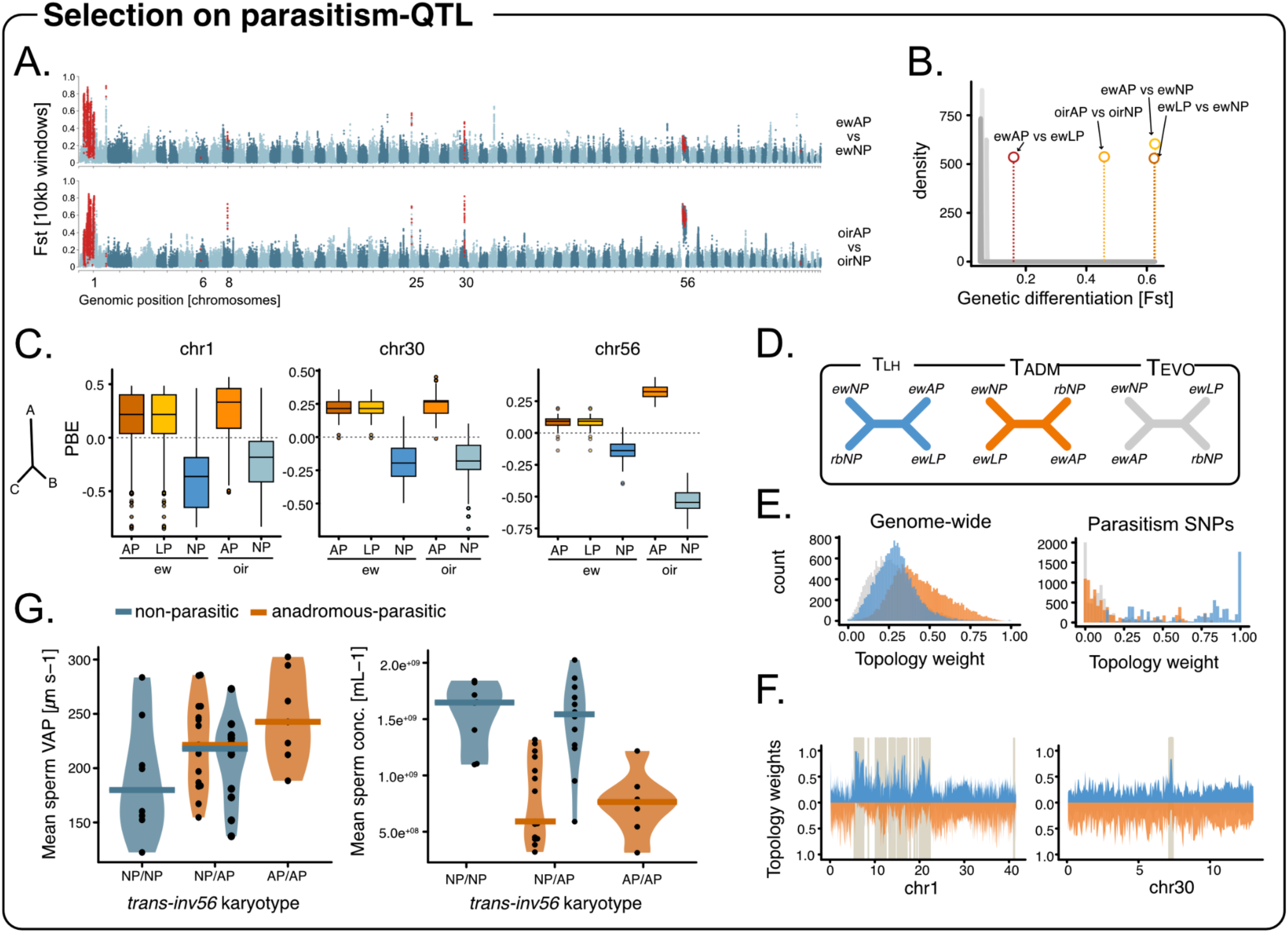
Selection on parasitism loci. **(A)** Landscape of genetic differentiation (Fst) in 10kb sliding windows (5kb steps) between sympatric life history forms in the Endrick (Top: Anadromous-parasitic vs non-parasitic) and the Oir (Bottom: Anadromous-parasitic vs non-parasitic). Genomic windows overlapping parasitism-QTL are highlighted in red. **(B)** Higher genetic differentiation of genomic windows overlapping parasitism-QTL compared to the genomic background. Comparisons are shown between all ecotypes in the Endrick (ew) and the Oir ecotypes. Grey distributions show the resampled distribution of mean Fst values for n windows per comparison (n = number of parasitism-QTL overlapping genomic windows). **(C)** Higher Population Branch Excess (PBE) in parasitic compared to non-parasitic ecotypes for genomic-windows overlapping parasitism-QTL on chr1, chr30 and chr56, which show some of the strongest signals of ecotype association and/or genetic differentiation. The drawing on the left shows the comparison, with the focal population A displaying a longer branch length compared to outgroups C and B, indicative of positive selection. PBE scores for all parasitism-QTL are shown in fig. S27. **(D)** Topologies for the three Endrick ecotypes and the non-parasitic ecotype from a neighbouring stream (Ross burn; rbNP) used in *TWISST*. The T_LH_ topology shows the phylogenetic relationship expected for clustering by life-history, the T_ADM_ topology shows the relationship expected based on the genetic admixture between lake-parasitic and non-parasitic Endrick ecotypes, and the T_EVO_ topology shows the phylogenetic relationship between ewNP and ewAP expected based on their evolutionary relationship. **(E)** Distribution of topology weights for each topology (in D) across the entire genome (left) and only genomic windows overlapping parasitism-QTL (right). Distributions are coloured according to their topology. **(F)** Genome scan of topology weights in 50kb windows across chr1 and chr30, with topology weights for the life-history topology (T_LH_; see D) shown in blue above 0 and the admixed (orange) and alternative (grey) topologies below 0. Parasitism-QTL are shown in black and broader parasitism-QTL regions highlighted as brown boxes in the plot. **(G)** Differences in mean sperm average path velocity (VAP) and mean sperm concentration between parasitic and non-parasitic individuals from the Oir with different karyotypes for *trans-inv56*. NP/NP = homozygous for the karyotype with higher frequency in non-parasitic populations; AP/AP = homozygous for parasitic karyotype; NP/AP = heterozygous. Violin plots are coloured by ecotype (see legend), with individual datapoints in black (n=42) and coloured bars showing the mean for each group (karyotype by ecotype).

In line with our above predictions on selection, we found that parasitism-QTL on chr1, chr8, chr30 and chr56 showed stronger signatures of positive selection (higher population branch excess, PBE) in parasitic ecotypes compared to non-parasitic ecotypes in the Oir and Endrick (Fig. 4C, fig. S26-S27). In contrast, the parasitism-QTL on chr25, containing the *PPP5C* gene, which is linked to fat metabolism and glucose balance (*56*), showed stronger signatures of selection in non-parasitic lamprey, suggesting the potential alteration of metabolic traits with the loss of parasitism.

Lastly, we made use of the extensive introgression from the lake-parasitic into the non-parasitic Endrick ecotype to test for signatures of selection against admixture around parasitism-QTL (Fig. 1D-F), which we argue would manifest in the (near) fixation of life-history topologies (T_LH_) around parasitism-QTL despite genome-wide admixture. Topology weighting analyses across all three Endrick ecotypes and an adjacent non-admixed, non-parasitic population (Ross Burn; Fig. 4D) showed that a topology clustering the lake-parasitic and non-parasitic Endrick ecotypes (Admixed topology: T_ADM_) was most common across the genome (Fig. 4E), which is expected due to their strong introgression (*24*). However, parasitism-QTL on chr1 and chr30 were enriched for windows with fixed or nearly fixed life-history topologies (T_LH_) (Fig. 4E,F, fig. S28), as expected for selection against admixture in those regions (*31*).

### Trans-inv56 is associated with ecotype divergence in sperm velocity

To better understand patterns of admixture and the role of *trans-inv56*, we genotyped 1072 individuals (945 adults and 127 larvae of unknown ecotype) from a hybrid zone in the Oir at 131 SNPs, targeting SNPs within parasitism-QTL on chr1 (n=9 SNPs; fig. S29A), *trans-inv56* (n=21 SNPs, fig. S29A) and non-divergent SNPs outside parasitism-QTL (‘neutral’: n=101 SNPs). This confirmed high levels of genome-wide admixture between ecotypes (fig. S29A), although proportions of admixed individuals with ancestry proportions (Q) above or below 25% and 75%, respectively, varied between life-stages (fig. S29). Of parasitic and non-parasitic adults, 48.9% and 58.9% were classified as admixed, respectively (adult mean: 53.9%), whereas 79.5% of larvae were admixed (Adult vs larvae admixture proportion: *X*^2^ = 28.8, df = 1, p = 8e-08). A decrease in admixture with age could point toward potential viability selection against hybrids at later life stages (*57*).

Individual-based simulations of hybridization between parasitic and non-parasitic lampreys (fig. S30) suggest that without selection or hybrid breakdown, the non-parasitic ecotype nearly disappears, likely due to the higher fecundity of the anadromous-parasitic ecotype. Thus, similar to previous models (*58*), our simulations indicate the need for selective advantages and / or selection against hybrids to explain the persistence of the non-parasitic ecotype in sympatry.

Despite the potential selection against hybrids, we detected an excess of *trans-inv56* heterozygotes (NP*^trans-inv56^*/AP*^trans-inv56^*) in both ecotypes, which was stronger in the non-parasitic ecotype (fig. S31). This could potentially indicate overdominance (i.e. heterozygote advantage) in non-parasitic lamprey. In line with this overdominance hypothesis, we found that non-parasitic heterozygotes (NP*^trans-inv56^*/AP*^trans-inv56^*) have faster but not more sperm compared to homozygotes (NP*^trans-inv56^*/NP*^trans-inv56^*) (LMM: beta = 27.72, t(_38_) = 2.50, p = 0.017; Fig. 4G; fig. S32), a likely fitness advantage in communal spawners and for smaller sneaker males (*59*). In contrast, heterozygous parasitic males (NP*^trans-inv56^*/AP*^trans-inv56^*) had slower sperm compared to homozygous males (AP*^trans-inv56^*/AP*^trans-inv56^*) (Fig. 4G), supporting a scenario of selection against the non-parasitic karyotype in the parasitic ecotype. A role of *trans-inv56* in sperm velocity is further supported by the presence of genes linked to sperm traits inside the inversion (e.g. *CUL4B*, *MFF*, CPT2, *IRS1*; table S2), providing key candidate genes underlying variation in sperm velocity between lamprey ecotypes, and targets for future functional studies.

## Discussion

Our study resolves the genomic basis of a striking life-history transition in an early-diverging vertebrate lineage; the repeated loss of parasitism in lampreys. Rather than the single supergene-style architecture often inferred for analogous transitions in other taxa, such as large inversions controlling life-history variation in rainbow trout (*6*), ruff (*7*), or cod (*5*), or the highly polygenic basis of migration-history in brown trout (*60*), we find that this life-history syndrome in lampreys maps to a small number of regions with distinct genomic architectures. Below we consider, in turn, what these regions implicate biologically, how their architectures are maintained, and what their non-shared identity in a North American sister lineage reveals about the repeatability of this transition.

The genes implicated by these regions cluster around three major axes that mirror the components of the syndrome itself. The peak on chr30, including a putative deletion in non-parasitic lampreys, overlaps key genes in the gonadotropin pathway (*FSHR*, *GPHB5*), a central regulator of vertebrate sexual maturation (*61*). This is consistent with the accelerated post-metamorphic maturation that distinguishes non-parasitic from parasitic lampreys (*16*). Direct functional evidence in fish establishes FSHR as a key component of maturation transition, with *fshr*-knockout in Atlantic salmon preventing testicular maturation (*62*), and *fshr* loss-of-function in medaka causing folliculogenesis arrest (*63*). In humans, FSHR variants are associated with timing of pubertal onset and reproductive length (*64*, *65*). Notably, the dominant maturation-timing locus in salmonids (*vgll3*) (*66*, *67*) acts upstream of the gonadotropin axis, whereas in European lamprey variation in sexual maturation seems to be orchestrated through changes in the receptor itself. Furthermore, genes under parasitism-QTL on chr1 and within the chr56-inversion implicate neural development and synaptic function (e.g. *CNTNAP2*, *KCNH1*, *AMPH*, *EN2*, *GSHR*, *WNT1*), pathways plausibly associated with the transition from a migratory and predatory behaviour in parasitic lampreys to a brief, stream-resident juvenile phase in non-parasitic lampreys (*16*). Metabolic genes under parasitism-QTL on e.g. chr1, chr8 and chr56, that were also differentially expressed in livers (e.g. *CPT1A*, *CPT2*, *IRS1*, *INSIG1*) further implicate fatty-acid β-oxidation and insulin signalling, consistent with the metabolic transition away from juvenile feeding in non-parasitic lampreys toward reliance on stored reserves through metamorphosis and spawning (*16*, *43*, *44*). This convergence between gene function and phenotypic axis points toward specific developmental and metabolic pathways that must be co-modified for this life-history transition to occur. However, the underlying mechanistic changes regulating this life-history shift remain unresolved.

Yet, how is genetic divergence between lamprey ecotypes maintained in the face of gene flow? Most parasitism-QTL on chromosome 1 fall within a ∼20 Mb region of suppressed recombination that does not seem to be caused by large-scale intra-specific rearrangements. Yet, the same region carries inter-specific rearrangements. One parsimonious interpretation is that these ancestral rearrangements established a recombination cold spot through altered chromatin architecture, which has subsequently captured and held together ecotype-divergent variation. This is consistent with recent work on ancient rearrangement-derived recombination cold spots in mammals (*12*, *13*) and the suggested impact of ancient chromosomal fusions on adaptation in other systems (*10*, *52*, *53*, *68*). However, the link between low-recombination regions due to ancient rearrangements and adaptive divergence has been relatively unexplored so far, and direct functional tests are needed to establish mechanistic links between these regions and phenotypic change. Altogether, these observations suggest that ancient, fixed rearrangements may act as a substrate for the maintenance of adaptive variation under gene flow in a manner that is mechanistically distinct from, but functionally analogous, to the well-described inversion supergenes. In contrast, a large translocated-inversion on chr56 (trans-inv56) seems to contribute to life-history variation by affecting variation in sperm velocity, an ecotype-related trait (*59*) that is critical for sperm competition in externally fertilising species (*59*, *69*, *70*). Interestingly, the genomic basis of sperm trait evolution has rarely been dissected within or between species in the wild (*71*, *72*), and links to chromosomal inversions are rarer still (*72*). The persistence of karyotype differences between ecotypes, despite the excess of heterozygotes, might indicate the presence of evolutionary trade-offs, a balanced-polymorphism mechanism well-established for inversion supergenes (*73–75*). Together with the chr1, trans-inv56 illustrates that different forms of recombination-suppressed architecture can coexist within a single life-history transition.

Lastly, our results for the North American sister lineage *Occidentis ayresii*, which has independently evolved a similar parasitic vs non-parasitic ecotype pair, indicates that the underlying loci are largely non-shared across lineages. Yet *O. ayresii* parasitism loss appears itself to involve a mosaic of broad and narrow regions of genetic differentiation, suggesting that the architectural mosaic is conserved across the genus while the specific loci are not. This pattern contrasts strikingly with canonical examples of repeatable parallel evolution at the locus level - *Eda* and *Pitx1* in three-spine sticklebacks (*76–78*) *or optix* in *Heliconius* (*79*) - and argues that life-history transitions of this type can recur through architecturally similar but locus-distinct genomic routes. Our comparisons across lineages further suggest that rearrangements on chr1 likely arose within the last 10 million years (*17*) and potentially following the split of *Lampetra* and *Occidentis* (divergence ∼8MYA) (*17*), consistent with this lineage-specific genomic architectures. We note, however, that the *Occidentis* comparison rests on low-coverage sequencing (∼0.5×) and that high-coverage re-sequencing will be required to definitively rule out shared loci of small effect. Future comparative and functional genomic studies - especially across more distantly related parasitic / non-parasitic species pairs and across the full lamprey phylogeny - will reveal whether mosaic architectures of this kind are a common feature of similar life-history transitions and provide insights into the mechanistic basis of parasitism loss and associated developmental changes in lampreys.

## Sequencing

We also want to thank Sophie Michon-Coudouel and Romain Causse-Vedrines (EcogenO Platform, University of Rennes) for the generation of the HiPlex data as well as Inês Trancoso for high molecular weight DNA extraction. The sequencing of Norwegian samples was performed by the Norwegian Sequencing Center (NSC).

Furthermore, we want to thank Kathryn Elmer and Thomas Boehm for help with sample collection, funding acquisition and data generation, Catherine Labbé for sperm speed analyses and the Evolution & Diversity Theme and Shared Interest Group at the University of Glasgow as well as Claire Mérot for stimulating discussions.

## Author contributions

Conceptualisation: AJ, GE; Sampling: AJ, GE, JT, JPD, QR, LAV, ND; Data generation: AJ, MC, ALB, SS, EJP, GL, DJP, BJP, GE, AS, QR, ND, RN, KSJ, SJ, SNKH, BGA; Analysis: AJ, ND, BGA, OKT, FG; Funding: AJ, GE, KSJ; Writing initial draft: AJ, GE; Writing final draft: All authors.

## Data and code availability

All data will be made available upon acceptance of the manuscript. The raw sequencing files will be accessible on ENA under the BioProject XXXXX. Additional data will be available on xxxx.

## Methods

### Sampling

European lamprey used in this study were collected using a range of methods in different waterways across Europe (Table S1). Sampling strategies are described below.

### UK

Juvenile and adult individuals were collected in the River Endrick, Gryfe and Ross Burn in Scotland (UK) using electrofishing, smolt traps or specialised lamprey traps (‘Maitland traps’) during the upstream migration (Autumn) between 2019 and 2022 (*80*). Ecotypes were identified based on size and colouration (Fig. 1A) (*24*, *81*), euthanised using an overdose of MS-222 and tissue samples collected for DNA extraction. Larvae were sampled in the River Tweed using electrofishing as described in (*82*) and whole-specimens frozen. These were determined to be non-parasitic, as sample sites were not reachable by anadromous-parasitic lamprey. Sampling was performed under the University of Glasgow ethical approval, with sampling permits from the Marine Directorate of Scotland.

### France/Oir

Parasitic and non-parasitic individuals were collected in sympatry in the Oir, a tributary of the Sélune, France, at spawning time as described in (*25*, *27*). Individuals were anesthetized with benzocaine to collect a fin clip, which was preserved in 95% EtOH.

### Norway

Non-parasitic individuals were sampled using electrofishing in Glomsrødbekken, Østfold county, and parasitic individuals were sampled using electrofishing in a tributary to the Saua River, Lake Nordsjø, Telemark county. The parasitic population presently have no or only limited access to the sea following construction of different dams and locks.

### Brigach, Danube

Larval samples were collected by the LAZBW in the Brigach, a tributary of the Danube in Germany, using electrofishing and whole larvae preserved in ethanol. These were determined to be non-parasitic, as anadromous-parasitic lampreys don’t exist in this system.

### Whole-genome resequencing

DNA was extracted from fin clips using either the Qiagen Blood & Tissue Kit or using a bead-based extraction protocol (dx.doi.org/10.17504/protocols.io.b46bqzan). Whole genome sequencing libraries for all individuals, except the Norwegian populations, were prepared using NEBnext Ultra II FS DNA library prep with 100ng of DNA as input and using half-reactions. Pooled libraries were sequenced on NOVASEQ X or NOVASEQ X Plus 25B lanes. Whole genome sequencing libraries for Norwegian samples were prepared at the Norwegian Sequencing Centre from DNA extracted using the Qiagen DNeasy Blood and Tissue Kit (from 25 mg muscle tissue) sequenced on an Illumina NovaSeq 6000 lane. We furthermore downloaded Illumina whole genome sequencing data for a single *Lampetra auremensis* individual from Portugal (SRR24224961), a sister species to *Lampetra fluviatilis/planeri*.

### WGS processing and SNP calling

Raw sequencing data for all individuals were processed and trimmed using FastP with the following settings: *-l 30 --trim_poly_g --cut_right --detect_adapter_for_pe* (*83*). We aligned processed reads to the Darwin Tree of Life *Lampetra fluviatilis* reference genome (kcLamFluv1.1; GCA_964198595.1 on NCBI) using *bwa-mem* (*84*), merged replicated samples using the merge command in *sambamba* v.0.8.2 (*85*), removed reads with mapping quality below 30 and removed duplicated reads with *sambamba* using *--remove-duplicates*, clipped overlapping reads using the *bamUtils clipOverlap* command (https://github.com/statgen/bamUtil), and indexed the bam files using *sambamba index*. Finally, we performed indel realignment per sample using *GATK3.8* with the *RealignerTargetCreator* and *IndelRealigner* commands. We estimated the depth of coverage for each bam file using *mosdepth* v.0.3.11 with the -n --fast-mode --by 500 --mapq 30 flags (*86*).

We performed population genomic analyses using genotype likelihoods in *ANGSD* v0.938 to account for genotype uncertainty (*87*, *88*). We identified variant sites across all individuals using *ANGSD* with the samtools genotype likelihood model (-GL 1), a minimum SNP value of 1e-6 (-SNP_pval 1e-6), determined major/minor alleles based on allele frequencies (-doMajorMinor 1), removed sites below and above a minimum and maximum total sequencing depth of 184 (number of individuals used to call SNPs) and 2500 (2 x total number of individuals x mean sequencing depth), respectively, across all individuals, and removed sites with missing data in more than 20% of individuals (-minInd 153), minimum base quality below 30 (-minQ 30) and a minimum individual depth of 1 (-setMinDepthInd 1). Furthermore, we removed multi-mapping reads (-uniqueOnly 1), unmapped or duplicated reads (-remove_bads 1) and those without a matching paired read (-only_proper_pairs 1). We also adjusted the mapping quality for excessive mismatches (-C 50). We only kept SNPs with a minor allele frequency of 5% (-minMaf 0.05). We then removed SNPs falling within potentially problematic genomic regions that can impact the analyses accuracy. We removed sites that fall within regions with low mappability (*89*). We estimated mappability using a kmer approach in *genmap* with 150-mers and we only kept SNPs within uniquely mapping regions (mappability of 1) (*90*). Furthermore, we removed sites with excess heterozygosity and all sites within 10kb of those sites. For that, we estimated inbreeding statistics using PCAngsd (*91*) and removed all sites (± 10kb) with F < -0.95 and P-value < 10−6 (*92*). This final, filtered SNP list was used as the basis for all further analyses across all samples.

### Mitochondrial variant calling and analysis

Genotypes were called for all whole genome reads mapping to the scaffold representing the mitochondrial genome across all individuals using *FreeBayes* (*93*) with ploidy set to 1. Subsequently, we filtered the resulting dataset using *vcftools* to keep all variant sites that were genotyped in more than 90% of all individuals, with mapping and genotyping quality above 30, and with minimum coverage of 5x and a maximum coverage of 100x. We reformatted the resulting VCF into nexus format using the *vcf2phylip* script (https://github.com/edgardomortiz/vcf2phylip) and generated a median joining haplotype network using *PopArt* (*94*).

#### Population genomics based on genotype-likelihoods

##### Population structure and phylogenetics

We investigated population structure across all populations using Principal components analyses with *PCAngsd* based on all SNPs and based on a thinned SNP dataset (1 SNP every 50kb; informed by decay of linkage-disequilibrium [LD]) with highly-divergent regions on chromosome 1 and 56 removed (thinned SNP dataset). We also inferred genetic ancestry using an admixture approach in *PCAngsd* for K=2 to K=12. We evaluated the most likely K using correlations matrices of residuals in evalAdmix (*95*). We also estimated population structure for the Oir and the Clyde catchment (Endrick, Ross Burn, Gryfe), separately, using *PCAngsd*, only using sites with minor allele frequencies above 5% in these subsampled datasets. Moreover, we also estimated PCAs for parasitism-associated regions using *PCAngsd*.

Furthermore, we inferred phylogenetic relationships between individuals and populations using several ways. First, we build neighbour-joining (NJ) trees using the bioNJ command in the *ape* v.5.8-1 R-package (R version 4.3.0 (2023-04-21)) based on a genetic distance matrix (for all SNPs and the thinned SNP dataset), which was inferred using *ANGSD* (--doIBS). Second, we estimated population relationships and inferred gene flow events between populations using *TreeMix* in the *BITE* V2 R-package (*96*, *97*). We first estimated minor allele frequencies for each population (‘ecotype by location’) using *ANGSD*, converted them into minor allele counts by multiplying the maximum likelihood sample frequency by the SNP-specific sample size, and created a minor allele count matrix across all SNPs and populations as input for *TreeMix*. We ran *TreeMix* with the single *L. auremensis* individual as the outgroup in the *BITE* R-package for 0 to 5 migration edges. First, *BITE* builds a phylip consensus tree with 500 bootstraps, and then models migration events along this tree. We analysed SNPs in blocks of 1000 SNPs to account for linkage between SNPs. We plotted trees with *BITE* for 1 to 3 migration events, and determined the most likely tree based on the Evanno method in OptM (*98*) and residuals plots. Second,, as the phylogenetic relationship between populations likely differs across the genome due to selection, gene flow and incomplete lineage sorting, we performed topology weighting analyses in *TWISST* (*30*) across the genome for i) the anadromous-parasitic and non-parasitic ecotypes from Oir and Endrick and ii) all three ecotypes from the Endrick together with the non-parasitic Ross Burn population. Due to the use of genotype likelihoods in our analyses to account for genotype uncertainty, we implemented a custom pipeline for *TWISST* (*89*). First, we estimated the IBS matrix per 50kb window (based on LD decay, below) using single read sampling (-doIBS 1) in *ANGSD* and then generated one NJ tree per window using the *bioNJ* command in *ape*. The weightings across all subtrees were calculated using the ‘complete’ method in *TWISST*.

##### Linkage disequilibrium (LD)

We inferred LD per ecotype by population using *ngsLD v1.1.1* (*99*). We created two different LD datasets. First, to identify windows of high LD across the genome, we ran *ngsLD* on all sites per ecotype by population and estimated LD between sites that were maximum 1kb apart to reduce computational burden. For plotting, we summarised LD estimates in 10kb sliding windows (2.5kb steps) using the *WindowScanR* R-package, and plotted the loess-smoothed values (span=0.1) across the genome using ggplot2. Second, to estimate LD decay rates, we only kept sites on chromosome 2, 5 and 10, and estimated LD up to a maximum distance of 100kb between sites for each ecotype by river using *ngsLD*. LD decay was plotted using custom scripts in R.

##### Genome-wide association study for life-history using GEMMA

To map the genomic basis of life-history, we used a binary genome-wide association analysis (GWAS) in *GEMMA* (*100*). We performed the GWAS for life-history based on mean genotypes to account for genotype uncertainty. Mean genotypes were inferred as the mean of scaled genotype probabilities per SNP using custom scripts and genotype probabilities stored in beagle format (generated with *ANGSD* using the -GL 3 command). Mean genotypes in BIMBAM format were used as input for *GEMMA* to first generate a covariance matrix and subsequently test for genetic associations with life-history coded as 0 for parasitic and 1 for non-parasitic using linear mixed models with the Wald test (-lmm 1). We performed GWAS for all individuals in the dataset and only for individuals from the Endrick catchment to reduce the potential impact of population structure.

##### GO term enrichment analysis

We performed the functional annotation based on annotated genes (REFseq annotation for kcLamFluv1.1 from NCBI) and gene ontology terms provided with the REFseq annotation (.gaf file). using *clusterProfiler* version 4.8.2 (*101*) in R. In *clusterProfiler*, we performed gene ontology overrepresentation analyses (ORA) for genes containing significant parasitism-associated SNPs, and gene set enrichment analyses (GSEA) with genes ranked by the highest p-value (from parasitism GWAS) within each gene that contained a SNP. We did not perform a correction for multiple testing due to the hierarchical structure of GO terms. However, we summarised the results based on semantic similarity using the *simplifyEnrichment* R-package (*102*) and plotted the top 25 GO terms for each test using *clusterProfiler*.

##### Genetic differentiation and selection

We estimated genetic differentiation between ecotypes within rivers using *ANGSD* based on all filtered SNPs. We estimated the 2D site frequency spectrum for each comparison using *realSFS* and used *realSFS fst* to estimate Fst for each comparison. We summarised Fst values in 10kb sliding windows (5kb steps) using the *realSFS fst stats2* command, and plotted the results as manhattan plots using the *CMplot* R-package and ggplot2.

To estimate ecotype specific selection, we estimated the population branch excess statistic (PBE), an extension of the population branch statistic (PBS), using the approach outlined in Shpak et al. (*103*). PBS aims to identify selective sweep signatures by comparing the focal branch length of a locus within a population to two outgroup populations, and the PBE statistic extends this approach and estimates selective sweeps by testing if the PBS value at a site exceeds the expected value under neutrality. PBE for each locus (i.e. Fst for 10kb sliding window) in the focal population A was estimated as: PBE_A_ = PBS_A_ – PBS_A-exp_ = PBS_A_ – [(T_BC_ × med(PBS_A_)) / med(T_BC_)], with med(PBS_A_) and med(T_BC_) being the median values of their respective statistics across all loci. PBS_A_ = (T_AB_ + T_AC_ - T_BC_) / 2, with T being the linearised Fst between populations at this locus (T = -log(1-Fst). Fst values were mean Fst values per 10kb sliding window as estimated above, and for estimates in the Endrick system we used the three ecotypes (ewAP, ewNP, ewLP), changing the focal population to estimate population-specific PBE values. For the Oir, we used the two ecotypes and the lake-parasitic ecotype from the Endrick as an outgroup. We used these estimates to compare PBE values for genomic windows containing parasitism associated SNPs between ecotypes by river using Wilcoxon rank sum tests.

#### Population genomics of Occidentis ayresii

To test if the genomic basis of parasitism loss is convergent or divergent across lamprey lineages, we re-analysed low-coverage whole genome sequencing data (mean coverage depth = 0.51x, range = 0.27x to 1.26x) for sympatric parasitic Western River Lamprey and the non-parasitic Western Brook Lamprey ecotypes of *Occidentis ayresii* (formerly *Lampetra ayresii* and *L. richardsoni,* respectively) (*33*). *Occidentis ayresii* is a North American sister lineage to *Lampetra*. These lineages have diverged approximately 5-10MYA (*17*, *33*). As no highly contiguous reference genome was available for *O. ayresii*, we performed synteny-guided scaffolding of a fragmented draft genome (*33*) against the DToL *Lampetra fluviatilis* reference genome (kcLamFluv1.1) using *RagTag* (*104*) with default settings, anchoring ∼882Mb (∼87% of the genome) into chromosomes. We also lifted over the annotation from kcLamFluv1.1 to the scaffolded reference using *Lifton* with default settings (*105*). We identified variant sites using *ANGSD*, retaining sites using the same filtering strategy as for European lamprey (see above), changing the depth and minimum individual filters. We retained sites with a minimum mean depth above 39 and below 100, and present in at least 18 individuals. We generated a genetic distance matrix based on single read sampling (-doIBS 1) in *ANGSD* and used multidimensional scaling to confirm the population structure. Furthermore, we estimated per-SNP Fst between ecotypes with *ANGSD* and *realSFS* as described for European lamprey. Per SNP Fst estimates were plotted using CMplot. Lastly, we performed principal components analysis for the main outlier regions (based on elevated Fst) between ecotypes, with *PCAngsd* to identify putative signals of structural variation.

#### Structural variant analysis

##### Genome assemblies

For the analyses of chromosomal rearrangements, we used newly, publicly available chromosome-scale reference genome assemblies for the anadromous-parasitic European river lamprey (kcLamFluv1.1 primary assembly from the Darwin Tree of Life [NCBI] and *kcLamFluv2.1* haplotype 1 and 2 from the Norwegian BioGenome Project (*49*)) and European brook lamprey (kcLamPlan1.2 haplotype 1 and 2 from the Norwegian BioGenome Project (*49*)). These haplotype-resolved assemblies from the Norwegian BioGenome Project are earlier versions of the ones available on NCBI (kcLamFLuv2.2 h1 and kcLamPlan2.2 h2), but are comparable and just differ in scaffold naming. Furthermore, we assembled haplotype-resolved and chromosome-scale genomes for one European river lamprey and one European brook lamprey from the Oir using a combination of PacBio Hifi and Hi-C (*49*) (see Table S3 for assembly details). For genome assemblies, fresh milt (sperm) was collected from spawning-ripe males caught in the Oir, immediately flash-frozen in liquid nitrogen, and stored at -80°C until further processing. High-molecular weight (HMW) DNA was extracted from 100µl of milt using the Nanobind DNA extraction kit for tissue, following the providers manual, and quality control performed using a Nanodrop Spectrophotometer, Agarose gel, Qubit and Agilent Tapestation with the Genomic DNA tape. HMW DNA with an average fragment length above 60kb was sent for library preparation and sequencing on one PacBio Revio SMRT cell per sample at Edinburgh Genomics. Hi-C libraries were prepared from 100µl of milt from the same individual using the PhaseGenomics Proximo Hi-C kit following the provider’s manual. Final Hi-C libraries were sequenced to 100M reads per library on a NovaSeq X Plus 10B flow cell (together with other samples). We performed genome assemblies using established pipelines for European lampreys (*49*). First, we filtered PacBio Hifi reads using HiFiAdapterFilt-3.0.0 and assembled haplotype-resolved contigs from filtered PacBio Hifi reads using *hifiasm v.0.19.7-r598* (*106*) with the Hi-C option for phasing, and everything else at default settings. Second, we processed Proximo Hi-C reads following provider’s recommendations, aligned reads to the contigs using *bwa mem* using the -5SP setting, and performed quality control of Hi-C alignments using the *hic_qc.py* script provided by PhaseGenomics (https://github.com/phasegenomics/hic_qc). After marking duplicates with *sambamba*, we scaffolded the haplotype-specific assemblies using *YAHS* (*107*) with a mapping quality threshold of 10 (-q 10) and default settings. We visualised scaffolded assemblies using *PretextView* (*108*). Lastly, we improved the scaffolded haplotype-resolved assemblies through an additional round of reference-guided scaffolding using *RagTag* v2.1.0 (*104*). The brook lamprey assembly was scaffolded against LamPlan1.1_h2 and the river lamprey assembly was scaffolded against the reference genome (kcLamFluv1.1).

##### Chromosomal rearrangement identification

We used two different approaches to identify chromosomal rearrangements between European lamprey ecotypes (see above for details on all assemblies) and across lamprey lineages. We used chromosome-scale assemblies for the parasitic *Petromyzon marinus* (UKy_Petmar_22M1.pri1.0 GCF_048934315.1) (*51*) and non-parasitic *Lethenteron reissneri* (ASM1570882v1 - GCF_015708825.1) (*50*) as outgroup species.

First, we investigated synteny and identified large-scale chromosomal rearrangements across different combinations of genome assemblies using *ntSynt* (*109*) using default settings and visualised results using *ntSynt-viz* (*110*).

Second, we identified rearrangements between *Lampetra spp.* assemblies using *SyRi* (*111*). As *SyRi* requires genomes to have the same sequence names, we scaffolded all genomes for each comparison against a common reference (kcLamFluv1.1) using *RagTag* v2.1.0 (*104*) and scaffolds renamed following the reference genome. Re-scaffolded assemblies were aligned against each other using *minimap2* with the -ax asm5 -t 8 --eqx settings (*112*), and pairwise alignments used as input for *SyRi* using default settings. SyRi results were plotted using *plotsr* (*113*).

### Copy-number variation

We inferred copy number variation (CNV) between ecotypes from normalised read depth following the approach outlined in (*114*). First, we estimated read depths in 10kb windows across the genome for each individual using *mosdepth* (*86*) for reads with mapping qualities above 30. Next, we normalised read depths by dividing the estimated read depth for each window by the genome-wide median using custom R scripts, and identified putative CNVs as windows with normalised read depth 2 standard deviations above or below the mean. Furthermore, to identify genomic windows that differ in normalised read depth between sympatric ecotypes, indicative of divergence CNVs, we estimated Qst values for each window between sympatric ecotypes using a linear mixed model approach outlined in (*114*), which adapted the Qst and population variation approach outlined in (*115*). In brief, we performed linear mixed models in the lme4 R-package to extract within and among population variance based on unnormalised read depths, correcting for variation in average read depth by including the per-sample median read depth as a covariate, and population/ecotype as a random factor. The variance component attributed to the population was used as the among-population variance (V_A, among_) and the model’s residual variance component was used as the within-population variance (V_A, within_), which were used to estimate Qst using the following formula: Qst = V_A, among_ / (V_A, among_ + 2 * V_A, within_). We conservatively identified outlier CNVs as those above the 1% genome-wide Fst distribution following a Qst - Fst approach (*114*), and inspected the normalised read depth distribution around putative outlier CNVs overlapping parasitism-QTL.

### Recombination rate estimation

To estimate recombination rate variation across the lamprey genome for specific ecotypes, we used different complementary approaches.

First, we used allele frequency data and machine learning approaches to estimate population-scale recombination rates for each ecotype in the Oir and Endrick population from whole-genome sequencing data using *ReLERNN* (*116*). We estimated allele frequencies per ecotype and population using *ANGSD* for all filtered SNPs, and ran *ReLERNN* using default settings with allele frequencies, which has been shown to be accurate for reconstructing recombination rate landscapes (*117*).

Second, we reconstructed linkage maps for a hybrid family based on RADseq data to investigate recombination patterns. To construct our linkage map, we used the RAD sequencing data generated for a single cross of parent caught in the Oir (*25*). We first demultiplexed the raw data using the program process_radtags of the *Stacks* software v2.4 (*118*, *119*). *process_radtags* was used to check reads overall quality, trimmed read to 85bp and removed reads containing Illumina adapters. The reads of each individual were aligned to the HiFi NP draft genome (earlier version) using BWA-mem (*84*), and the output transformed to bam format with samtools (*120*). We then ran *gstacks* (Stacks v2.4) in reference-based mode and removed the PCR duplicates via gstacks. We removed unpaired reads and a minimum PHRED-scaled mapping quality of 25 was set to consider a read. We also applied a maximum soft clipping level of 5% of the read length to reduce sequencing errors. The *populations* program (*Stacks*) was used to export the results in VCF format for further filtering and only keep informative individuals for linkage map construction: the 2 parents (anadromous-parasitic female and non-parasitic male from Oir population) and their 90 offspring. The SNPs at positions 60 were removed due to a sequencing problem occurring at this position (*25*). We filtered SNPs using *vcftools* (*121*) to remove low quality genotypes and SNPs. We kept genotypes with a (mean) coverage ranging from 5x to 100x. We only included bi-allelic sites with less than 60% missing data, and a minor allele count of one. Subsequently, we filtered out two individuals with a mean depth below 10 and more than 25% missing data. We checked for mendelian inheritance errors with *bcftools*, leading to the removal of one individual. We then repeated the SNP filtering as before but only kept SNPs with less than 30% missing data. We also sought and removed singletons with the --singletons option of *vcftools*.

We used *LepMap3* (*122*) to construct the linkage maps. Two different pipelines were tested to construct the map: with or without *Filtering2*. When using filtering, *Filtering2* was run with the following parameters: removeNonInformative=1 and dataTolerance=0.0001. Small dataTolerance like this is advised in single-family crosses. Then, for the SeparateChromosomes2 module we computed different linkage maps testing different lodLimit ranging from 6 to 12 and removed the LG computed with less than 20 SNPs with sizeLimit=20. When not using *Filtering2*, we ran *SeparateChromosomes2* the same way as before but with the parameter distorsion Lod=1 as advised for single family data. We expected approximately 82 chromosomes in *L. planeri* and *L. fluviatilis* (*123*). We chose to continue the analysis with the pipeline with filtering because without, the first LGs contained too many markers and very few were left in the others. The selected map was passed through the module *JoinSingles2All* with parameters lodLimit=6 and lodDifference=2.

Subsequently, we aligned RADtags containing SNPs to the kcLamFluv1.1 reference genome using *bwa mem* and ordered SNPs used for linkage mapping along the reference genome. We removed RADtags that did not uniquely mapped and filtered out SNPs, for the female and male map separately, that did not show a collinear relationship between physical and genetic distance, unless ten or more SNPs showed that pattern (e.g. inversions).

#### Individual-based simulations of hybridization

We aimed at comparing the observed proportions of admixture with simulated proportions computed according to various scenarios of mating and hybrid breakdown. We used the forward-in-time individual-based simulation software *NEMO* v.2.3.55 (*124*). We aimed at simulating a life cycle as close as possible as the one of lampreys.

We implemented two different models. First, we modelled assortative mating (i.e. within ecotype) based on a binary indicator phenotype (AP (1) or NP (0)) and tested different levels of random mating (i.e. possibly between ecotypes): 10%, 30% or 50%. Second, we implemented a model of hybrid incompatibility where hybrid survival depended on their pedigree, with overdominance expressed in the F1 hybrids and underdominance (hybrid breakdown) expressed in the F2, backcrosses and later hybrid crosses. Individual fitness (offspring survival) was determined by ten pairs of Dobzhansky Müller Incompatibilities (DMI) loci, with free recombination. NP and AP parental types were initialised with opposite alleles (0/0 in AP, 1/1 in NP), facilitating the distinction between F1 hybrids (all 0/1 or 1/0 genotypes at all loci) from F2 and later generation crosses (mixed 0/1 and 1/0 genotypes). Genotype fitness were set such that survival rates were 0.82 for pure-type offspring, 1 for F1’s, and a minimum of 0.11 for 100% incompatible heterozygotes in F2’s.e. Given the different body size of ecotypes, the fecundity of parasitic lamprey (AP) was multiplied by 3 or 5 during breeding. The phenotype of the offspring (parasitic lamprey or non-parasitic lamprey) was inherited by ancestry percentage and generations were non-overlapping. We simulated three populations: one sink population mimicking a hybrid zone receiving a unidirectional migration from two other populations containing either non-parasitic or parasitic individuals (fig. S29A). At the first generation, the sink population contained only non-parasitic individuals. We recorded the individual genotypes at the DMI loci and their phenotypes in the sink population after 200 generations for further analyses. The populations’ carrying capacities were set at 1,000. We performed 50 replicates of each combination of parameters (dispersal, fecundity and assortative mating, fig. S29B). The classification of hybrids with the *NEMO* output was performed with a custom script.

#### Hi-Plex genotyping

To investigate the population structure and role of chr1 and chr56 in ecotype divergence, we performed targeted genotyping of over 1000 lampreys from a hybrid zone in the Oir using a Hi-Plex amplicon sequencing approach.

##### Target SNP selection

Hi-Plex is an amplicon sequencing technique allowing co-amplification of all SNPs in a single multiplex PCR prior to Illumina sequencing. Here we developed a set of 148 SNPs allowing the distinction of *L. planeri* and *L. fluviatilis* and including both loci with a high divergence between ecotypes, and randomly selected SNPs. We used RAD sequencing data (609 individuals distributed across 17 populations) from (*25*) for target SNP selection. Only individuals from French populations with a genome-wide Fst < 0.1 were retained for downstream analyses (n = 121 individuals). These populations experience high gene flow and are therefore more appropriate for investigating reproductive isolation than allopatric populations with low gene flow in which highly differentiated markers may reflect population-specific drift. To call SNPs, we first demultiplexed the raw data using *process_radtags* in *STACKS* v2.4 (*119*), and used process_radtags to check read quality, trim reads to 85bp and removed reads containing Illumina adapters. Reads were aligned to a *L. planeri* draft genome assembly (based on PacBio Hifi data; see (*125*) for details) using *BWA mem*, and transformed to a sorted .bam file with SAMtools. We ran the *gstacks* module in reference-based mode to call SNPs, after using it to remove PCR duplicates, unpaired reads and removed reads with a minimum PHRED-scaled mapping quality of 25. We also applied a maximum soft clipping level of 5% of the read length to reduce sequencing errors. We exported a VCF file for further processing using the *populations* module of *STACKS*. SNPs at position 60 were removed due to sequencing problems occurring at this position (*25*).

After data visualization, we retained genotypes with a mean coverage between 5× and 100× and a minimum minor allele count of 5. Only biallelic sites with a maximum of 50% missing genotypes were kept. Eight individuals with more than 40% missing data were removed. Sites with more than 20% missing data in either *L. planeri* or *L. fluviatilis* individuals were excluded. Finally, we retained biallelic sites with a minimum minor allele count of 5 that were genotyped in at least 85% of individuals. The filtered VCF file was converted into a GENEPOP file using *PGDspider* v2.1.1.5 (*126*). Fst was computed for both GENEPOP files using the function diffCalc from the package *diveRsity* (*127*). In total, 190 outlier loci with Fst > 0.3 between ecotypes were identified. To improve genome-wide representativeness, we added putatively neutral SNPs (hereafter “neutral SNPs”), as outlier SNPs were primarily located on chromosomes 1 and 56. Neutral SNPs were identified using the populations program (Stacks) with individuals from the Oir river. A minimum of 2% of individuals were required to present the locus, and a minimum minor allele count of 3 was required to process loci. The dataset was then split by ecotype, and genotypes with a mean coverage between 5× and 100× and a minimum minor allele count of 5 were retained. Only biallelic sites with a maximum of 50% missing data were included. Individuals with more than 60% missing data were removed. Using *VCFtools*, loci with more than 15% missing data, loci deviating from Hardy–Weinberg equilibrium, and loci with a minor allele frequency below 0.1 were excluded. The populations program was then used to filter out loci with heterozygosity below 0.5. Markers present in both *L. planeri* and *L. fluviatilis* individuals were retained.

##### Hi-Plex primer design

To design primers for the HiPlex assay, we extracted a 145bp sequence around each selected SNP from *L. planeri* draft genome contigs. These sequences were aligned against the *L. planeri* genome using BLASTn and only matches with an E-value < 1×10^-20^ were retained. A custom script parsed BLAST output and retained only sequences with a unique match. The original FASTA file was filtered accordingly, yielding 138 sequences for outlier SNPs and 9,105 sequences for neutral SNPs. Repeats were screened using Rad4SnpsRepbaseFilter.py, resulting in the removal of 9 outlier SNP sequences and 1,037 neutral SNP sequences. A custom script was then used to retain only non-overlapping SNPs, followed by filtering of non-sequenceable sequences. This resulted in 63 outlier SNP sequences and 5,120 neutral SNP sequences. Neutral SNP sequences were randomly subsampled to retain one sequence per contig, resulting in 585 neutral SNP sequences. The final set of 648 sequences was then used for primer design using the Hi-Plex protocol (*128*). Designed primers were blasted against the *L. planeri* draft genome contigs to remove repeated loci and nonspecific amplifications.

##### Hi-Plex genotyping

A total of 1,291 individuals (536 non-parasitic *L. planeri*, 480 parasitic L*. fluviatilis* and 275 *Lampetra* larvae of unknown ecotype) were sampled in the Oir River between 2010 to 2021 by electrofishing. DNA was extracted from fin clips using the NucleoSpin® 96 Tissue kit (Macherey-Nagel, Düren, Germany) following the manufacturer’s instructions. All individuals were genotyped for 148 SNPs using a modified version of the Hi-Plex protocol (*128–130*). Dual indexing was applied to each sample to allow demultiplexing. Reads were aligned to the *L. planeri* draft genome using *BWA mem*, and allele counts were obtained with *SAMtools* (*120*). For each SNP in every individual, genotypes were inferred from the observed allele counts obtained through sequencing by calculating genotype likelihoods under a multinomial probability model. The genotype exhibiting the highest posterior probability was assigned to the corresponding SNP–individual combination only if this probability exceeded a threshold of 70%; otherwise, the genotype was treated as missing. We only retained SNPs that were genotyped in more than 75% of individuals and retained individuals with less than 25% missing data. We re-aligned RADtags around target SNPs to the kcLamFluv1.1 reference genome used in this study using *BWA mem.* We removed ‘neutral SNPs’ that were located on chromosomes with parasitism-QTL, even if they were not located directly within these, to remove potentially impacts on neutral population structure through linkage. This left a total of 131 SNPs, with 9 SNPs within parasitism-QTL on chr1, 21 SNPs within *trans-inv56* and 101 non-divergent SNPs outside parasitism-QTL (‘neutral SNPs’).

##### Analyses of Hi-Plex genotyping data

To investigate the population structure of lampreys in the Oir hybrid zone, we performed principal components analyses and admixture analyses for the three different SNP subsets, SNPs on chr1, trans-inv56 and neutral SNPs. First, we performed PCA for each SNP set in *adegenet* and plotted them with *ggplot2*. For SNPs within *trans-inv56* we determined the karyotype of each individual using a k-means clustering approach based on PC1 and PC2, and estimated the karyotype frequencies for each ecotype and life stage based on these estimated karyotypes. Furthermore, to test for an excess of *trans-inv56* heterozygotes within each ecotype, we tested for deviations from Hardy-Weinberg Equilibrium within each ecotype using the *HWChisq* function in *HardyWeinberg* R-package. Furthermore, we estimated genetic ancestry proportions for each SNP dataset using *Admixture* (*131*) with a K of 2. Lastly, we used linear mixed models with *lmer* in the *lme4* R-package to test for correlations of genetic variation for each SNP subset (PC1 for chr1 and neutral SNPs, karyotype for chr56) with sperm concentration and sperm speed (VAP), which were measured for a subset of genotyped individuals (n=42) in a previous study (*59*). In brief, adult *L. fluviatilis* and *L. planeri* were caught electrofishing in the Oir, kept at the field station and anaesthetized with benzocaine before fin clipping and stripping. Fin clips were used for genotyping using the HiPlex protocol as described above. Sperm traits were measured within 5 to 30min post stripping. Sperm velocity was measured using a Computer-assisted Sperm Analysis system (*OpenCASA*) and sperm number was counted using a Thoma cell counting chamber on three subsamples per male (see (*59*) for details). We included ecotype as a random effect in the model.

#### RNAseq analysis

To identify differentially expressed genes between ecotypes across tissues, we performed RNA-seq on liver, gonad, gill and brain of spawning-ripe anadromous-parasitic and non-parasitic lamprey caught in the Oir. Eight parasitic *L. fluviatilis* adults (four males and four females) and eight non-parasitic *L. planeri* (four males and four females) were caught by electrofishing in the Oir River in April 2018, anesthetized, and then killed with a lethal dosage of benzocaine. We collected their liver, brain, gonad and gill (total = 32 samples) for the adults and immediately froze them in liquid nitrogen. Total RNA was extracted from brains and gonads with the NucleoSpin® RNA Plus XS extraction kit following manufacturer’s instructions (Macherey-Nagel), and other organs were extracted using TRIzol Reagent and chloroform phase separation. Individual Illumina Stranded TruSeq RNA libraries were 150bp paired-end RNA sequenced on a HiSeq3000 Illumina sequencer to an average of 3×10^7^ reads per sample.

Raw RNAseq data were aligned to the kcLamFluv1.1 reference genome using a 2-pass mapping approach in *STAR* (*132*). We created read count tables against transcript annotation features for the REFseq annotation using *featurecounts* (*133*), only using uniquely mapping reads. For the differential gene expression analyses we removed female gonad samples, leaving 40 samples. We performed gene expression analyses across all individuals and tissues using *DESeq2* (*134*), retaining all genes with a minimum of 10 counts per million (∼1 cpm per individual) in at least 3 individuals, and the following model: ∼ sex + group; with group being a combination of ecotype and tissue. PCA was performed based on vst-transformed data in *DESeq2*. We tested for differential expression between ecotypes by tissue, identifying genes as differentially expressed with an FDR < 0.05. We identified differentially expressed genes that overlapped with genes under parasitism-QTL using *bedtoolsr* intersect (*135*).

