## Supplementary Material for "A mosaic of genomic architectures underpins parasitism loss in a jawless vertebrate"

### Supplementary Tables

**Table S1:** Summary of populations and individuals used for whole-genome resequencing.

| Species | Location | N. Individuals | Mean coverage depth ± s.d. |
|---|---|---|---|
| *Lampetra planeri* | Oir | 20 | 6.7 ± 3.0 |
| | Endrick | 26 | 8.1 ± 3.0 |
| | Gryfe | 5 | 4.9 ± 0.5 |
| | Norway | 18 | 13.4 ± 4.5 |
| | Ross Burn | 15 | 7.4 ± 2.9 |
| | Tweed | 8 | 8.8 ± 4.4 |
| | Danube | 6 | 5.4 ± 1.8 |
| *Lampetra fluviatilis* | Oir | 20 | 5.4 ± 1.4 |
| | Endrick | 24 | 8.5 ± 3.5 |
| | Gryfe | 2 | 5.5 ± 0.31 |
| | Norway | 18 | 14.1 ± 2.8 |
| *Lampetra fluviatilis (lake-parasitic)* | Endrick | 27 | 7.3 ± 2.3 |
| Total | | 189 | 8.4 ± 4.1 |

**Table S2 -** Genes within *trans-inv56* with evidence for impact on sperm speed or motility.

| Gene | Biological Role | Key Reference(s) |
|---|---|---|
| ***CUL4B*** | **E3 ubiquitin ligase; spermatogenesis** | (Lin et al. 2016) |
| ***MFF*** | **Mitochondrial fission factor; midpiece assembly** | (Varuzhanyan et al. 2021) |
| ***CPT2*** | **Carnitine shuttle; mitochondrial fatty acid β-oxidation** | (Zhi et al. 2025) |
| ***IRS1*** | **Insulin receptor signalling; AKT pathway in sperm** | (Aitken et al. 2021; Castiglione et al. 2023; Neirijnck et al. 2019) |
| ***ATP1B4*** | **Na,K-ATPase β4 subunit; ion transport in sperm** | (Jimenez et al. 2011) |
| ***EFHC2*** | **Axonemal microtubule inner protein (MIP); motile cilia** | (Hwang et al. 2019) |
| ***SUCLA2*** | **TCA cycle (succinyl-CoA ligase); mitochondrial ATP** | (Woodhouse et al. 2022) |
| ***PRKX*** | **cAMP-dependent protein kinase (PKA family); sperm capacitation** | (Nolan et al. 2004) |
| ***PFN1*** | **Actin dynamics (profilin 1); spermatid cytoskeleton** | (Dai et al. 2015) |

**Supplementary file 1**

A) Assembly statistics for all chromosome assemblies
B) Linkage Map information
C) Genes under parasitism-QTL
D) Gene ontology enrichment results

### Supplementary Figures

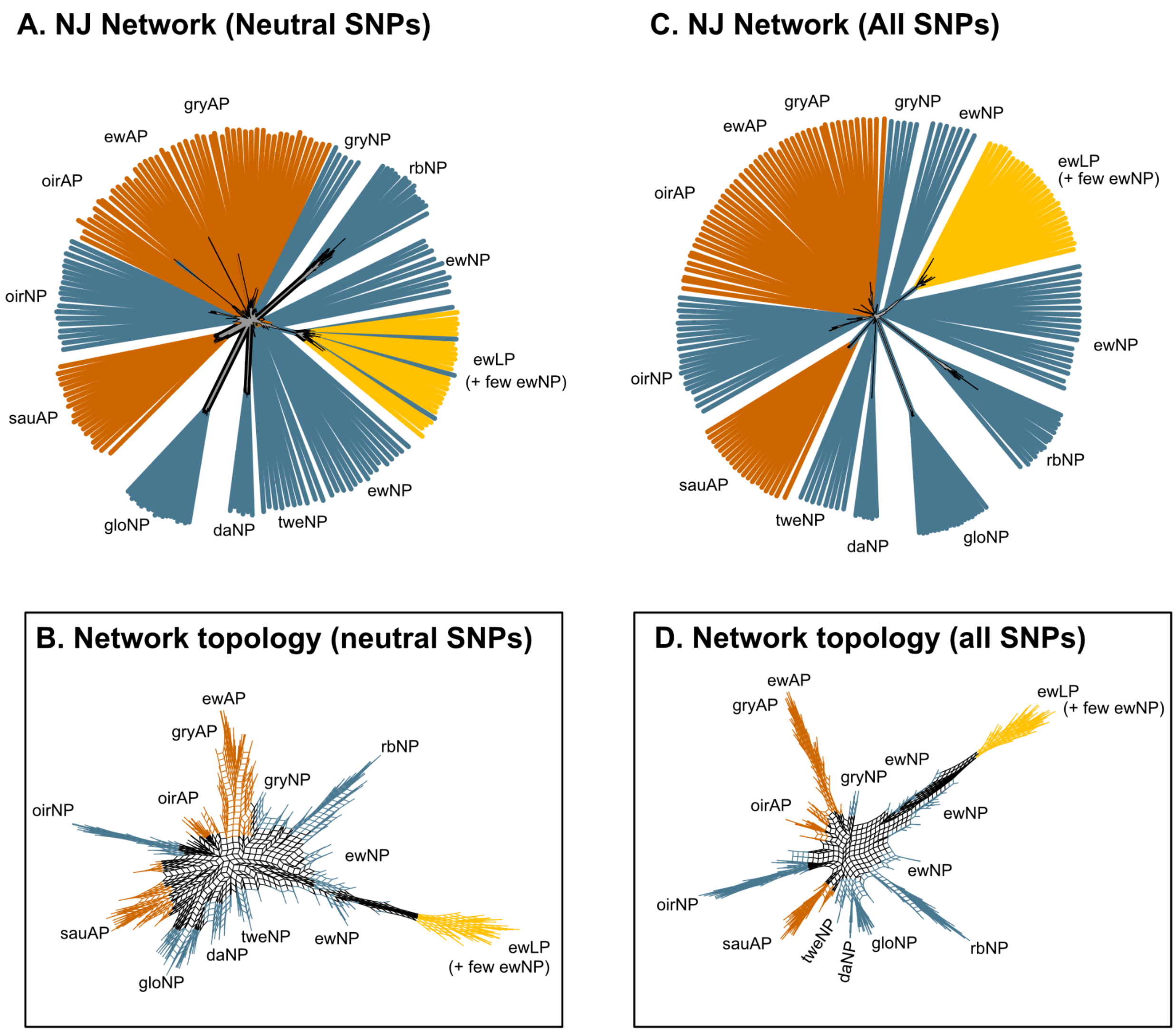


**Figure S1. Neighbour Networks.** Splitstree networks based on **(A)** thinned (1 SNP/50kb), non-divergent SNPs (without SNPs in high LD-regions on chr1 and chr56), with (B) showing the topology of splits in the network (zoomed in on base in A). **(C)** Splitstree NeighbourNet network based on all SNPs in the genome, with (D) showing the network topology at the base of the network (zoomed in on base in C).

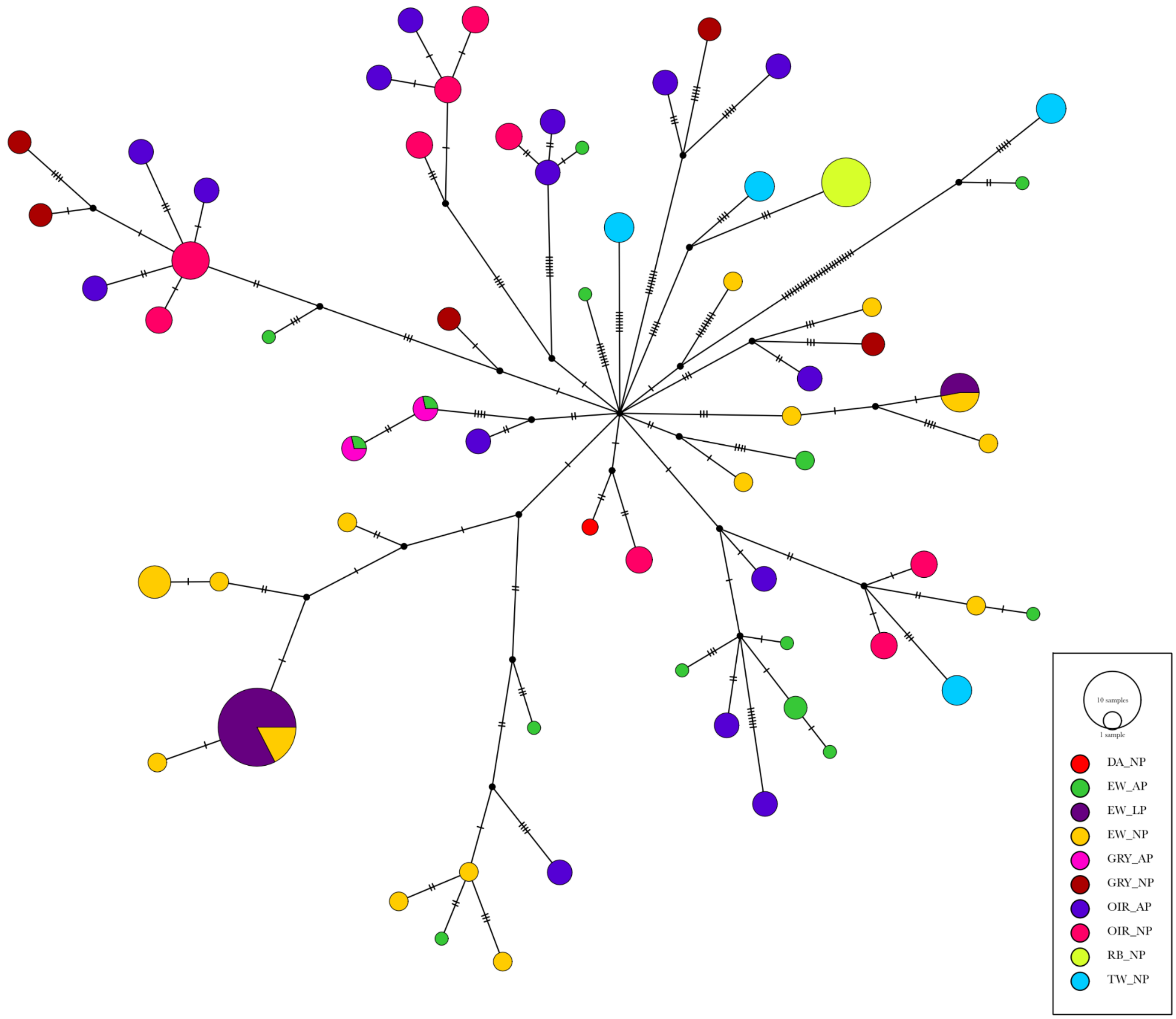


**Figure S2. Mitogenome Haplotype Network.** Median joining haplotype network based on 263 variant sites across the full mitochondrial genome. Circles represent observed haplotypes, coloured by population and ecotype and with the size representing the number of individuals sharing a haplotype (See legend). Cross bars between haplotypes represent the number of mutations separating two haplotypes. Note that haplotypes are not population and ecotype specific.

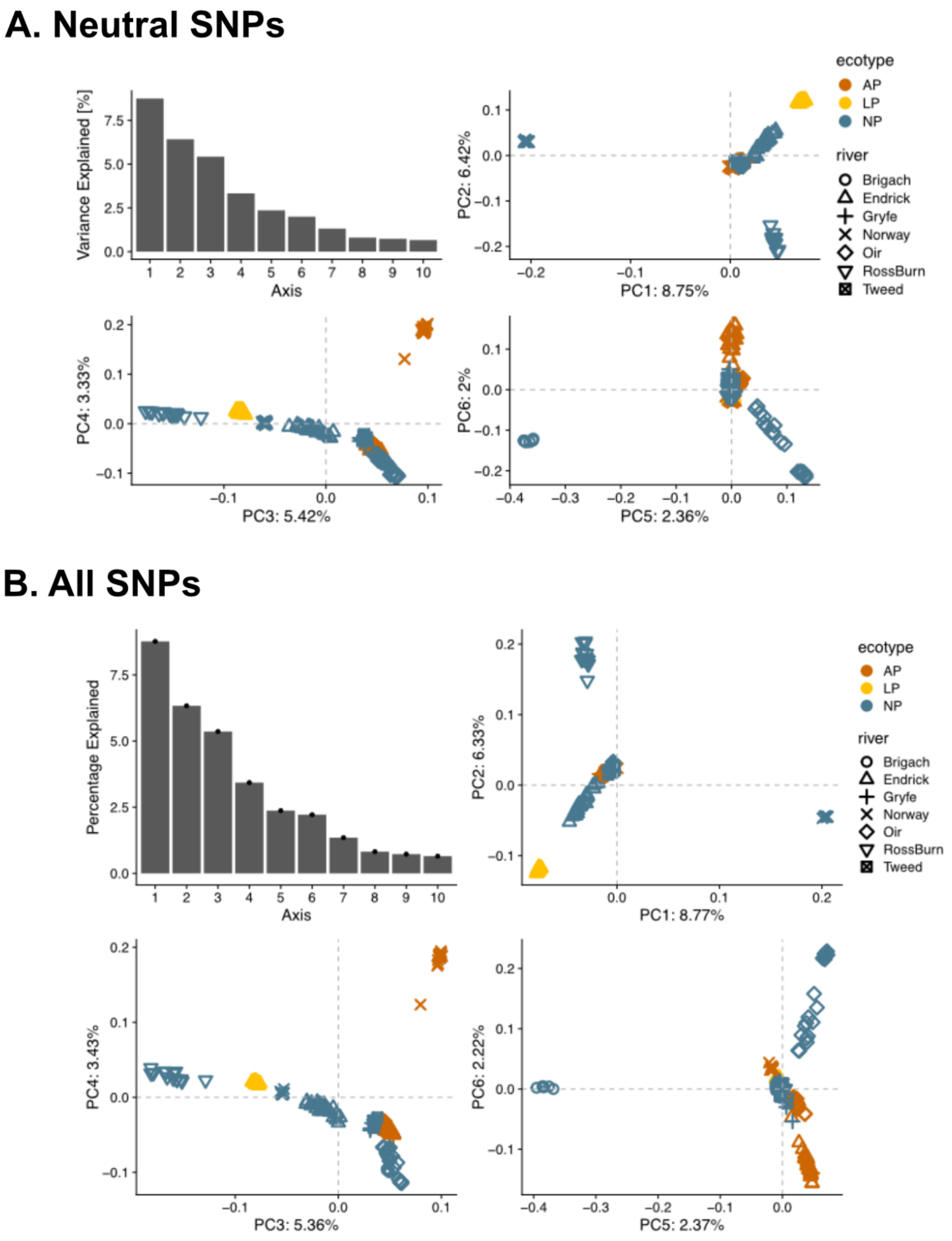


**Figure S3.** Principal components analysis. (A) PCA and genetic ancestry for all populations based on a thinned SNP dataset without highly divergent, high-LD regions on chr1 and chr56. The barplot shows the variance explained by each PC axis from 1 to 10. (B) PCA for all populations based on all SNPs. Colours and shapes are described in the legend.

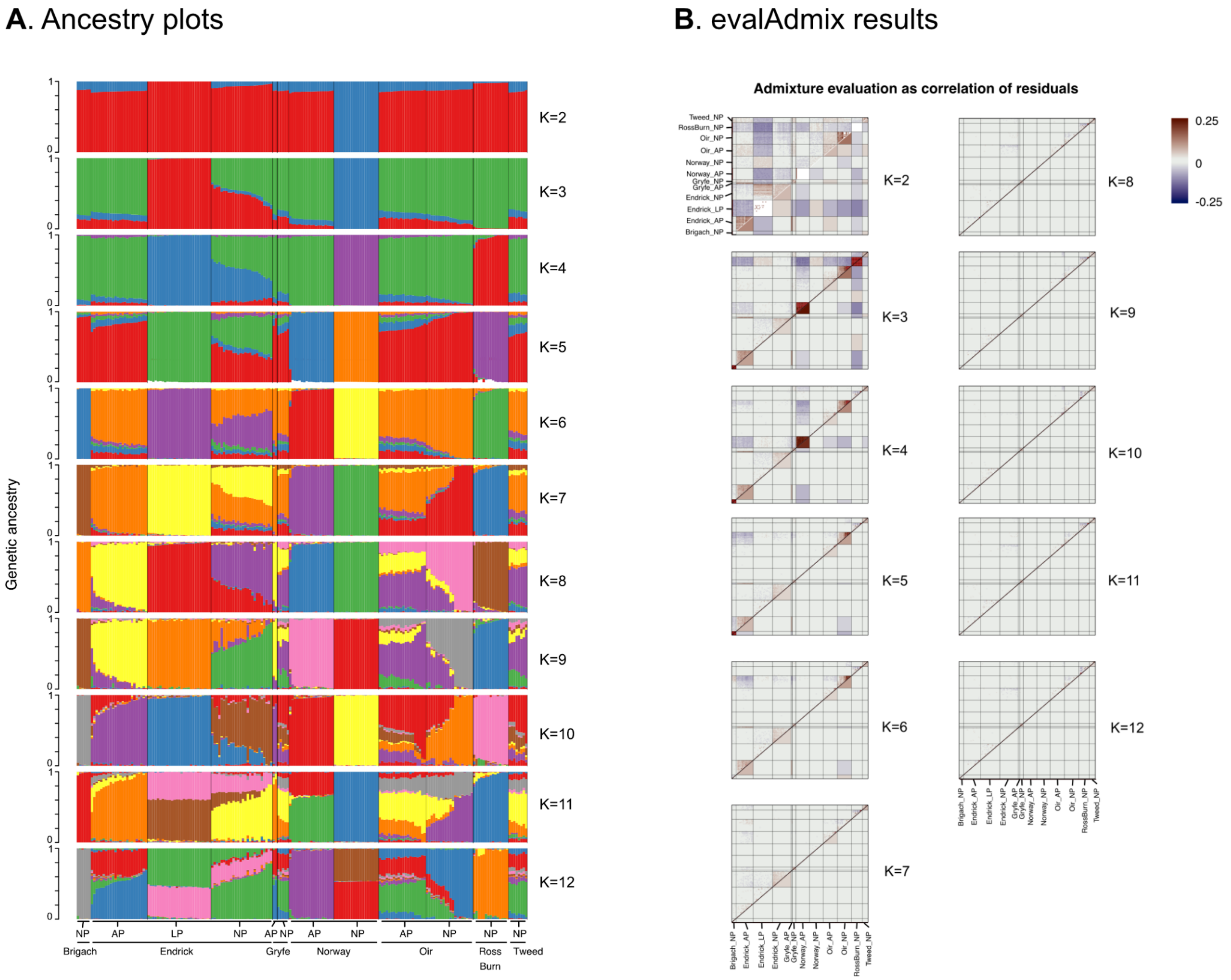


**Figure S4. Admixture results**. (**A**) Ancestry proportions inferred for all individuals based on thinned, non-divergent SNPs with PCAangsd. Shown are results for values of K from 2 to 12. (**B**) Shown are correlations of residuals for each pairwise comparison of individuals. Correlations were inferred with evalAdmix for ancestry proportions in (A), and were used to select the value of K at which correlations of residuals are minimised (K=8). Cells above the diagonal are individual-specific, and cells below the diagonal are averaged by population. Population and ecotype labels are shown next to each group. AP = Anadromous-parasitic; NP = non-parasitic; LP = lake-parasitic.

### **A**. Population structure in Loch Lomond system.

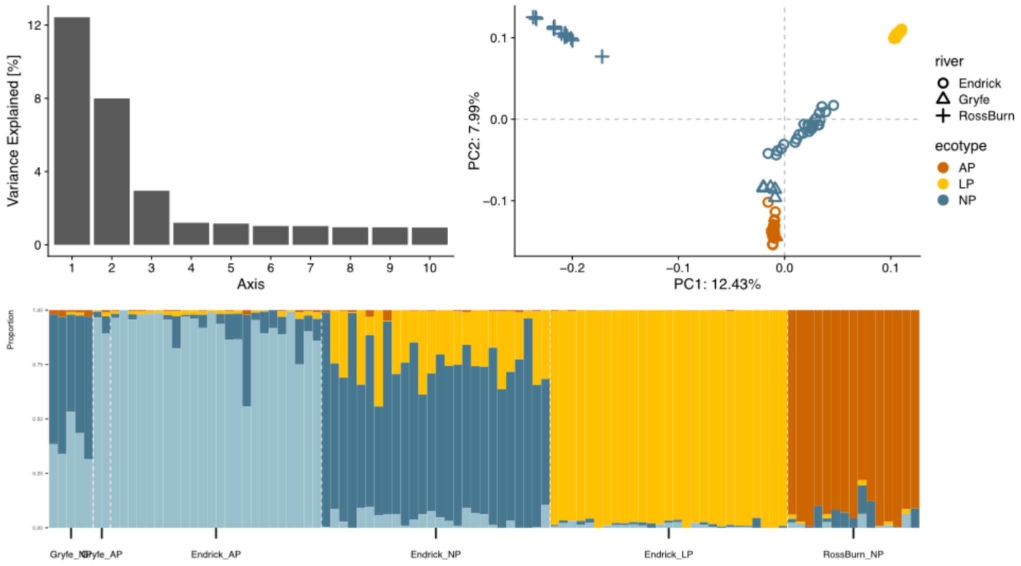


### **B**. Population structure in the Oir.

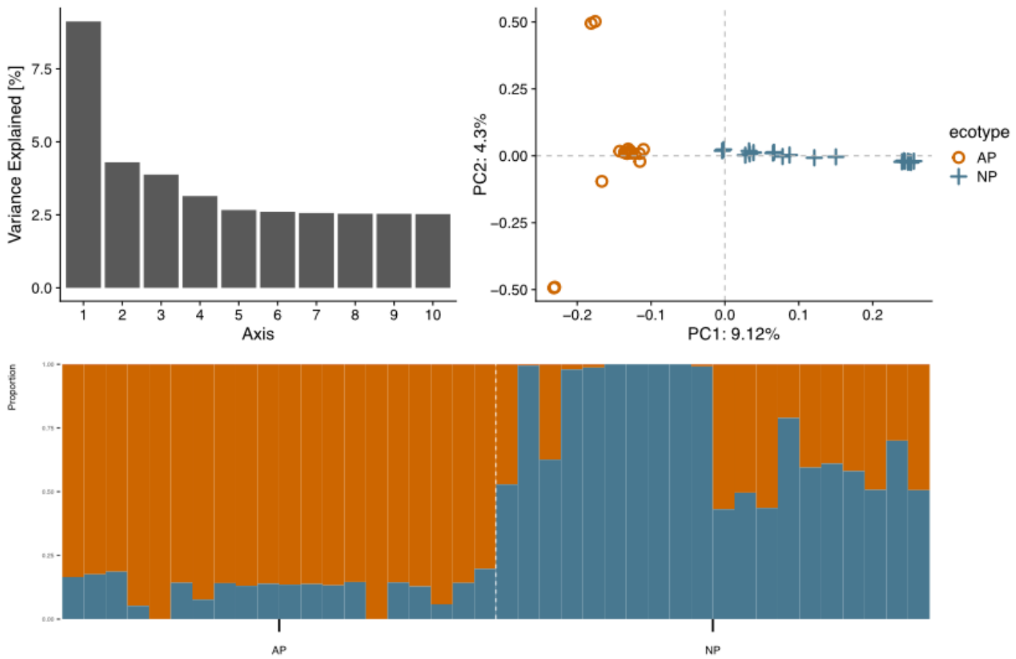


**Figure S5. Population structure by population.** (**A**) PCA and ancestry proportions based on all SNPs for individuals from the Loch Lomond catchment in Scotland (Endrick, RossBurn) and surrounding populations (Gryfe). The barplot shows the variance explained by each PC axis. (B) PCA and ancestry for all individuals from the Oir based on all filtered SNPs. Colours and labels are explained in the respective legend.

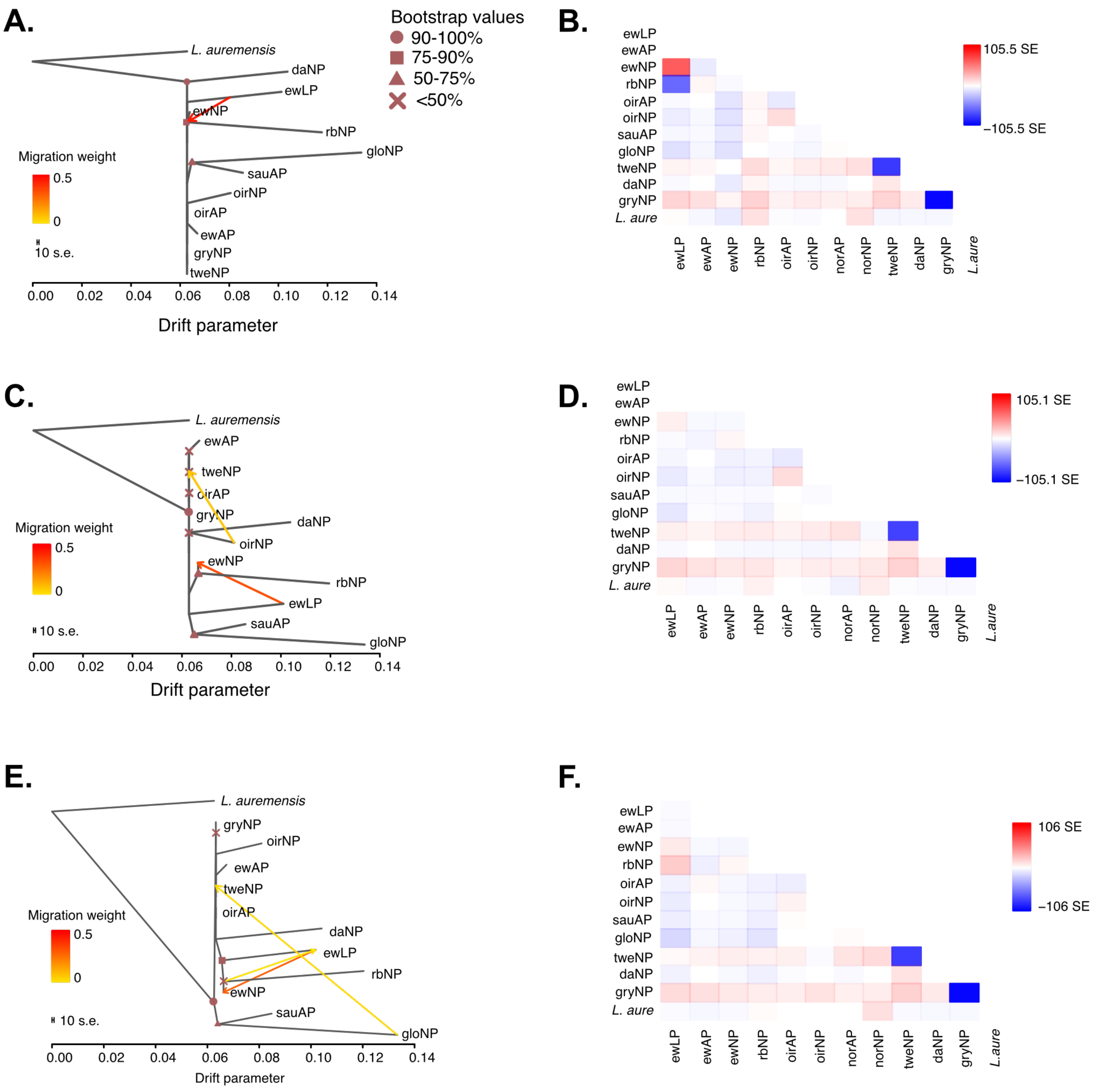


**Figure S6 -** TreeMix results showing the maximum likelihood based trees for neutral SNPs with different numbers of fitted migration events, m=1 (**A**), m=2 (**C**) and m=3 (**E**). Plots (**B, D, F**) show the residuals for respective ML trees.

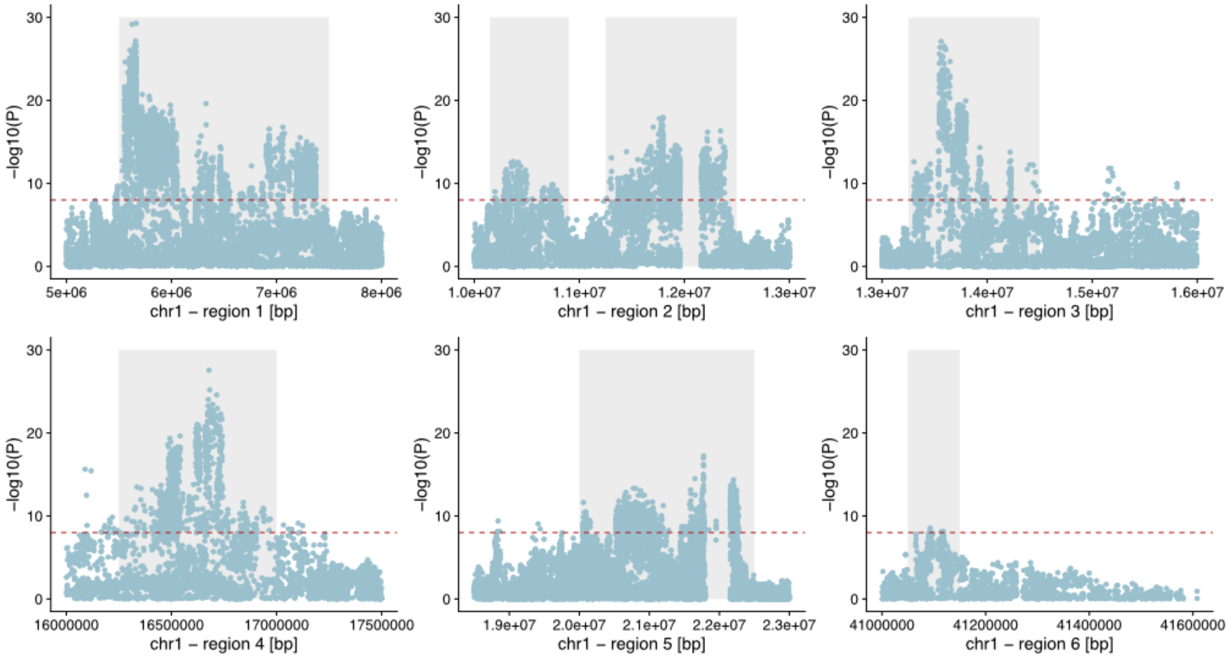


**Figure S7 - Genome-wide association with life-history (parasitism-QTL)** on chromosome 1. The plots show association results for each SNP on chromosome 1 by sub-region with distinct peaks. The dashed red line shows the genome-wide significance threshold. The grey bars highlight distinct association peaks.

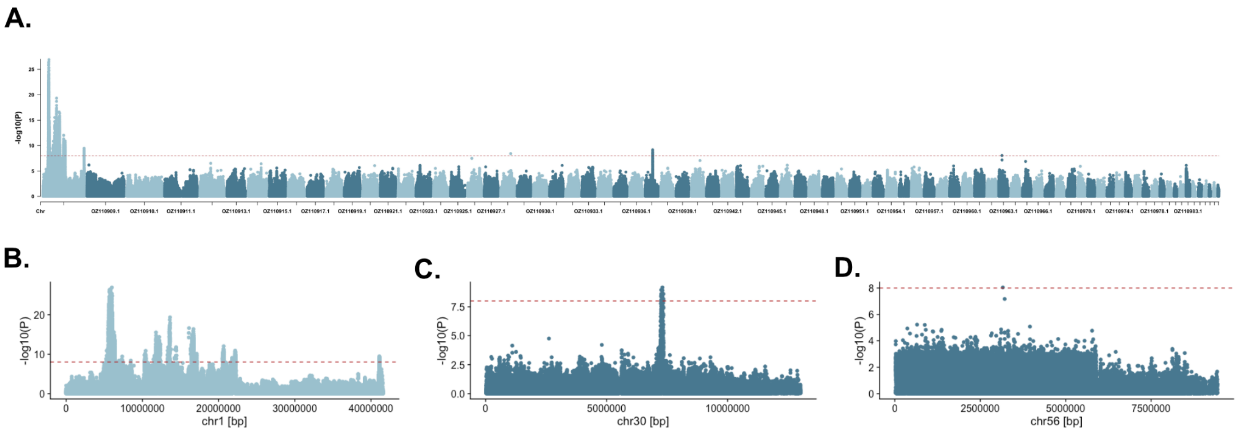


**Figure S8 GWAS for life-history in Endrick**. (A) Manhattan plot showing genome-wide association results from GEMMA for association with life-history in the Endrick population. (B-D) Each plots shows the association results for each SNP on chromosomes with significant signals on chr1, chr30 and chr56. The dashed red line shows the genome-wide significance threshold.

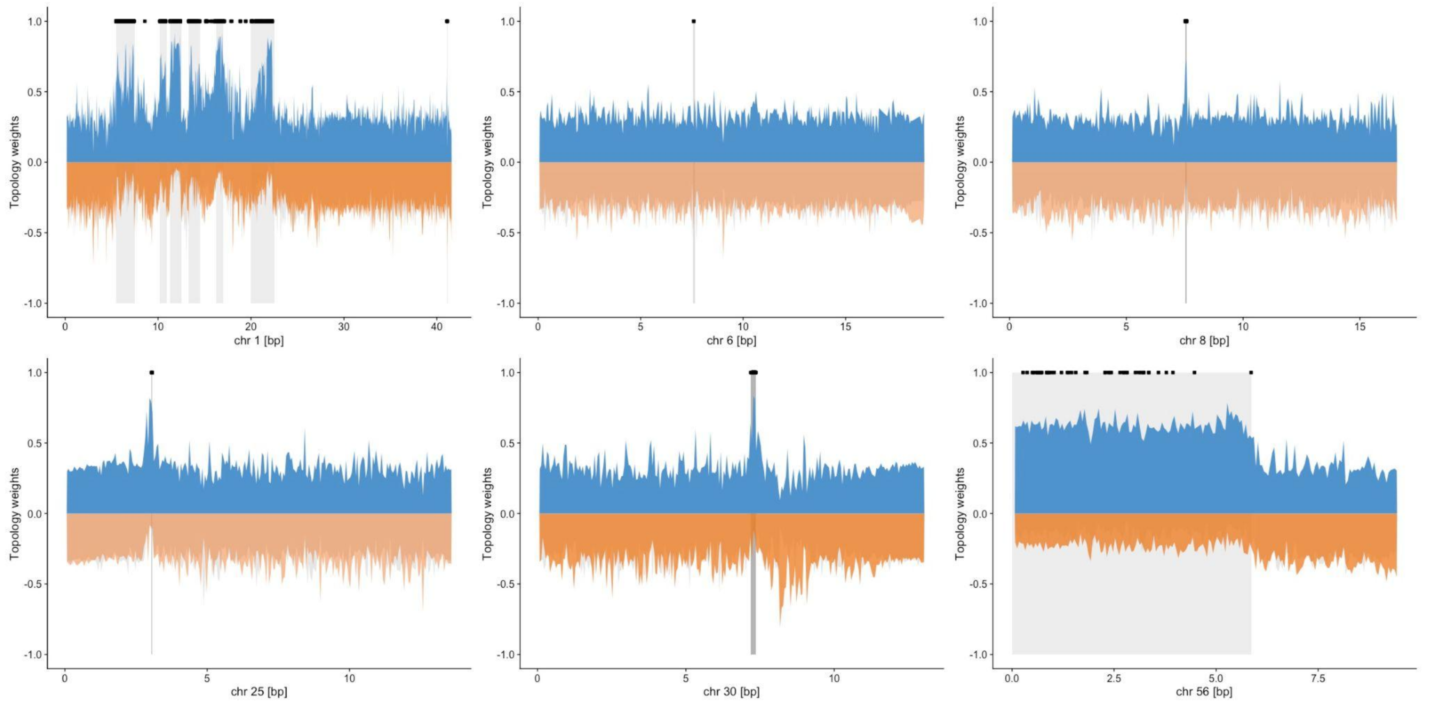


**Figure S9. TWISST results** for the anadromous-parasitic and non-parasitic ecotypes from the Endrick and Oir for chromosomes with parasitism-QTL. Positive values in blue show the weighting for the life-history topology, clustering individuals by ecotype, and negative values show the weightings for the geography topology, clustering individuals by river of origin, and the control topology. Black dots show all parasitism-associated SNPs and grey bars highlight the broader peaks.

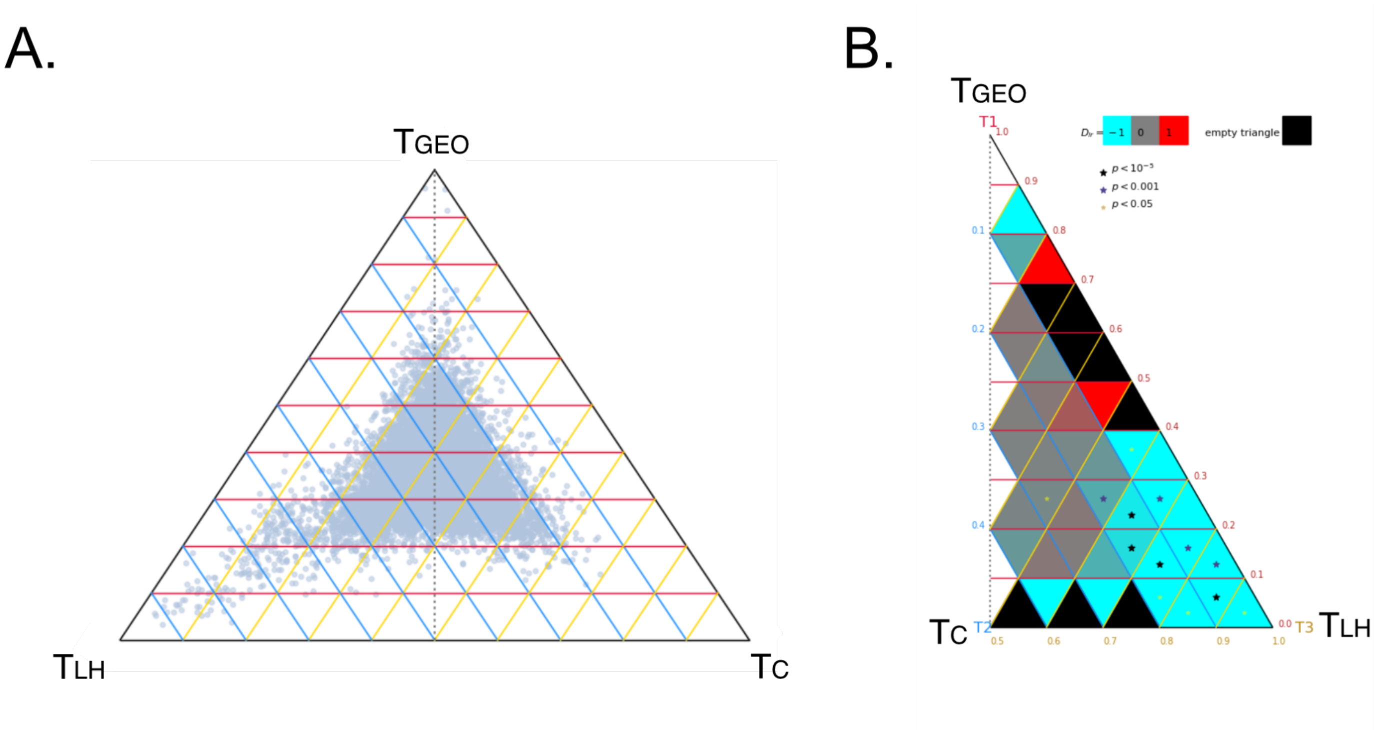


**Figure S10. TWISST'N'TERN results**. (**A**) Skew in topology weights toward the life-history topology ($T_{LH}$) in the ternary plot (each dot is a single window). (**B**) The asymmetry in topology weights is quantified using the $D_{LR}$ statistic and results plotted per sub-triangle (summarizing a range of topology weights). Asterisks indicate a significant asymmetry between corresponding left- and right- sided subtriangles, with negative $D_{LR}$ values indicating an enrichment of the left-sided subtriangles ($T_{LH}$).

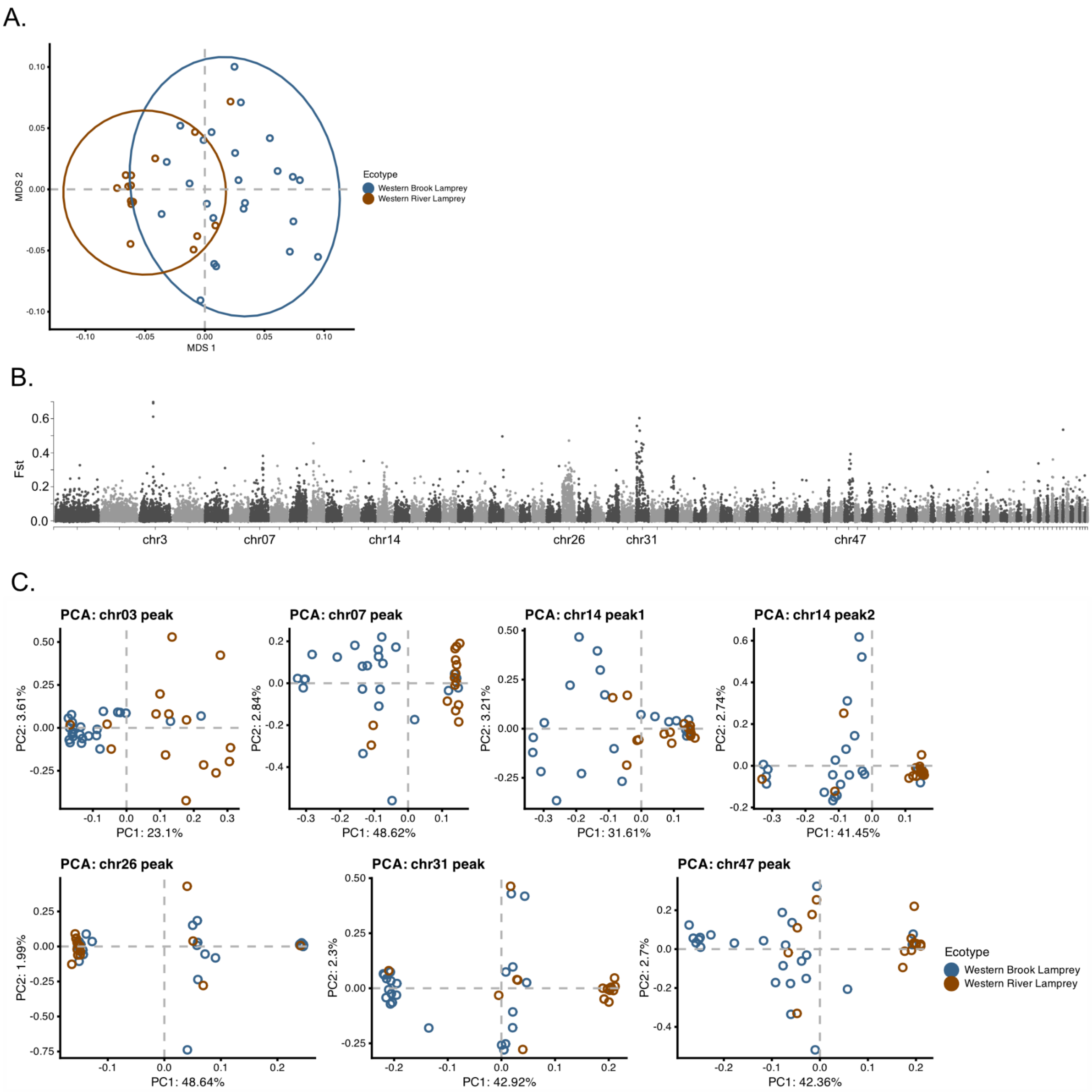


**Figure S11. (A)** MDS plot of Western brook and Western river lamprey ecotypes of *Occidentis ayresii* (formerly *Lampetra ayresii*) based on 102,000 SNPs. See legend for colours. (**B**) Manhattan plot of by-SNP Fst between Western brook and Western river lamprey ecotypes of *O. ayresii* across the genome. Divergence between ecotypes was relatively weak based on all identified SNPs (mean Fst = 0.016294), suggesting recent divergence and/or ongoing gene flow. Genome-wide Fst scans revealed several strong peaks of differentiation between ecotypes across the genome. However, none of these peaks overlapped with parasitism-associated regions in European *Lampetra*, suggesting that replicated ecotypes in *Occidentis* and *Lampetra* have evolved independently from each other using different genomic paths. (**C**) PCAs of SNPs within outlier peaks on six chromosomes. The PCAs for chr14_peak2, chr26 and chr31 show three distinct genetic clusters that could indicate the presence of recombination-suppressing chromosomal rearrangements.

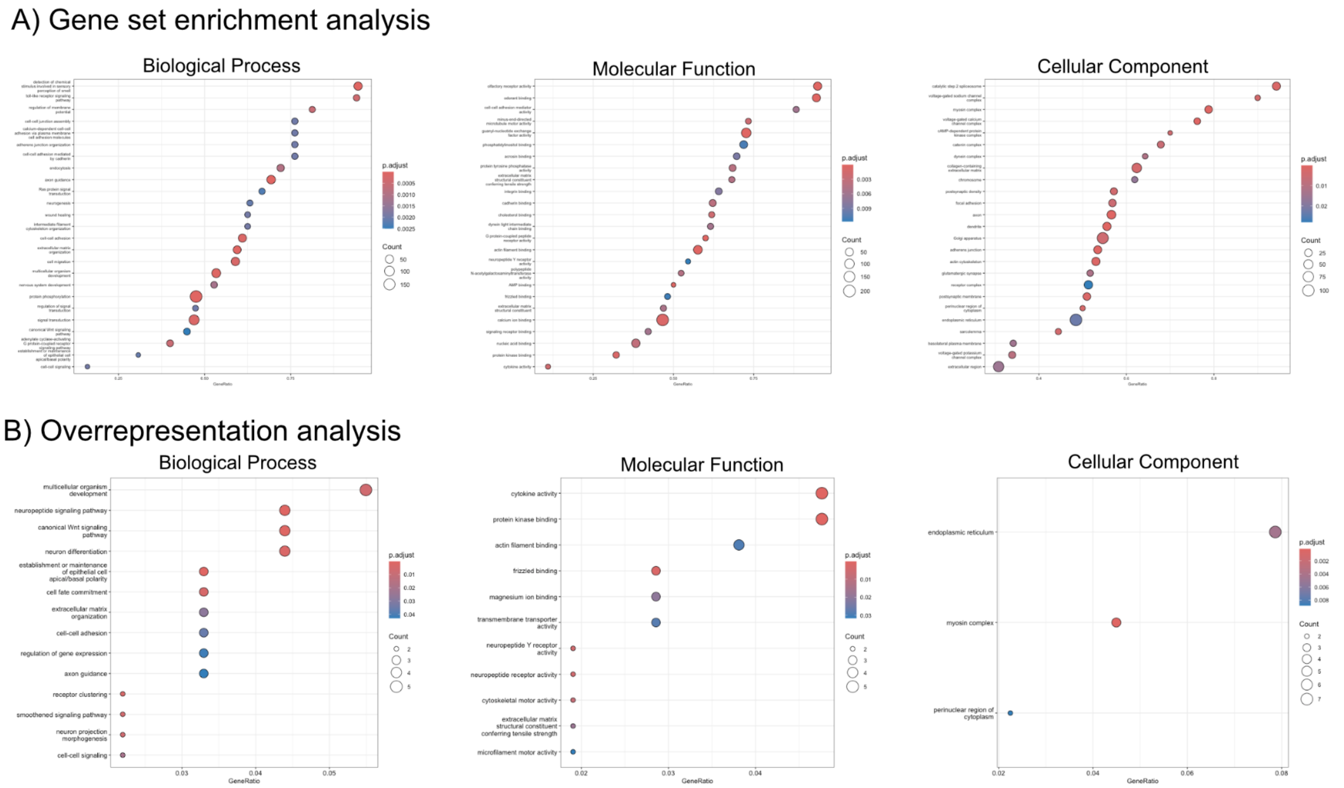


**Figure S12: Gene ontology enrichment.** Results of GO term enrichment analysis with *clusterProfiler*, showing the enrichment of GO terms associated with genes directly overlapping parasitism-QTL (table S2). (A) Results for the gene set enrichment analysis for GO terms separated by biological process, molecular function and cellular component. The size of the dot shows the number of significant genes and the colour the p-value (see Legend). The gene ratio shows the proportion of genes associated with a GO term that are present in our parasitism-QTL dataset. Shown are the top 25 terms. (B) Results for the overrepresentation analysis based on genes containing significant parasitism-QTL. Results are separated by biological process, molecular function and cellular component. All significant terms are shown. Full tables with results are shown in Supplementary file 1.

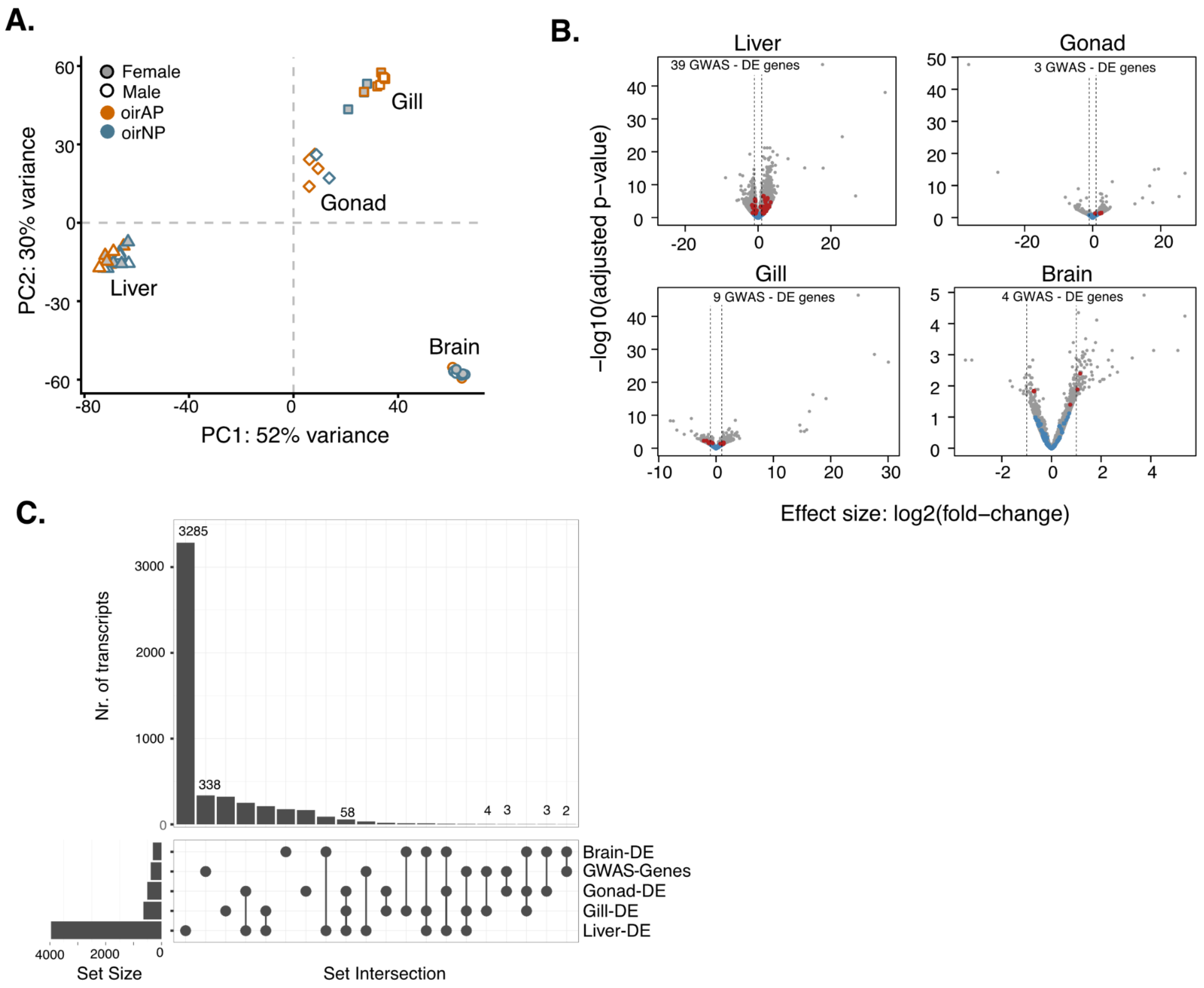


**Figure S13: Differential gene expression between anadromous-parasitic and non-parasitic lamprey across tissues.** (**A**) PCA based on vst-transformed expression of 12,605 genes across all tissues and individuals. Shape represents the tissue (triangle = Liver; diamond = Testis; Circle = Brain; Square = Gill), colour the ecotype (see Legend) and fill colour the sex (see Legend). (**B**) Volcanoplots showing the differential expression between oirAP and oirNP individuals by tissue. Genes that overlap GWAS SNPs are highlighted in red if they were differentially expressed (padj < 0.05) and blue if not (padj > 0.05). The number of differentially expressed (DE) genes under parasitism-QTL (GWAS) by tissue are given at the top of each volcano plot. (**C**) Overlap of differentially expressed transcripts with genes under parasitism-QTL.

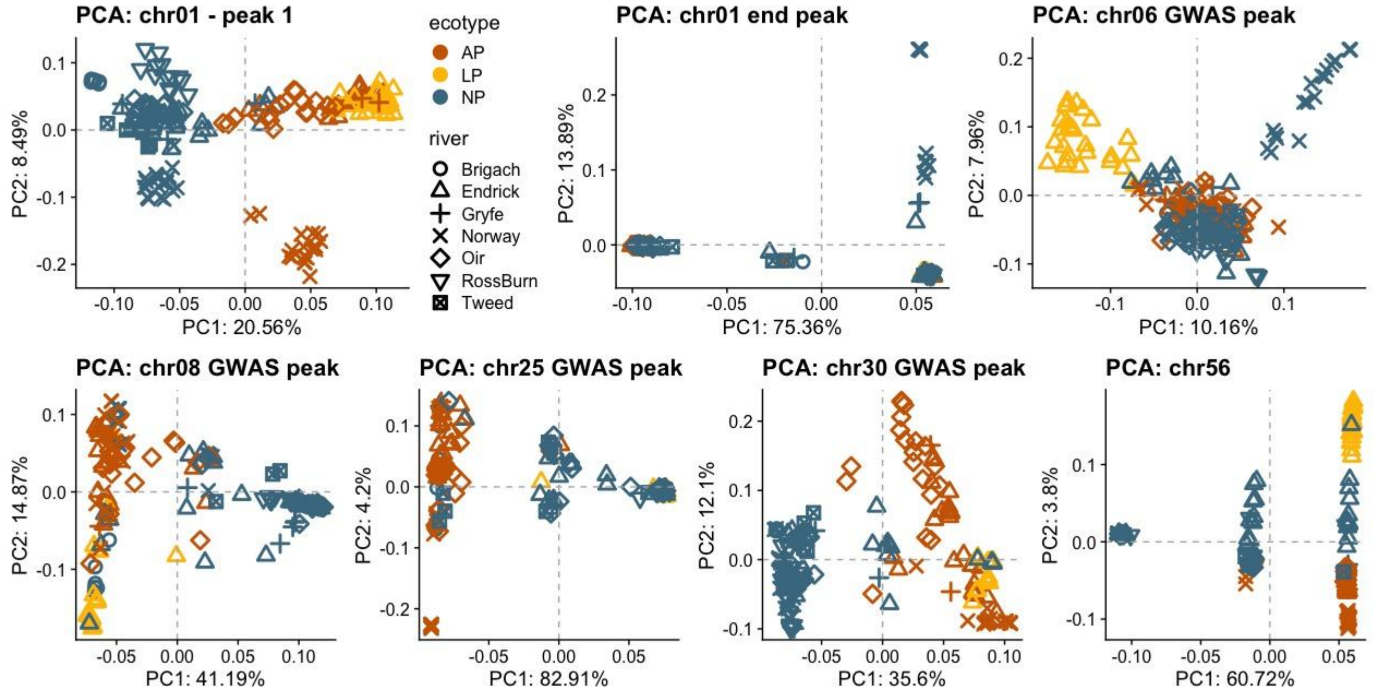


**Figure S14 -** PCAs of GWAS peaks across the genome. The PCA for chr1 - peak 1, includes all SNPs in the low-recombination region.

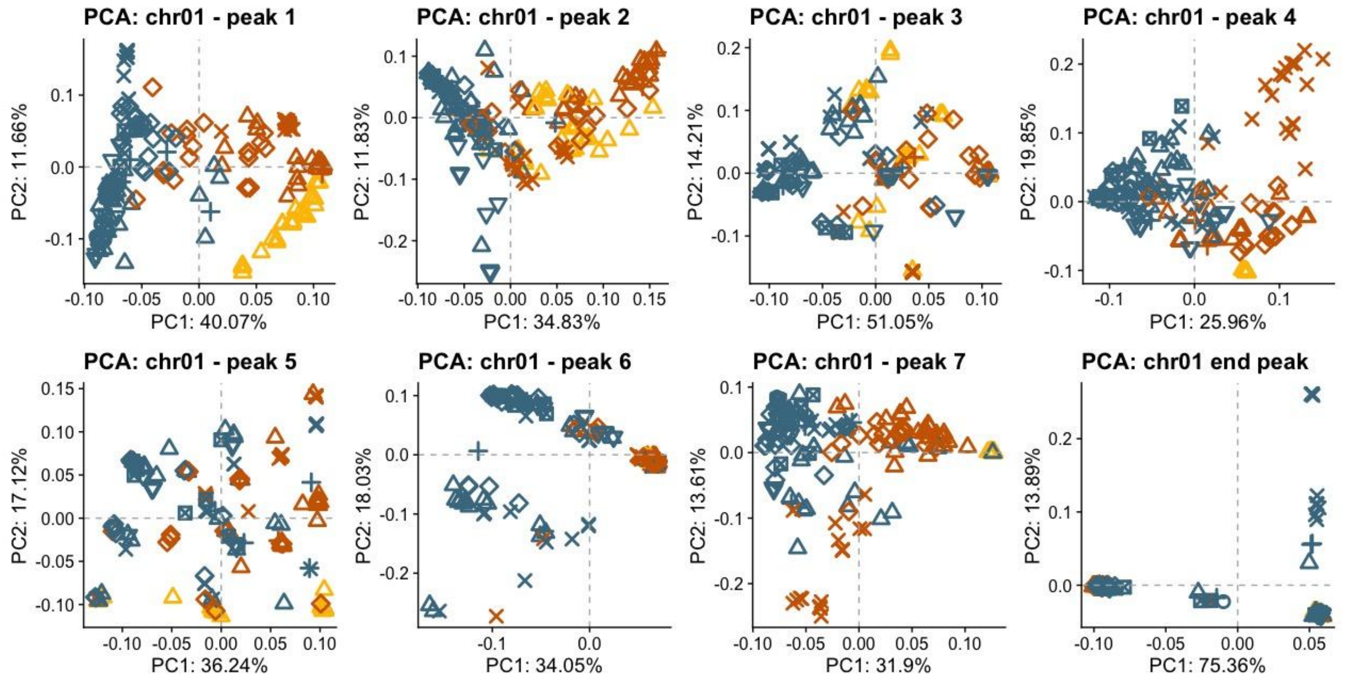


**Figure S15 -** PCAs for GWAS peaks on chromosome 1. The coordinates (in bp on chr1) for each peak are as follows: peak 1: 5,465,235-7,376,605; peak 2: 10,196,996-10,825,207; peak 3: 11,261,220-12,389,738; peak 4: 13,303,962-14,490,407; peak 5: 15,123,006-15,294,451; peak 6: 16,089,399-17,124,188; peak 7: 20,055,473-22,278,462; end peak: 41,092,658-41,117,697

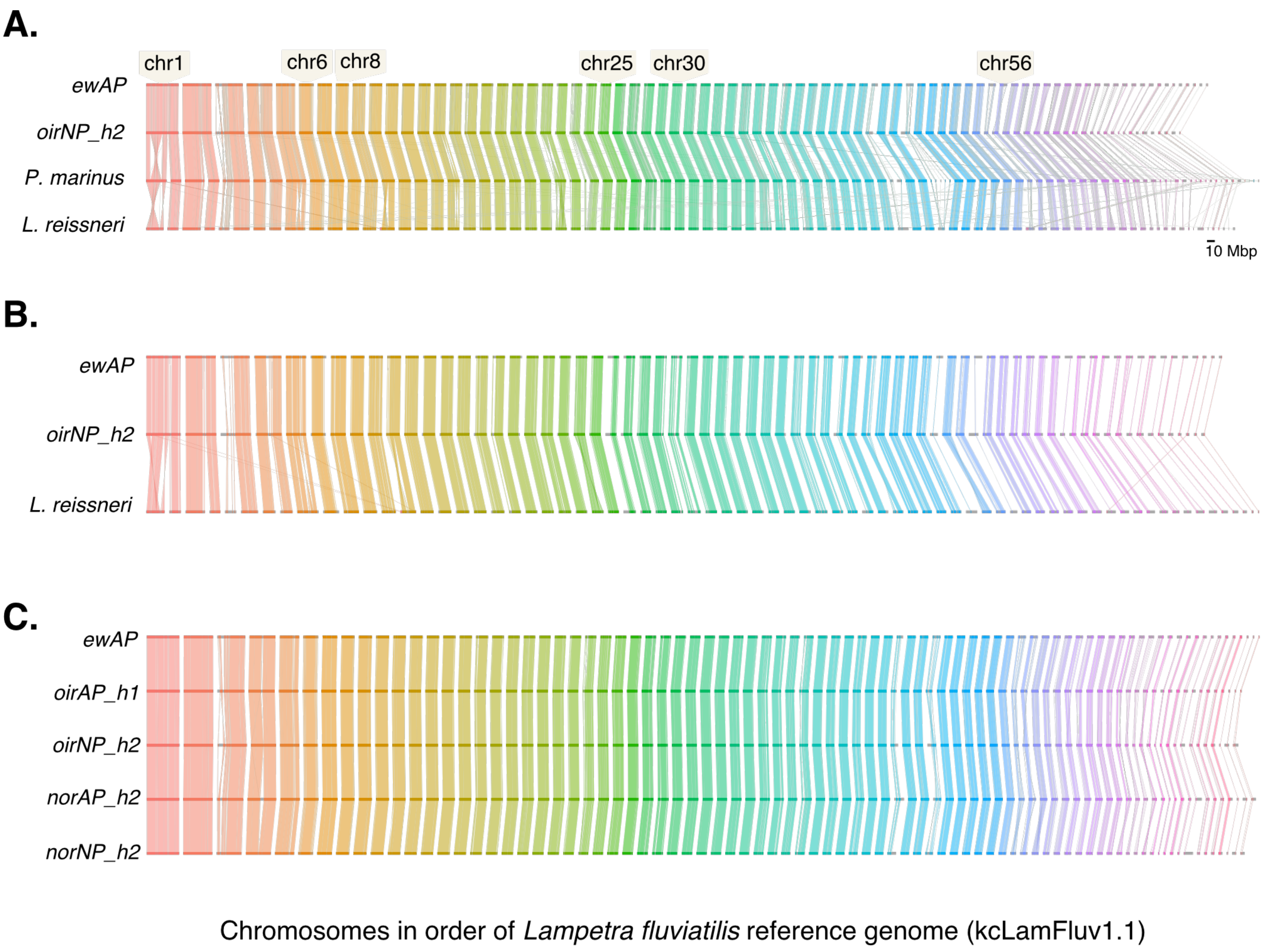


**Figure S16. Synteny across all chromosomes estimated using ntSynt.** (**A**) Synteny plots between ewAP (kcLamFluv1.1 primary assembly), oirNP-haplotype2, and both outgroup species. (**B**) Synteny plots between ewAP (kcLamFluv1.1 primary assembly), oirNP-haplotype2 the *L. reissneri* assembly. (**C**) Synteny plots between ewAP (kcLamFluv1.1 primary assembly), oirAP-haplotype1, oirNP-haplotype2, norAP-haplotype2 and norNP-haplotype2.

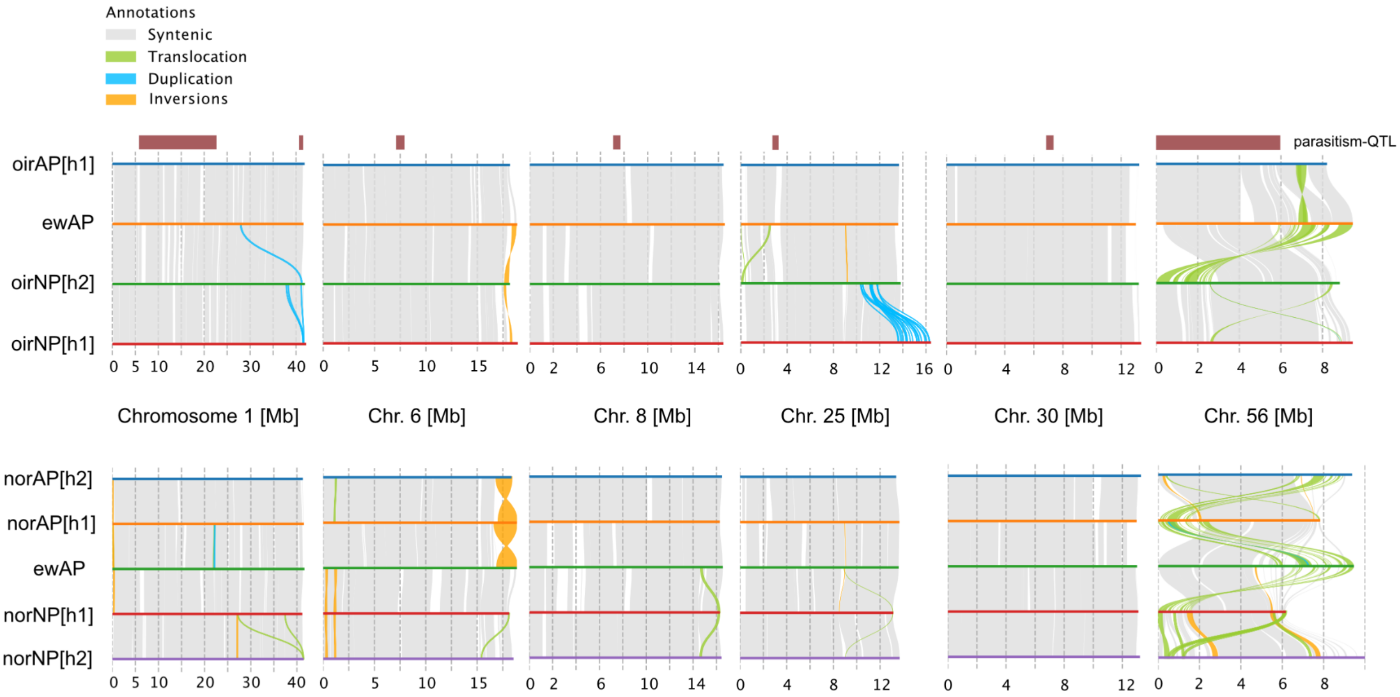


**Figure S17. Structural variation detection using SyRI.** (Top) Synteny plots for candidate chromosomes between genome assemblies from the Oir and Endrick produced using SyRI. Shown are haplotype 1 for the oirAP ecotype, and both haplotypes for the oirNP ecotype, compared to the reference assembly kcLamFluv1.1 (ewAP). Structural variant annotations are shown in the legend and the locations of broad parasitism-QTL regions are shown as red rectangles above the chromosomes. (Bottom) Synteny plots for the same chromosomes but comparing haplotype 1 and 2 for the parasitic (norAP) and non-parasitic (norNP) lamprey genomes from the Norwegian BioGenome Project. ewAP is shared between two plots and served as an anchor.

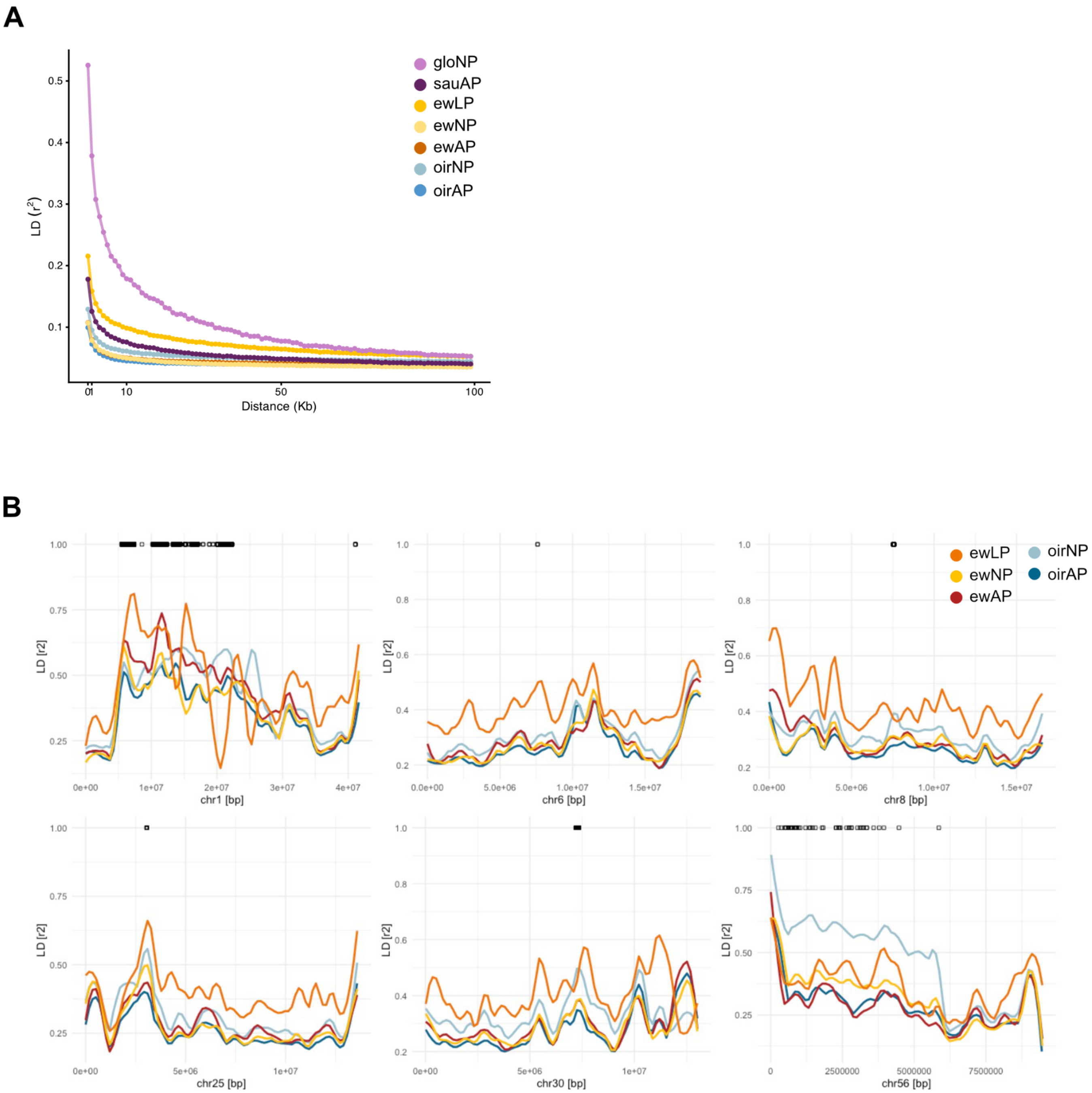


**Figure S18**. Linkage disequilibrium across the genome. (**A**) LD decay estimated for SNPs across chr2, chr5 and chr10 and plotted by population. See legend for colours. (**B**) LD in 10kb sliding windows across candidate chromosomes containing parasitism-QTL (shown in black at the top). LD values were loess smoothed using a span of 0.2 and coloured by population as shown in the legend.

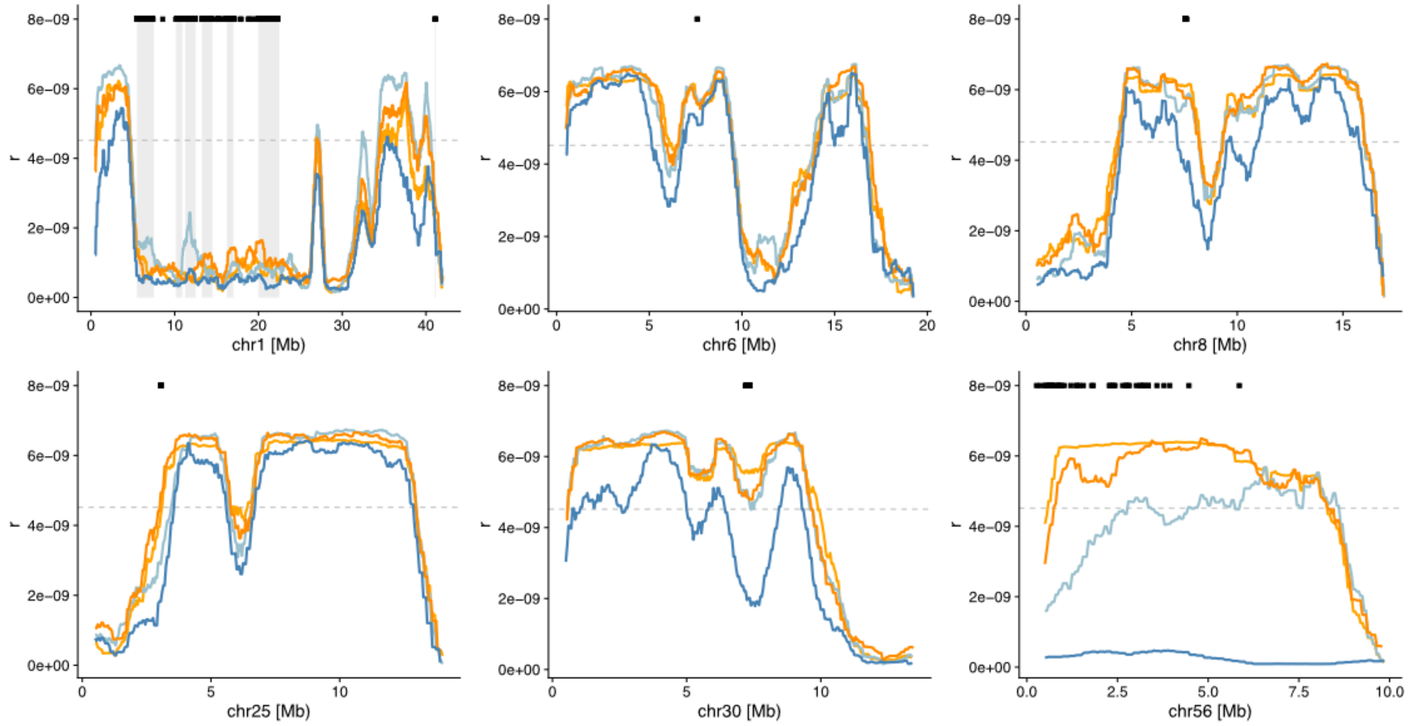


**Figure S19.** Population recombination rate [r] estimated using *RELERNN* for anadromous-parasitic and non-parasitic ecotypes in the Oir and Endrick. Recombination rates were plotted as mean recombination rates in 1Mb sliding windows with 50kb steps. Black dots highlight the locations of parasitism-associated SNPs and grey bars on chr1 show the location of distinct association peaks. Dark orange = oirAP, Orange = ewAP, dark blue = oirNP, light blue = ewNP.

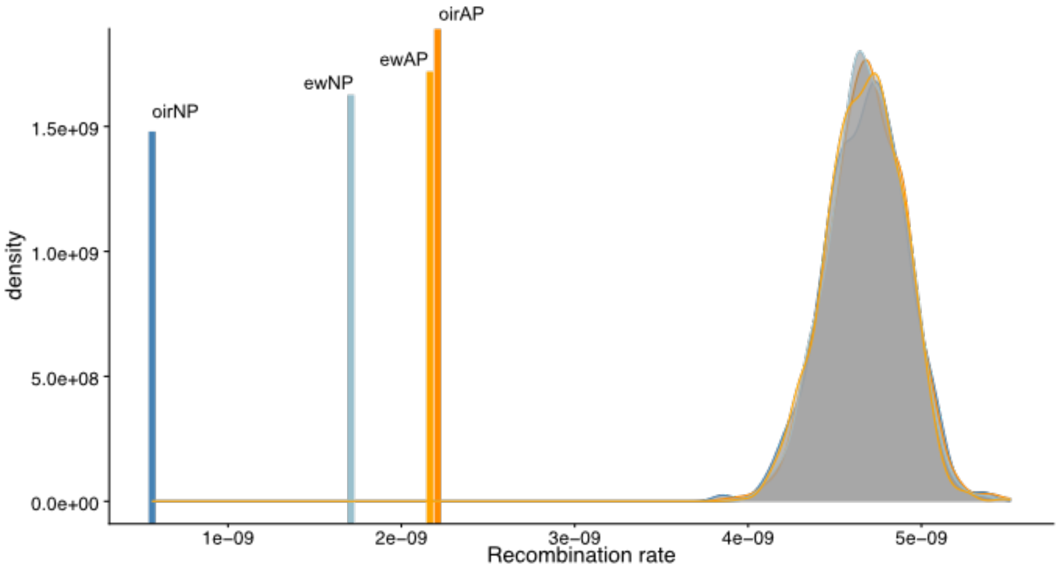


**Figure S20.** Differences in recombination rate [r] between GWAS peaks (lines) and the genomic background. Density distributions show the permuted background recombination rate across the genome by resampling x 1Mb windows with replacement 1000 times, with x being the number of windows overlapping significant parasitism-associated SNPs (n = 120). Background recombination rates (grey distributions) are very similar between ecotypes and populations, but recombination rates in parasitism-QTL differ more strongly, with the lower rates in non-parasitic (NP) compared to parasitic (AP) ecotypes.

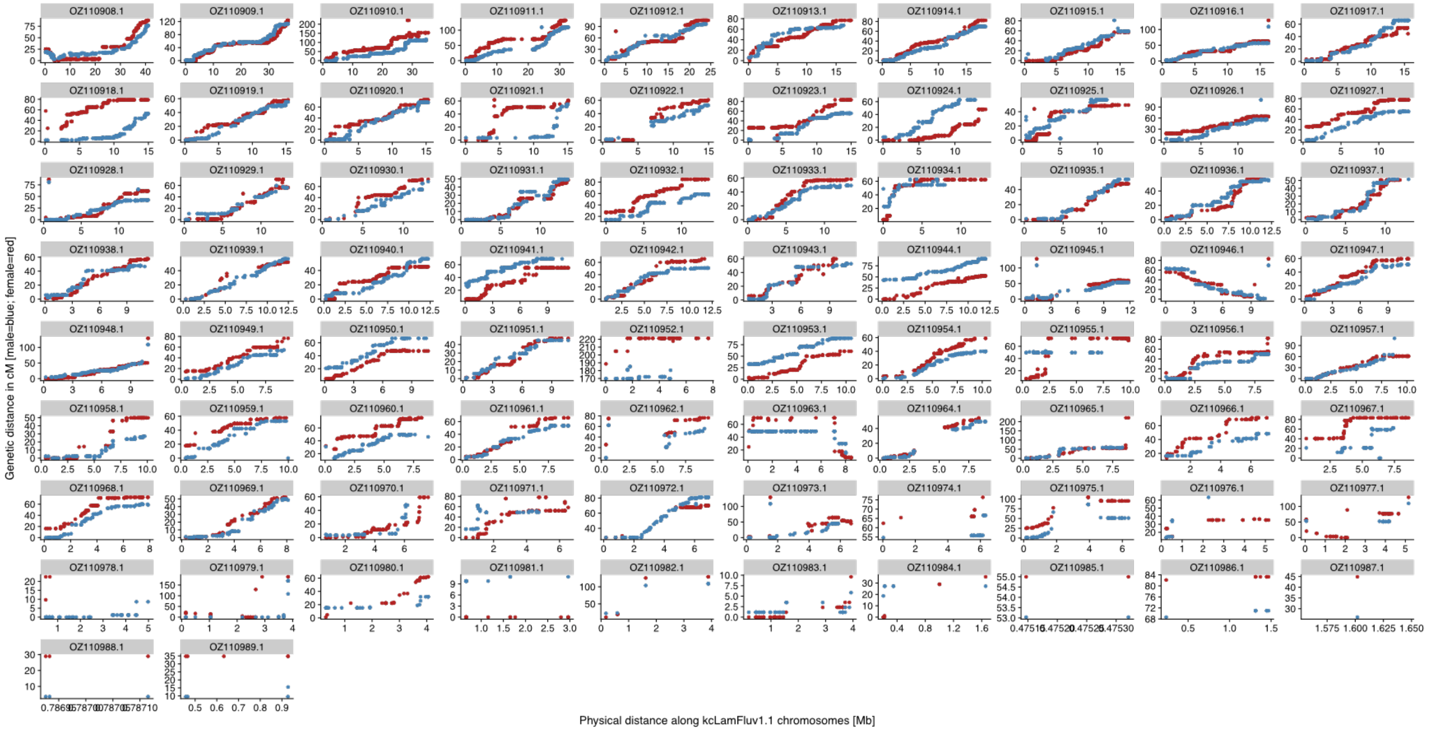


**Figure S21.** Male (blue) and female (red) linkage maps for Oir hybrid family, plotted along the kcLamFluv1.1 genome. Because the linkage map was created from an anadromous-parasitic female and a non-parasitic male, the male map represents the recombination landscape for non-parasitic individuals and the female map the recombination landscape for a parasitic individual.

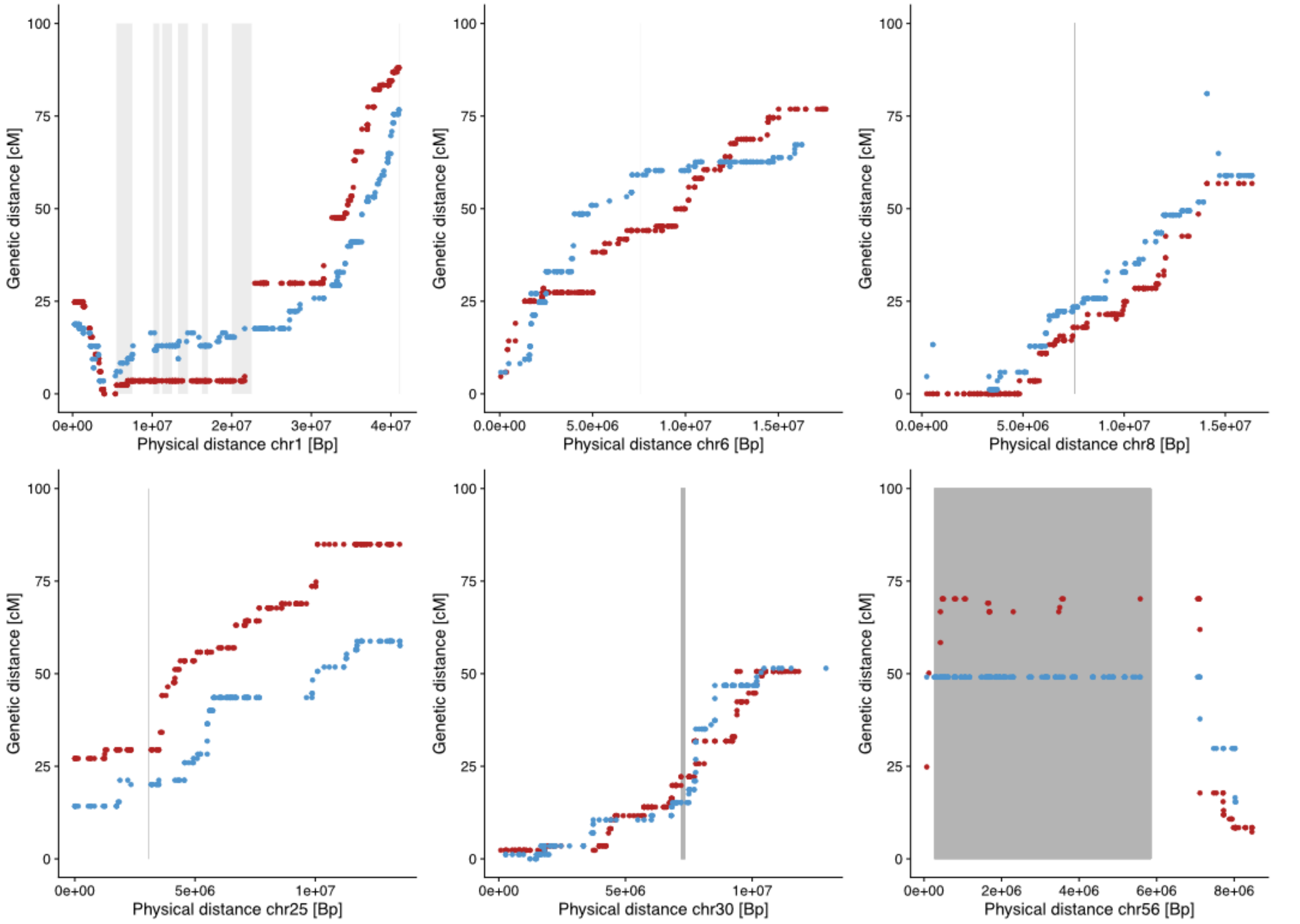


**Figure S22.** Male (blue) and female (red) linkage maps for Oir hybrid family, plotted for chromosomes with parasitism-QTL on kcLamFluv1.1 genome. The linkage map highlights the lack of recombination in the region containing parasitism-QTL (grey blocks) on chromosome1, particularly in the parasitic linkage map (male; blue) and the lack of recombination in the non-parasitic map (female, red) on chr56. Low recombination results in horizontal lines, as the genetic distance (cM) does not increase with physical distance (Mb).

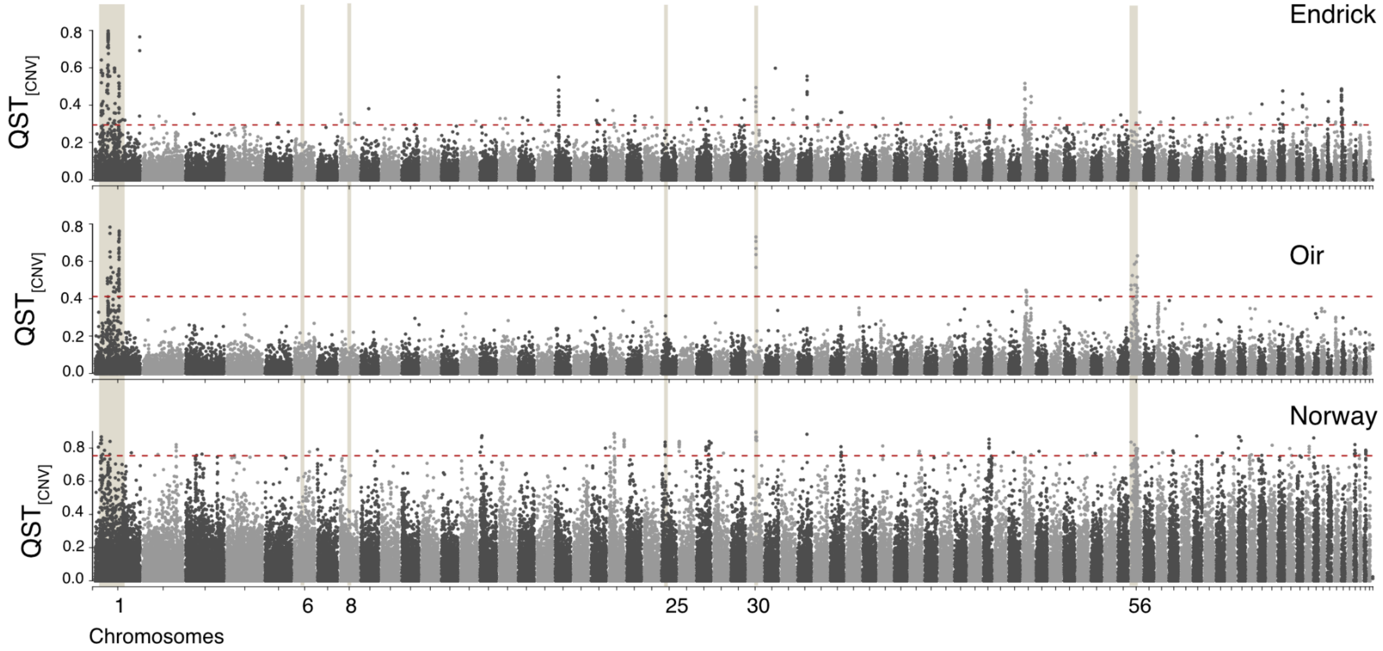


**Figure S23. Copy number variation between ecotypes.** QST for difference in normalised read depth between ecotypes. The dashed red line shows the top 1% Fst threshold, with QST values above that threshold considered to be under divergent selection. Beige fields show the location of parasitism-QTL identified using the GWAS between parasitic and non-parasitic lamprey.

(**Top**) $QST_{cnv}$ between sympatric anadromous-parasitic vs Non-parasitic ecotype from the Endrick, Scotland.

(**Middle**) $QST_{cnv}$ between sympatric anadromous-parasitic and non-parasitic ecotypes from the Oir, France.

(**Bottom**) $QST_{cnv}$ between allopatric anadromous-parasitic and non-parasitic ecotypes from Norwegian rivers.

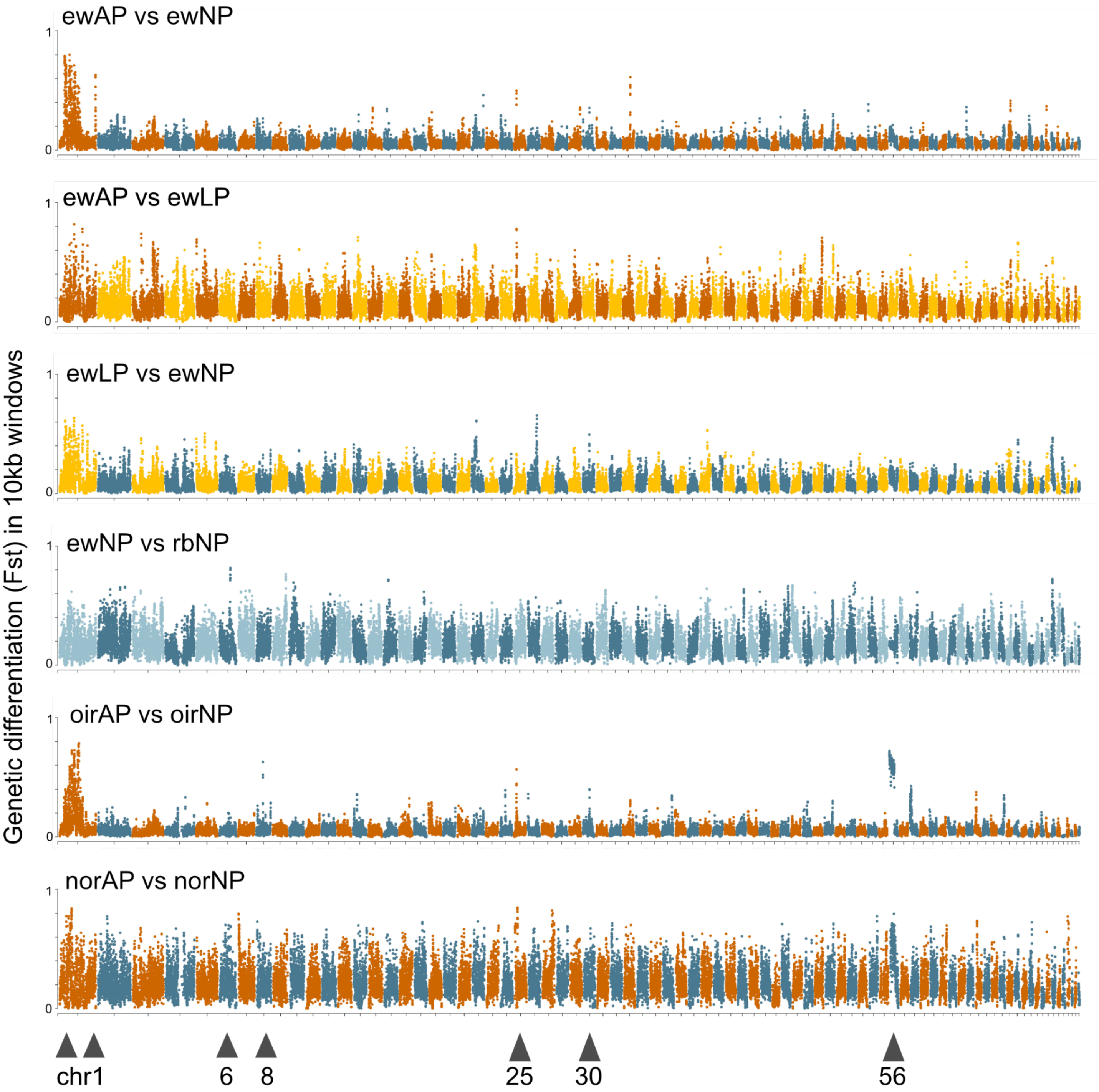


**Figure S24.** Manhattan plots showing Fst between sympatric ecotype comparisons in the Endrick, Ross Burn and Oir and between allopatric Norwegian populations. Fst is plotted in 10kb sliding windows (5kb steps). Comparisons are shown above. Chromosomes containing parasitism-QTL are marked with triangles at the bottom. Populations and ecotypes are labelled in the following way: ew = Endrick, rb = Ross Burn, oir = Oir, nor = Norwegian rivers; AP = anadromous-parasitic, NP = non-parasitic, LP = lake-parasitic.

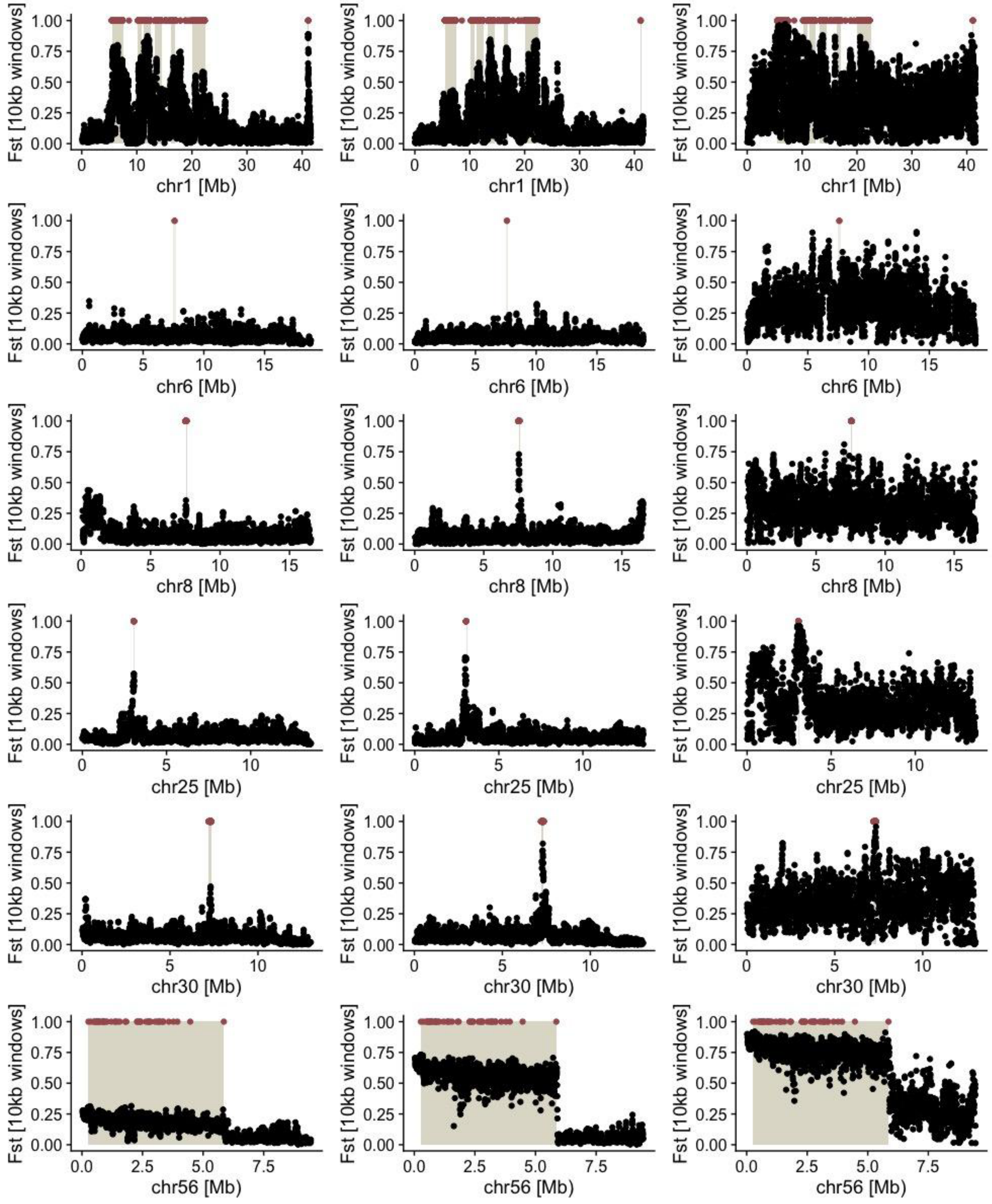


**Figure S25.** Plot Fst in 10kb sliding windows (5kb steps) between anadromous-parasitic and non-parasitic ecotypes along candidate regions. Each column is a population, from Endrick (left), Oir (middle) to Norwegian rivers (right), and each row is a candidate chromosome. Red dots show significant parasitism-QTL and beige rectangles show wider parasitism-associated regions.

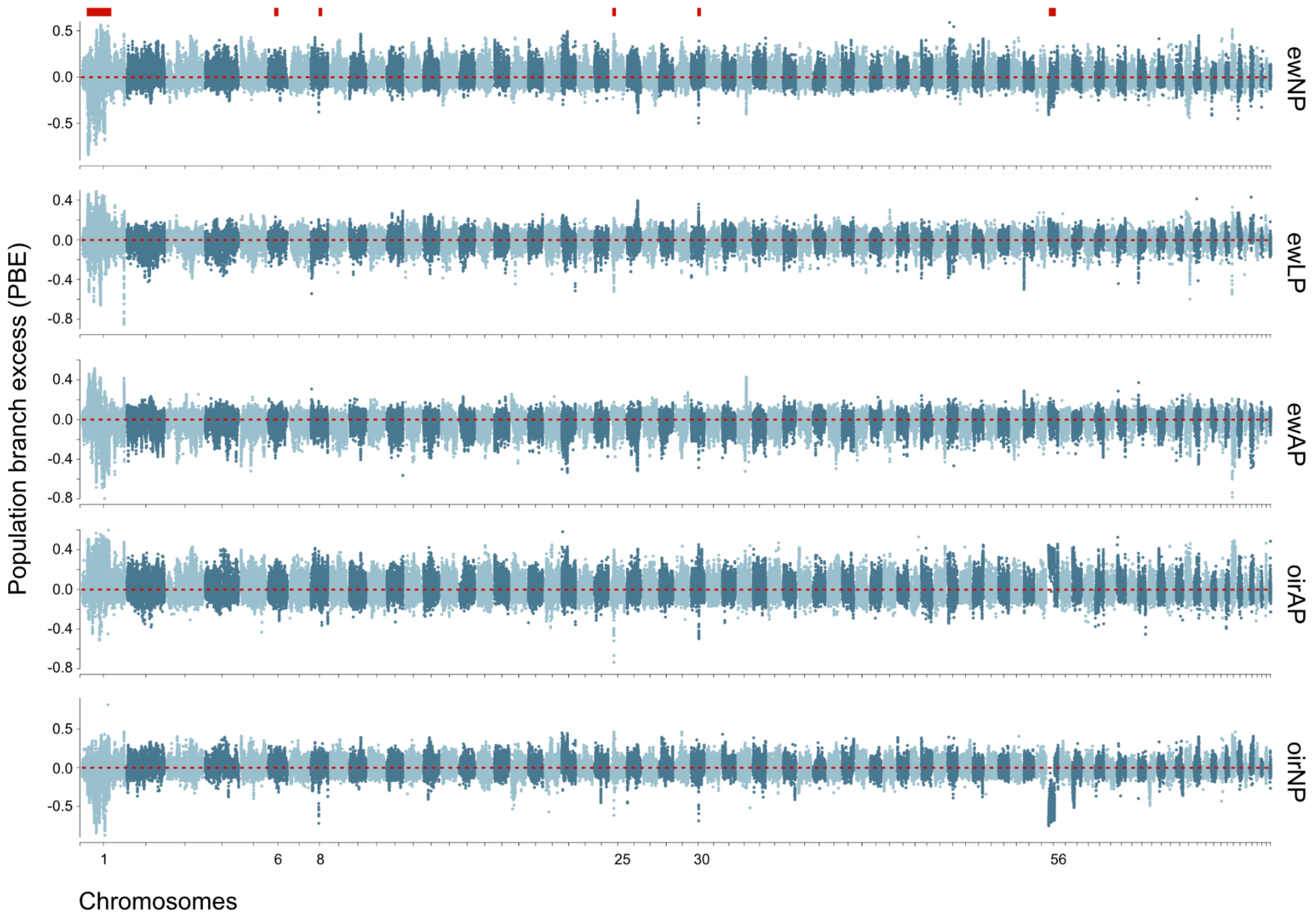


**Figure S26**. Population branch excess statistics in 10kb sliding windows across the genome for ecotypes from the Endrick (ewAP, ewLP, ewNP) and the Oir (oirAP and oirNP). Positive PBE values indicate that the branch length for this window is longer compared to expected branch lengths derived from comparisons of the other populations, which is indicative of positive selection. Negative selection suggests shorter than expected branch length, potentially through shared ancestry, gene flow, and/or relaxed/absent selection in the focal population with the other populations.

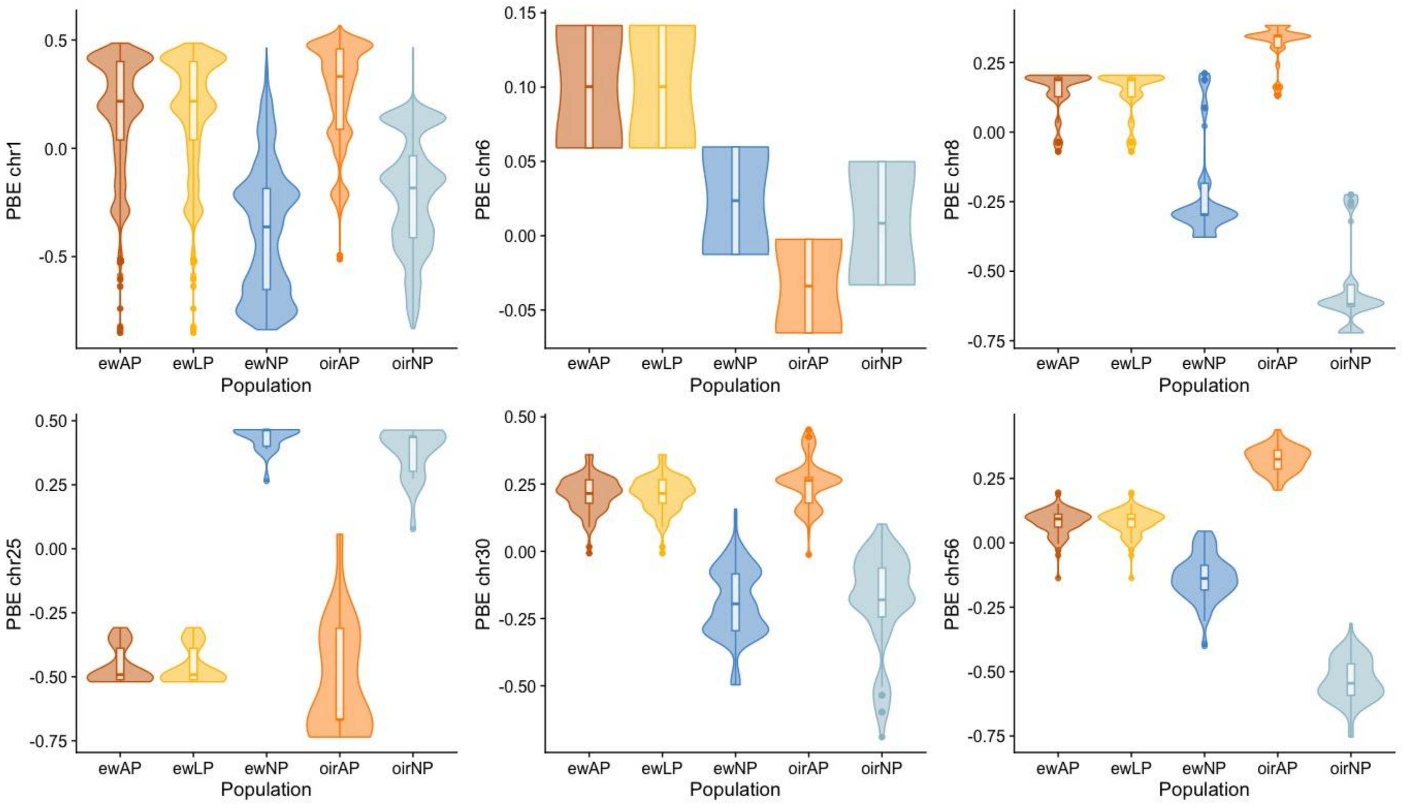


**Figure S27.** Population branch excess for genomic windows (10kb with 5kb steps) across parasitism-associated regions. PBE values were significantly higher in parasitic (ewAP, ewLP, oirAP) for GWAS associated SNPs on chr1, chr8, chr30 and chr56 , but significantly higher for non-parasitic ecotypes for chr25. There was no significant difference for chr6, but only four genomic windows overlapped significant SNPs on that chromosome. The pattern suggests that the difference in selection might not be consistent across rivers, with higher PBE values in parasitic ecotypes in the Endrick, and lower ones in parasitic individuals from the Oir. We compared PBE distributions in a pairwise fashion using Wilcoxon rank sum tests, with a p-value threshold of $p < 0.05$ for establishing significance.

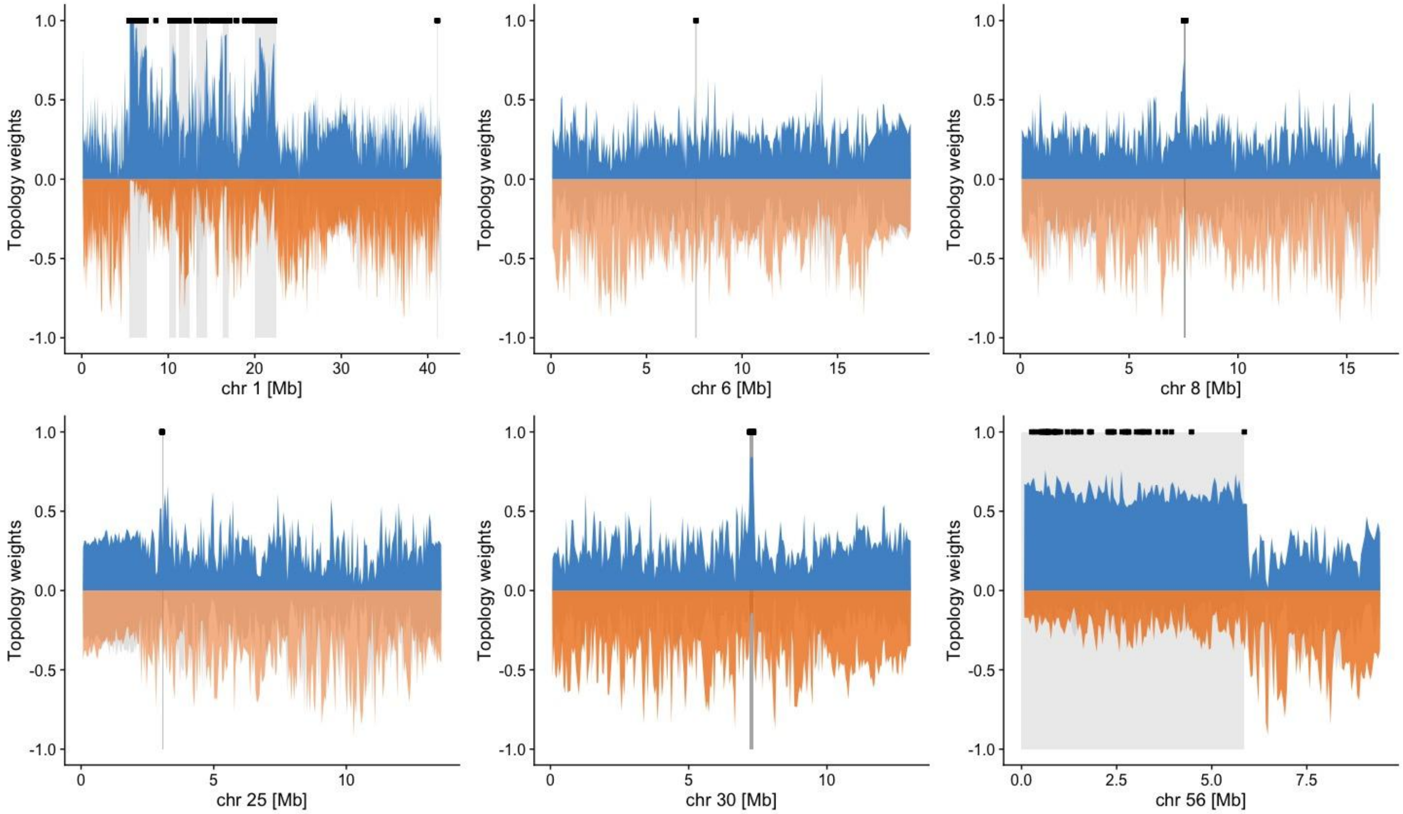


**Figure S28.** Genome scans of TWISST Topology weighting results for the three ecotypes in the Endrick (ewAP, ewNP, ewLP) and the adjacent Ross Burn. Results are shown for chromosomes with parasitism-QTL (shown in black and highlighted with grey blocks).

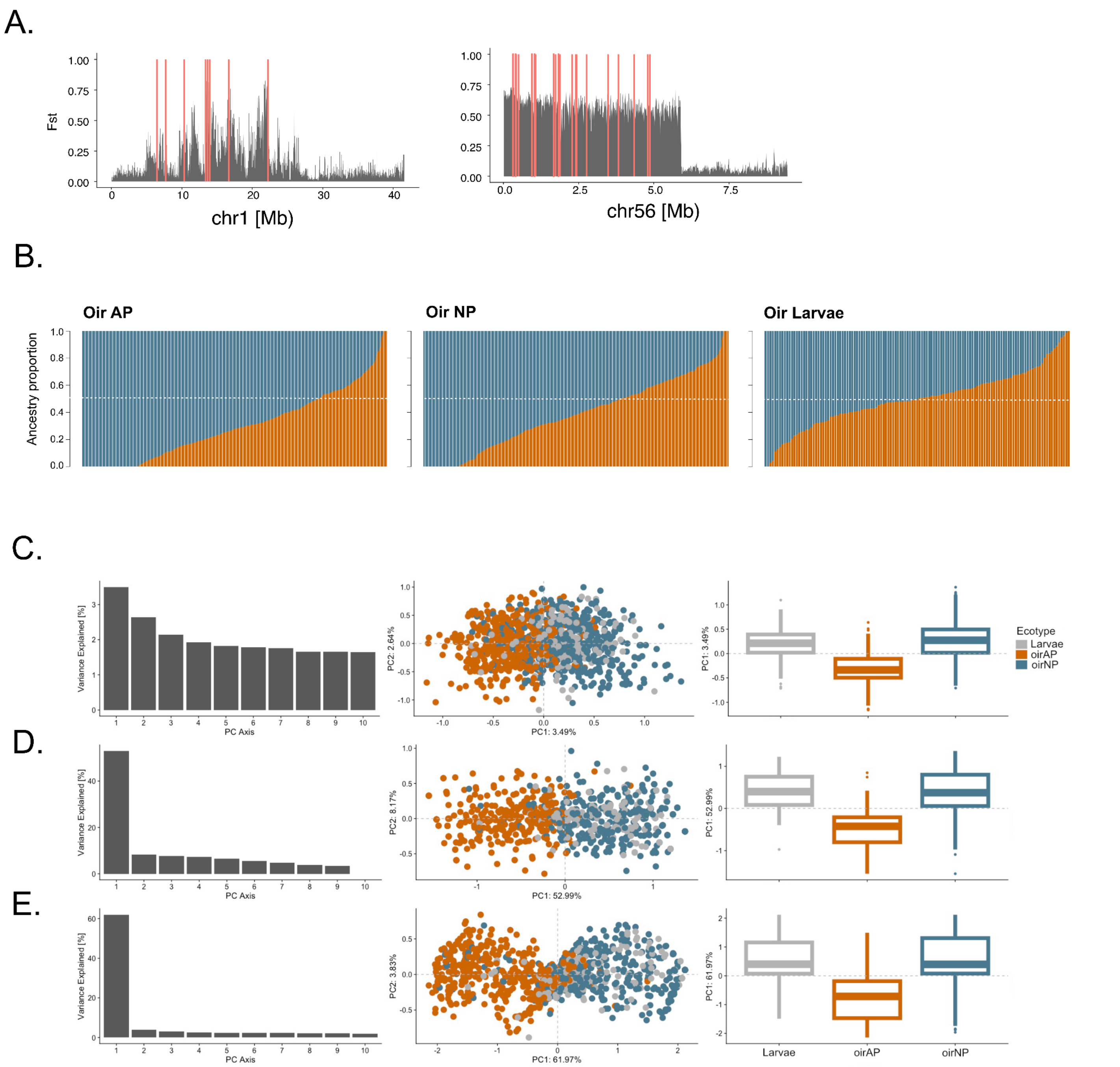


**Figure S29.** (**A**) Location of divergent SNPs on chr1 and chr56 in the HiPlex assay. The grey areas show the level of Fst between anadromous-parasitic and non-parasitic from the Oir based on whole-genome data. (**B**) Ancestry proportions based on neutral HiPlex SNPs for each ecotype and mixed larvae. Ancestry proportions were estimated using Admixture with a K of 2. (**C-E**) PCA based on HiPlex SNPs for non-divergent 'neutral SNPs (B), SNPs on chr1 (C) and SNPs on chr56 (D). Ecotypes differed significantly in genotype (summarised across SNPs as PC1) on chr1 ($t_{(941)}$ = 10.10, p<0.001), neutral SNPs ($t_{(941)}$ = -10.28, p<0.001), and in the frequency of *trans-inv56* ($t_{(941)}$ = -3.56, p<0.001), in line with whole genome data.

**A.**

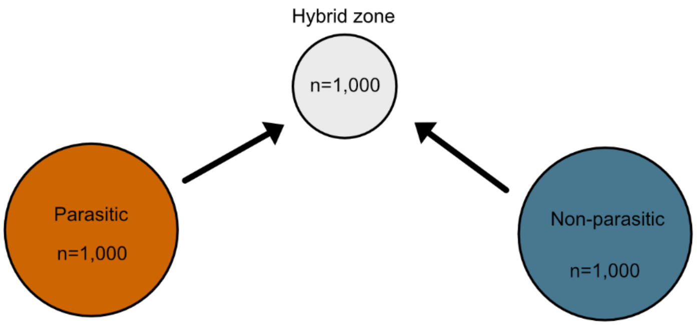


**B.**

| Fitness | Parasitic fecundity | % Random mating | Dispersal factor for parasitic indiv. | Dispersal of parasitic and non-parasitic indiv. |
|---|---|---|---|---|
| Similar | 3 | 10 | 1 | 0.001 |
| Hybrid breakdown | 5 | 30 | 5 | 0.005 |
| | | 50 | 15 | 0.01 |
| | | | | 0.05 |
| | | | | 0.1 |
| | | | | 0.3 |

**C.**

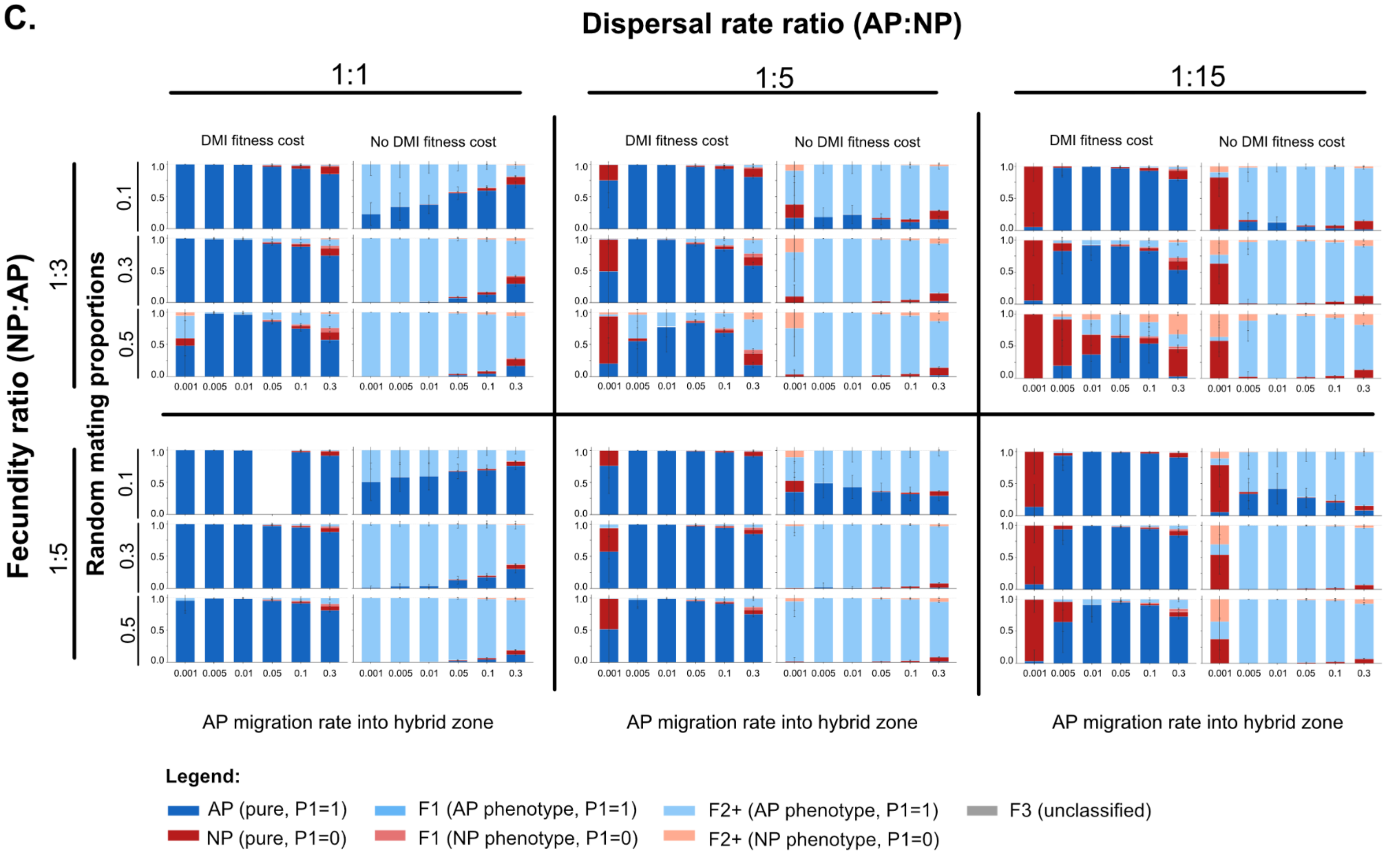


**Figure S30. NEMO hybrid zone simulations.** (**A**) Schematic of the NEMO simulation (the arrows indicate the dispersal from source populations to the hybrid zone). (**B**) Parameters used in simulations with the NEMO software. The transmission of the phenotype from parents to offspring results from the ancestry of the offspring (i.e. phenotype of the ecotype with the higher ancestry). The dispersal of NP is multiplied by 1 (same as AP), 5 or 15 to simulate a lower survival during the marine phase of AP. The fitness can be either equal among all categories of individuals or decreased in admixed individuals (i.e. hybrid individuals) due to Dobzhansky-Muller Incompatibilities (DMI). The fecundity of AP is multiplied by 3 or 5 compared to NP. Assortative mating within ecotypes can include either 10%, 30% or 50% of random matings (i.e. possibly between ecotypes). The dispersal rates vary from 0.1% to 30%. (**C**) Examples of the simulated hybrid proportions for a limited range of the parameters tested.

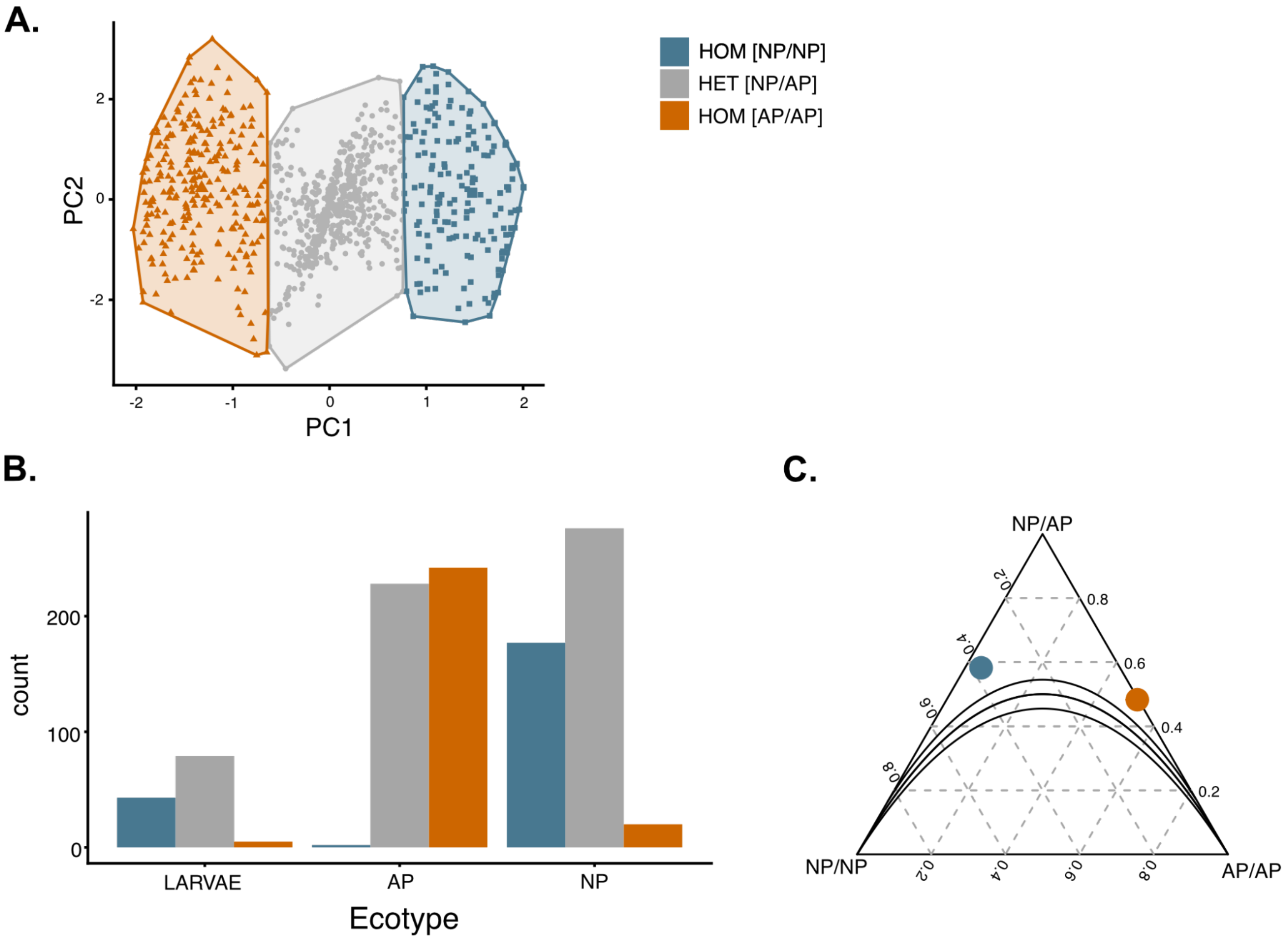


**Figure S31. HiPlex results for *trans-inv56*** (**A**) PCA for SNPs on chr56, with individuals labelled by inferred karyotype based on k-means clustering. (**B**) Count of the translocated inversion on chr56 (*trans-inv56*) by ecotype in adults and a random subset of larvae. (**C**) Deviation from Hardy-Weinberg Equilibrium for inversion frequencies in parasitic lamprey (red dot; $X^2$= 42, DF = 1, p-value = 9.09e-1) and non-parasitic lamprey (blue dot; $X^2$= 44.8, DF = 1, p-value = 2.16e-11) in the Oir. Expected frequency distributions under HWE are shown as curved lines.

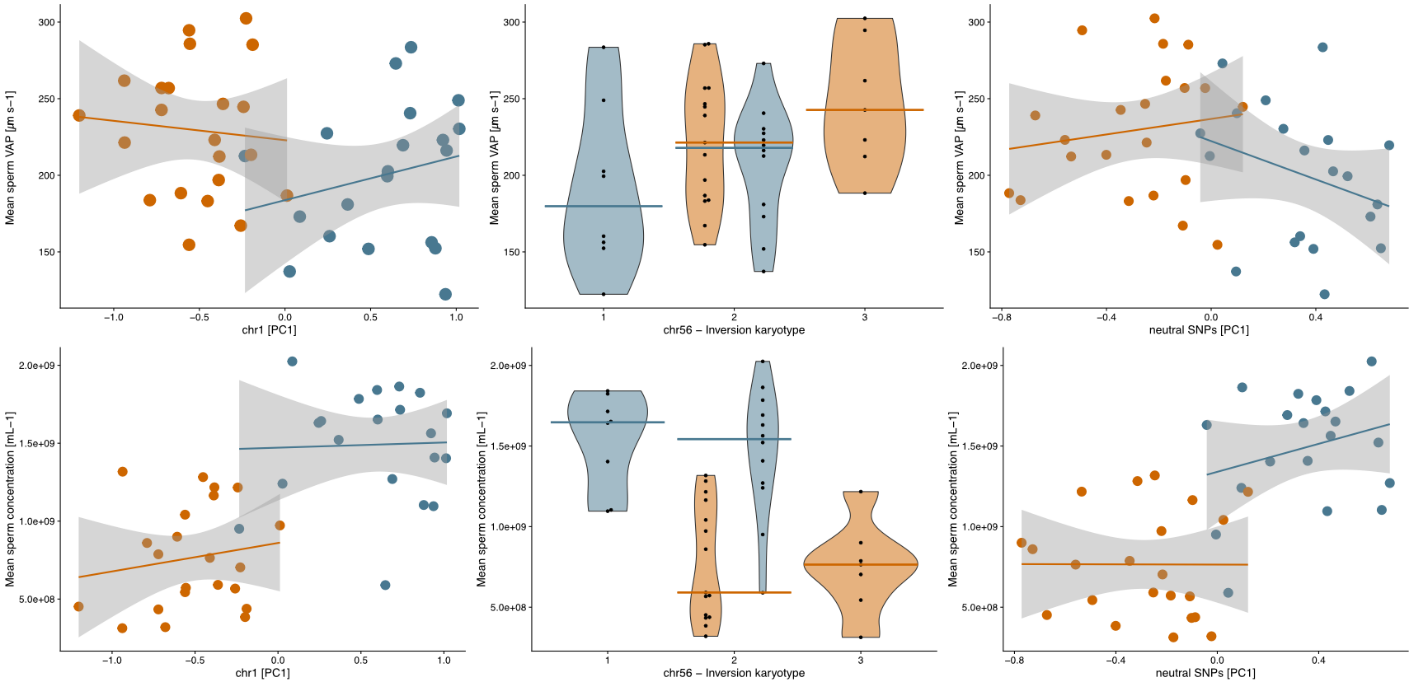


**Figure S32.** Correlation of genotypes at chr1 (genotype PC1), chr56 (karyotype) and neutral SNPs (genotype PC1) with mean sperm average path velocity (VAP) (**Top**) and mean sperm concentration (**Bottom**) in parasitic (orange) and non-parasitic (blue) lamprey from the Oir. Sperm concentration significantly differed between ecotypes (ANOVA: $F_{(1,40)} = 46.37$, $p<0.001$).
